# Modular DNA nanosensors enable state-aware multiplexed single-molecule mass photometry in complex media

**DOI:** 10.64898/2026.02.07.704534

**Authors:** Seham Helmi, Roi Asor, Maya Miller, Jan Christoph Thiele, Di Wu, Raman van Wee, Siyuan Song, Konstantin C. Zouboulis, Justin L. P. Benesch, Carol V. Robinson, Philipp Kukura

## Abstract

Proteins exist in diverse biochemical states, including oligomers, complexes, heterogeneous proteoforms and shed fragments that encode functional and regulatory information. Yet, these molecular states remain difficult to resolve with existing analytical techniques, which typically require labelling, immobilisation or amplification and collapse this information into a single readout. Here, we introduce a modular platform that integrates programmable DNA nanostructures with single-molecule mass photometry for rapid, label-free and multiplexed protein analysis in native and complex media, without washing, enrichment or immobilisation, making native biochemical states observable directly in serum and plasma. DNA nanostructures act as nanosensors whose mass and mobility on supported lipid bilayers provide orthogonal identifiers for target identity and biochemical state, thereby decoupling recognition from readout. We define the analytical specificity and response window, demonstrate quantitative affinity determination, and resolve oligomeric and proteoform differences under native conditions. The nanosensors are rapidly reprogrammable to new targets, support multiplexed detection with internal controls for non-specific interactions, and enable selective resetting via strand displacement. Together, these capabilities establish nanoscale programmability as a route to state-resolved single-molecule protein profiling adaptable to both diagnostic and mechanistic applications.

## Introduction

Proteins govern essential biological processes, regulate cellular signalling networks, and serve as central biomarkers for health and disease. Accurate detection and characterisation of proteins are therefore foundational to progress in proteomics, molecular biology, and diagnostics. Yet, unlike nucleic acids, proteins lack intrinsic amplification mechanisms, making them fundamentally more difficult to detect with high sensitivity and specificity. Detection becomes particularly challenging in complex biological environments where abundance, modification and oligomeric state, and interaction context vary widely^1^.

Most protein-detection technologies couple molecular recognition to signal generation, constraining modularity and limiting the information content. Enzyme-linked immunosorbent assay (ELISA) and fluorescence-based assays^2–6^ are the workhorses of protein detection, offering sensitivity and scalability but rely on labelling, multi-step washing, and indirect readout of molecular identity. As a sandwich-type immunoassay, ELISA requires two antibodies to bind simultaneously to distinct epitopes on the target and to remain tightly associated through repeated washing steps, thereby restricting detectable interactions to those with slow dissociation kinetics^2^. This dependence on high-affinity antibody pairs limits applicability to well-characterised targets and complicates adaptation to emerging biomarkers^2^.

Surface-based sensors, such as surface plasmon resonance (SPR)^4,7^, provide kinetic information but are restricted by immobilisation geometry, surface crowding, and limited multiplexing. Consequently, existing methods typically report only a single physical parameter—optical intensity or resonance shift—without revealing stoichiometry, oligomeric state, or molecular identity. Yet, many disease-associated proteins exist in multiple oligomeric or conformational states that govern their biological activity and clinical relevance^8,9^, remaining indistinguishable in ensemble-averaged measurements. Additionally, diagnostic accuracy frequently improves when multiple biomarkers are measured together rather than individually^10–13^, underscoring the need for assays that support intrinsic multiplexing.

These limitations have motivated the development of next-generation platforms that decouple recognition from signal generation while enabling programmable, high-resolution readouts. Fluorescence-based strategies^14–18^ or conformational switching^14,18,19^ have expanded the dynamic range of optical sensors, while DNA nanostructures^14,17–26^ have introduced a programmable design space for spatially controlling ligand presentation, enabling logic-gated or multiplexed recognition. Parallel to these advances, mass photometry (MP)^27–32^ has emerged as a label-free technique for detecting and quantifying single proteins in solution. MP directly reports on molecular mass with minimal sample processing and resolves binding stoichiometry, oligomerisation and complex formation without labelling or immobilisation.

Despite these developments, a generalisable platform that combines programmable recognition, real-time analysis, and multiplexed state resolution in complex biological media remains lacking. Most current MP implementations are restricted to purified systems^28–30,32^, with target identity inferred from expected mass values alone. There is no established framework for orthogonal encoding of recognition specificity or for resolving multiple targets simultaneously in heterogeneous samples such as plasma or cell lysate.

Here, we introduce a modular platform that integrates programmable DNA-origami nanostructures^19–21^ with MP to enable real-time, multiplexed and state-aware protein analysis directly in complex media. The platform reports target binding directly through measurable mass shifts rather than through indirect signal amplification. DNA nanostructures serve as spatially addressable nanosensors for capture ligands while diffusing on supported lipid bilayers (SLBs). This combination enables correlated measurement of molecular mass and diffusivity changes upon target binding. The SLB provides a passivating surface that suppresses nonspecific adsorption and allows analysis directly in complex media without the need for disruptive washing steps. Together, these features decouple molecular recognition from signal generation and allow discrimination between specific and nonspecific interactions. Performed under native, solution-based conditions, the platform delivers real-time, single-molecule mass readouts with rapid reprogrammability and broad adaptability across biomarker panels.

### Engineering a DNA-based single-molecule capture complex

Our detection platform consists of individual DNA-origami nanostructures bound to SLBs formed on a microscope cover glass (Fig. 1a). Each origami, folded from a long scaffold and short staple strands, provides a programmable nanostructure carrying capture strands that hybridize to DNA-conjugated protein-recognition elements and anchor strands that hybridise to cholesterol-modified oligonucleotides for SLB attachment. The origami design consists of two stacked layers of parallel helices arranged on a square lattice and was folded from a 1,496-nucleotide custom scaffold (p1496), generated via double restriction digestion^33^ of the p8064 scaffold using BsaBI and XmnI enzymes (Fig. 1b). Defined double-stranded termini were produced by hybridising short complementary oligonucleotides to the digestion flanking regions (Fig.S1, SI Methods 1-3), which dissociate upon mild heating to enable efficient scaffold isolation and subsequent folding. SLBs were chosen for their exceptional passivating properties^34,35^ and for enabling specific readout through correlated analysis of mass and diffusivity.

**Figure 1.**
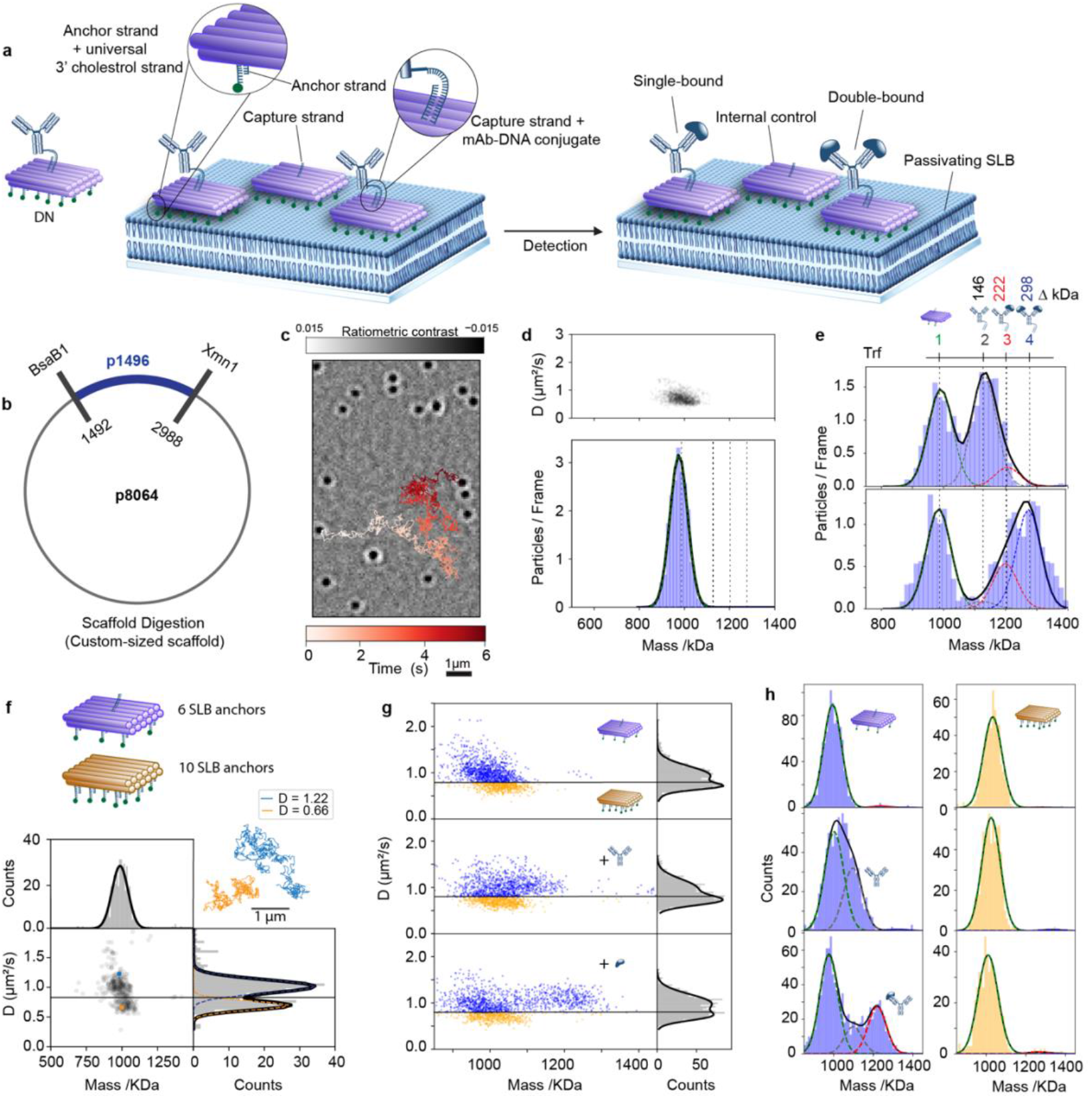
Engineering a membrane-tethered DNA origami nanosensors for single-molecule protein detection via MP. **a**, Schematic of the DNA-nanosensor (DN) platform. **b**, MP image of 660 kDa origami nanostructures diffusing on a SLB with highlighted 6s single trajectory trace. **c**, Measured mass and mobility distributions from 1670 trajectories extracted from two 60 s recordings. **d**, Mass distribution for Trf-capture DNs in the absence (top) and presence (bottom) of Trf. **f**, Schematic of two nanosensor variants bearing different numbers of cholesterol–linked anchors, with two representative single-particle trajectories illustrating their distinct diffusivities on the SLB. **g**, Mass–diffusion scatter plots showing two diffusion-separated populations before (top) and after incubation with anti-Trf mAb (middle) and addition of 100 pM recombinant Trf (bottom), with ROIs indicated by black horizontal lines set at the intersection of the fitted diffusion Gaussians. **h**, Corresponding mass distributions for the two diffusion-resolved populations before (top), after anti-Trf mAb (middle), and after recombinant Trf (bottom) corresponding to the scatter plot in **g**.

Individual structures on SLBs appear as mobile, diffraction-limited features^28,29^ in MP (Fig. 1b), with an optical contrast proportional to the origami mass (990 ± 40 kDa) and mobility (0.8 ± 0.3 µm^2^/s) set by the number of cholesterol-anchors (Fig. 1c). Upon addition of a complementary ssDNA-antibody conjugate (anti-transferrin mAb), a second population appears shifted by 146 ± 1 kDa (1136 kDa), confirming specific hybridization of a single mAb to the capture strand (Fig. 1d, peak 2) and forming the detection population, hereafter referred to as DNA-nanosensors (DNs, movie S1). Importantly, the native unbound origami peak at 990 kDa (Fig. 1d, peak 1) remains visible, conveniently serving as an internal control for nonspecific binding and as a mass reference.

Addition of purified antigen (transferrin (Trf), 76±1.5 kDa) leaves the control origami population unchanged, while shifting the DNs to higher mass, even at very low analyte concentrations (400 pM, movie S2). The new population is well-described by a mixture of Gaussian distributions consisting of the control (990 kDa), single-bound (1212 kDa) and double-bound DNs (1288 kDa) (Fig. 1d, peaks 3 and 4, respectively). The observed mass shift provides a direct readout of Trf binding, while the unchanged internal reference peak (990 kDa), together with the correlated redistribution of the DN population into single- and double-protein-bound species (1212 and 1288 kDa), distinguishes specific target binding from nonspecific interactions. Due to the simplicity of the interaction, even unresolved spectra can be easily deconvoluted to provide relative abundances of free, 1-bound and 2-bound DN sensors.

To explore how nanosensor mobility can be tuned within the same measurement environment, we prepared two variants of the p1496-based origami bearing different numbers of cholesterol anchors, resulting in indistinguishable masses but well-separated lateral diffusivities on the SLB (Fig. 1f). When combined, the two populations remained resolvable solely by diffusion, enabling the definition of mobility-based regions of interest (ROIs), separated by black horizontal lines. This establishes diffusion as an orthogonal physical encoding mode without altering nanosensor geometry or target affinity.

Functionalising only the faster-diffusing population with capture strands while leaving the slower-diffusing population unmodified enabled assessment of target binding in the presence of a non-binding population. Upon incubation with anti-transferrin mAB followed by recombinant Trf, only the functionalised nanosensors exhibited mass increases, while the unmodified population remained unchanged (Fig. 1g, h). These results demonstrate that diffusion-resolved encoding can be engineered and maintained under binding conditions within the same measurement.

### Modular, stoichiometry-resolved, and quantitative affinity measurements

To evaluate the quantitative performance and modularity of our platform, we first focused on Trf as a model system. We established solution-phase reference signatures using standard MP landing assays in which Trf and its DNA–mAb conjugate were mixed at 1.5:1 protein:mAb molar ratios, yielding four discrete species corresponding to free protein, unbound conjugate, and one- and two-bound complexes (Fig. 2a, top and Fig. S2a). These stoichiometrically resolved references served as mass benchmarks for downstream quantitative analysis on SLBs.

**Figure 2.**
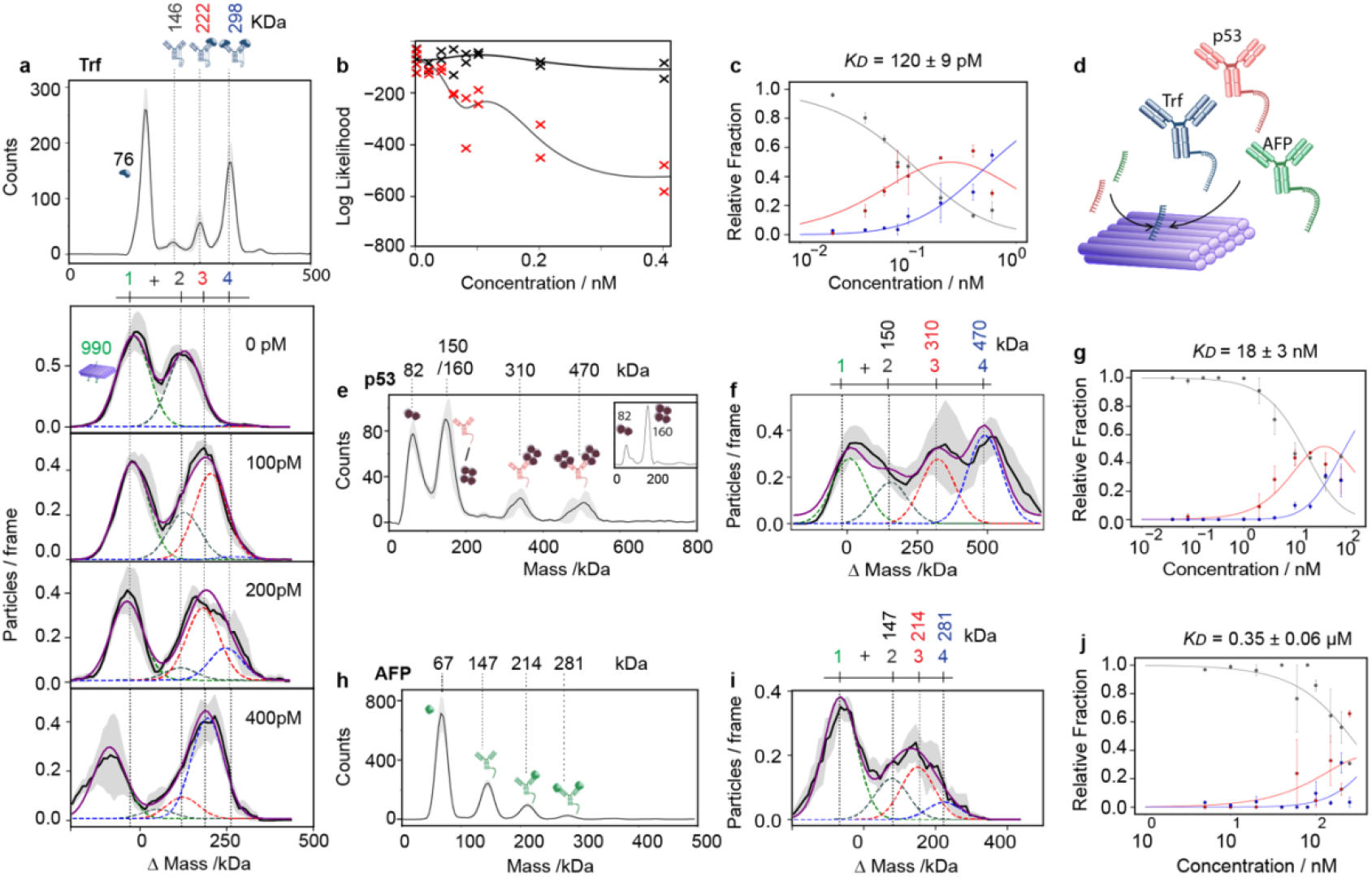
Quantitative and modular binding analysis of DNA nanosensors. **a**, Solution landing assay resolving free Trf, unbound DNA–mAb conjugate, and one- and two-bound complexes (top) and membrane-tethered nanosensor titration (bottom) showing discrete occupancy states, across three technical repeats at 1.5:1 Trf:Trf-mAb. **b**, Fractional occupancies extracted by constrained global Gaussian fitting across the Trf titration series and interpreted using a Poisson-based binding model, across two technical replicates. **c**, Binding curve yielding a dissociation constant of ∼0.12 nM for Trf. **d**, Modular reprogramming of the platform for p53 detection by exchanging capture strands. **e-g**, Reprogramming to p53: solution landing assay resolving tetrameric and dimeric species across three technical repeats at 5:1 p53:p53-mAb **(e)**, membrane-based titration showing successive +1c0 kDa shifts **(f)**, and binding curve with KD ∼18 nM **(g). h–j**, Reprogramming to AFP: solution landing assay resolving discrete binding transition across three technical repeats at 3:1 AFP:AFP-mAb **(h)**, membrane-based titration showing fractional occupancy fitting **(i)**, and binding curve yielding KD ∼ 350 nM **(j)**.

Using the Trf-capture DNs, we quantified binding affinity under membrane-tethered conditions. Titrations (0–400 pM) produced discrete mass shifts compatible with singly and doubly occupied states (Fig. 2a, bottom and Fig. S5). Fractional occupancies were extracted by constrained global Gaussian fitting across the titration series, and interpreted using a log-likelihood optimisation (SI Methods 13). Black and red symbols indicate the converged log-likelihood values for the four-peak and two-peak models, respectively (Fig. 2b). The resulting binding curve yields a dissociation constant of ∼120 pM (Fig. 2c).

We next assessed the platform’s programmability by reconfiguring the capture strand to target two additional proteins, p53 and alpha-fetoprotein (AFP), while maintaining the same origami scaffold (Fig. 2d-j). To evaluate sensitivity to target oligomeric state, Trf-capture DNs were reprogrammed to pull down p53 by changing the capture strand sequence and binding to an anti-p53 antibody (Fig. 1e, movie S3). Solution-phase landing assays resolved tetrameric and dimeric species, establishing oligomeric benchmarks (Fig. 2e and Fig. S2b). On SLBs (Fig. 2f and Fig. S3), titrations yielded two successive +160 kDa shifts, consistent with binding of intact tetramers^36^, with fractional occupancies fitted to yield an effective K_D_ of ∼18 nM (Fig. 3g and Fig. S6). Thus confirming the platform’s capacity to resolve and preserve native oligomeric states while being swiftly adaptable to new targets. Reprogramming for AFP produced analogous discrete binding transitions, while intermediate states were less abundant than for Trf or p53, binding was resolvable across the full concentration range, yielding a dissociation constant (K_D_ ∼0.35 µM) (Fig. 2h–j and Fig. S2c, S4 and S7), consistent with lower-affinity interactions. Together, these measurements demonstrate that the nanosensors support modular, stoichiometry-resolved binding analysis and quantitative affinity determination across a broad dynamic range using a single structural framework and simple capture-strand reprogramming.

**Figure 3.**
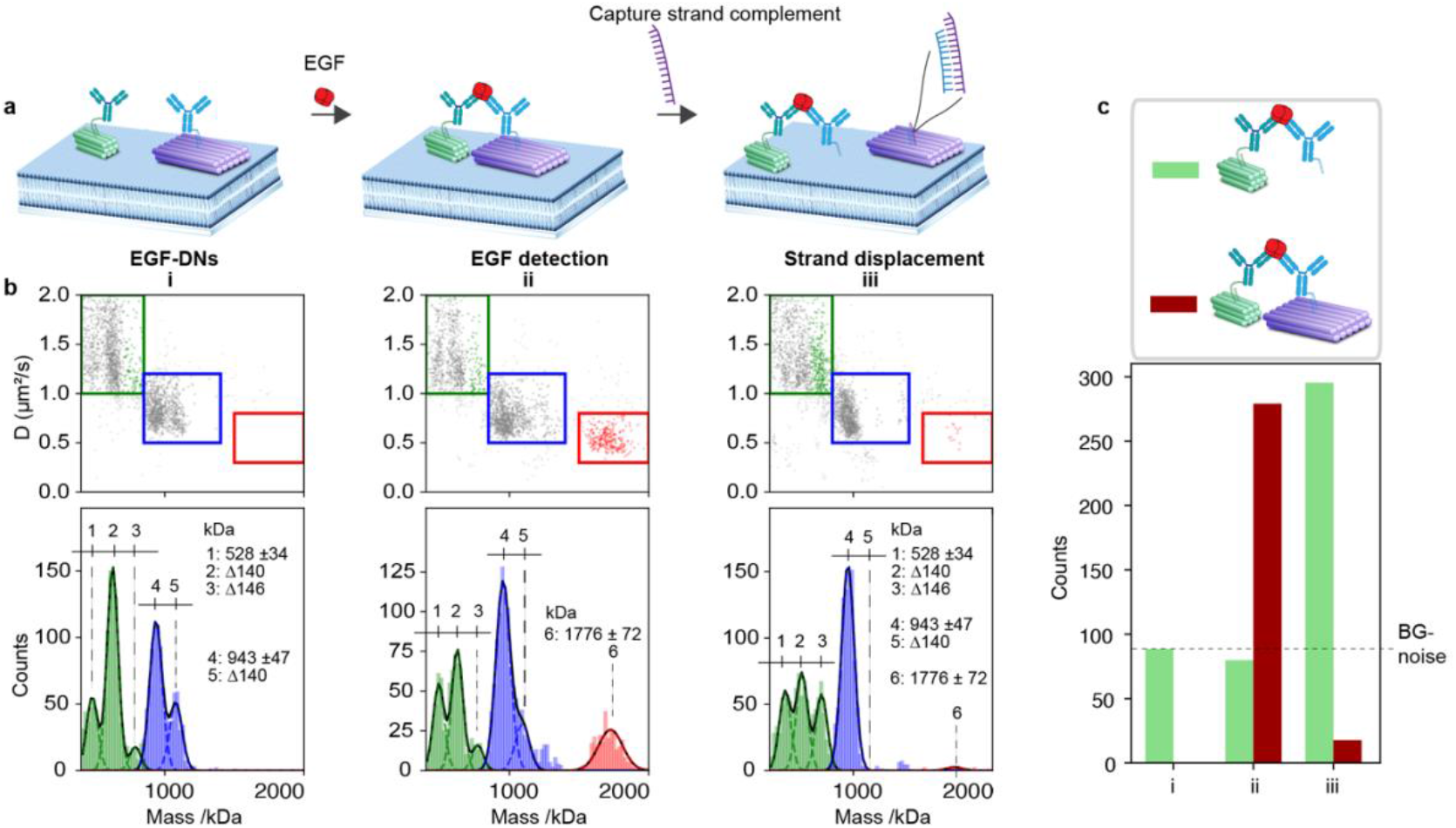
Resolving molecular compositions and programmable nanosensors’ resetting. **a**, Schematic of the dual-DN assay for EGF showing the initial state (i), EGF-induced dimerization (ii), and the selective release of a single mAb by strand displacement, programmably resetting one nanosensor (iii). **b**, Corresponding mass–diffusion plots and histograms for i–iii, showing the appearance of the dimer population and its selective removal upon strand displacement. **c**, Quantitative analysis of crosslinked (red, peak c) and strand-displaced (green, peak 3) species across the pre-EGF, post-EGF, and post-strand-displacement stages, confirming EGF detection fidelity and programmable topological switching and signal reconfiguration.

### Resolving molecular compositions and programmable nanosensor resetting

To enable simultaneous detection of multiple species, we generated scaffold fragments of defined lengths from the p8064 scaffold via restriction digestion (SI Section II, Methods 1) and varied the number of cholesterol-linked anchors, tuning both DNs mass and mobility (Fig. 3a,b and Fig. S8). To demonstrate the platform’s ability to resolve the molecular composition of detected species, we applied the dual mass–diffusion readout to detect very small targets, such as epidermal growth factor (EGF, ∼6 kDa). We exploited the lateral mobility of DNs on SLBs to design an assay in which EGF detection is evidenced by DN dimerization. We configured the origami used in the Trf-capture DN with an EGF-specific capture strand and paired it with a smaller origami carrying fewer cholesterol-anchors (528 ± 34 kDa, D = 1.2 ± 0.3 μm^2^/s).

The two nanostructures populate different ROIs in the mass–mobility plane and are functionalised with distinct mAbs targeting orthogonal binding epitopes of EGF (Fig. 3b, green and blue boxes, movie S4). Upon addition of EGF, a new population appeared at the combined DN mass and lower mobility, consistent with dual-tethered dimer formation (Fig. 3b, red box, peak 6, movie S5), confirming the detection of EGF. Crucially, unlike endpoint immunoassays—in which detection is inferred only indirectly from the amplified signal—the molecular composition of detected species is confirmed directly in our platform through its unique mass–diffusion signature, revealing the precise assembly state of each complex.

We further confirmed the specificity of this interaction using programmable toehold-mediated strand displacement^37^ to remove one of the mAbs from its DN, which produced two expected outcomes: the displaced DN reverted to its unbound state (Fig. 3b, blue box, disappearance of peak 5 in iii), and the partner DN retained both antibodies bridged by EGF (Fig. 3b, green box, peak 3 in iii). This operation simultaneously eliminated the EGF-induced dimer population (Fig. 3b, red box, disappearance of peak 6 in iii), verifying that both assembly and disassembly arise solely from specific molecular interactions. Signal-to-background analysis confirmed these topological transitions across three stages—pre-EGF, post-EGF, and post-displacement (Fig. 3c and Fig. S8c), demonstrating direct visualisation of specific assembly, and programmable, selective nanosensor resetting in real time—capabilities inaccessible to conventional endpoint immunoassays.

### Operation in complex biological media

A central limitation of most current protein-detection schemes is the need for extensive sample processing and washing to allow for specific signal generation and amplification, complicating the detection procedure and restricting detectable interactions to those with slow dissociation kinetics. We therefore evaluated the performance of our assay in clinically relevant scenarios such as human plasma and serum, using Trf-capture DNs to detect transferrin as the readout. Stepwise dilution series were performed on bare (non-functionalised) origami on the same SLB for each biological matrix (serum and plasma in separate experiments), and noise fractions were quantified as the fitted mass above the origami peak—corresponding to medium-derived protein adsorption detectable as an antibody-like component (SI Fig. S9). Noise increased beyond ∼1 % serum or plasma (Fig. 4a; ∼100-fold dilution, ∼0.65 mg mL^−1^ total protein), defining a practical dilution limit for subsequent capture experiments. At this working dilution, MP images showed elevated background (Fig. 4b). Nevertheless, the elevated mass of the DNs made them easily detectable, and the unbound origami peak (Fig. 4c, peak 1) remained clearly visible at the correct mass, serving as an internal reference that verified the absence of nonspecific mass shifts in high-background conditions.

**Figure 4.**
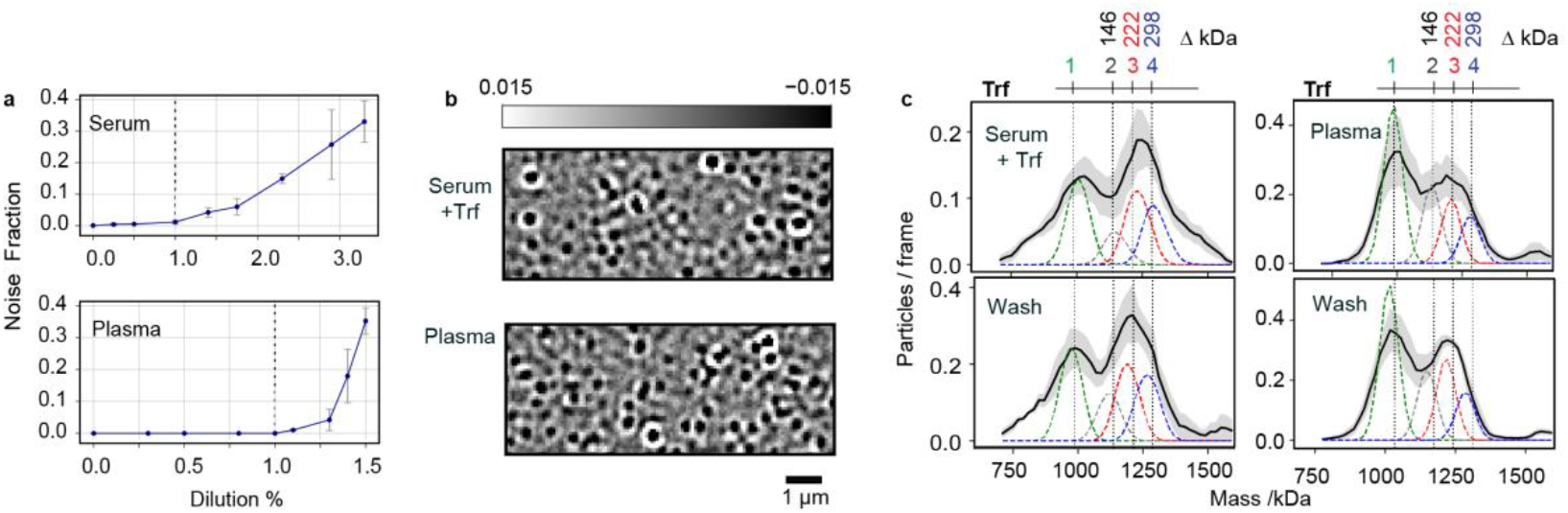
Operation in complex biological media. **a**, Noise fractions extracted from bare (non-functionalised) origami across serum and plasma dilution series. **b**, MP images of SS0 kDa Trf-capture DNs in 100-fold diluted serum and plasma, showing specific detection of transferrin spiked into serum and native in plasma. **c**, Mass distributions of Trf-capture nanosensors during complex media detection(top), and after washing (bottom) for serum and plasma. Scale bars, 1 µm.

Upon Trf binding, the detection population shifted to its characteristic single- and double-bound peaks—matching those observed in buffer (Fig. 2a)—confirming that specific Trf interactions remain directly resolvable in complex media. The coexistence of an invariant origami reference peak and correctly shifted Trf-bound peaks demonstrates that nonspecific interactions in serum or plasma do not distort mass fitting and that the detected signal arises from genuine target engagement. Importantly, dilution was the only sample preparation step; no washing, purification, or depletion of abundant proteins was required. A post-binding wash preserved the bound Trf states (Fig. 4c and Fig S10), further confirming that signal arises from stable, specific interactions rather than transient adsorption.

### Multiplexed protein analysis of biological samples

To evaluate the platform’s multiplexing capability in comparative protein analysis, a critical requirement given that diagnostic accuracy often improves through the simultaneous detection of multiple biomarkers and that clinical sample quantities are frequently limited^10–13^, we deployed a four-structure panel (DN1–DN4, Fig. S11, and movie S6) with distinct mass– diffusion encoding for multiplexed detection (Fig. 5a). Each array was exposed to lysates from four human cancer cell models: AsPC-1 and BxPC-3 (pancreatic)^38^, HepG2 (hepatocellular carcinoma)^39^, and HepG2-SupN^40^ (matched cell-free supernatant). The same four-structure encoding was retained, while the capture strands and antibody–DNA conjugates were reprogrammed in separate experiments to probe a representative set of cancer-associated proteins spanning secreted, membrane, and intracellular compartments. Secreted biomarkers included Trf ^41^ and alpha-fetoprotein (AFP)^42,43^; membrane receptors included epidermal growth factor receptor (EGFR)^44^, human epidermal growth factor receptor 2 (Her2/ErbB2)^44^, and cancer antigen (CA125/MUC16)^45^, and the intracellular tumour suppressor retinoblastoma protein (Rb)^46^ was analysed in both total and phosphorylated (pSer807/811)^47^ forms. This selection covers a broad range of molecular masses, abundances, and localisations, illustrating the platform’s ability to resolve distinct proteoforms in complex mixtures.

**Figure 5.**
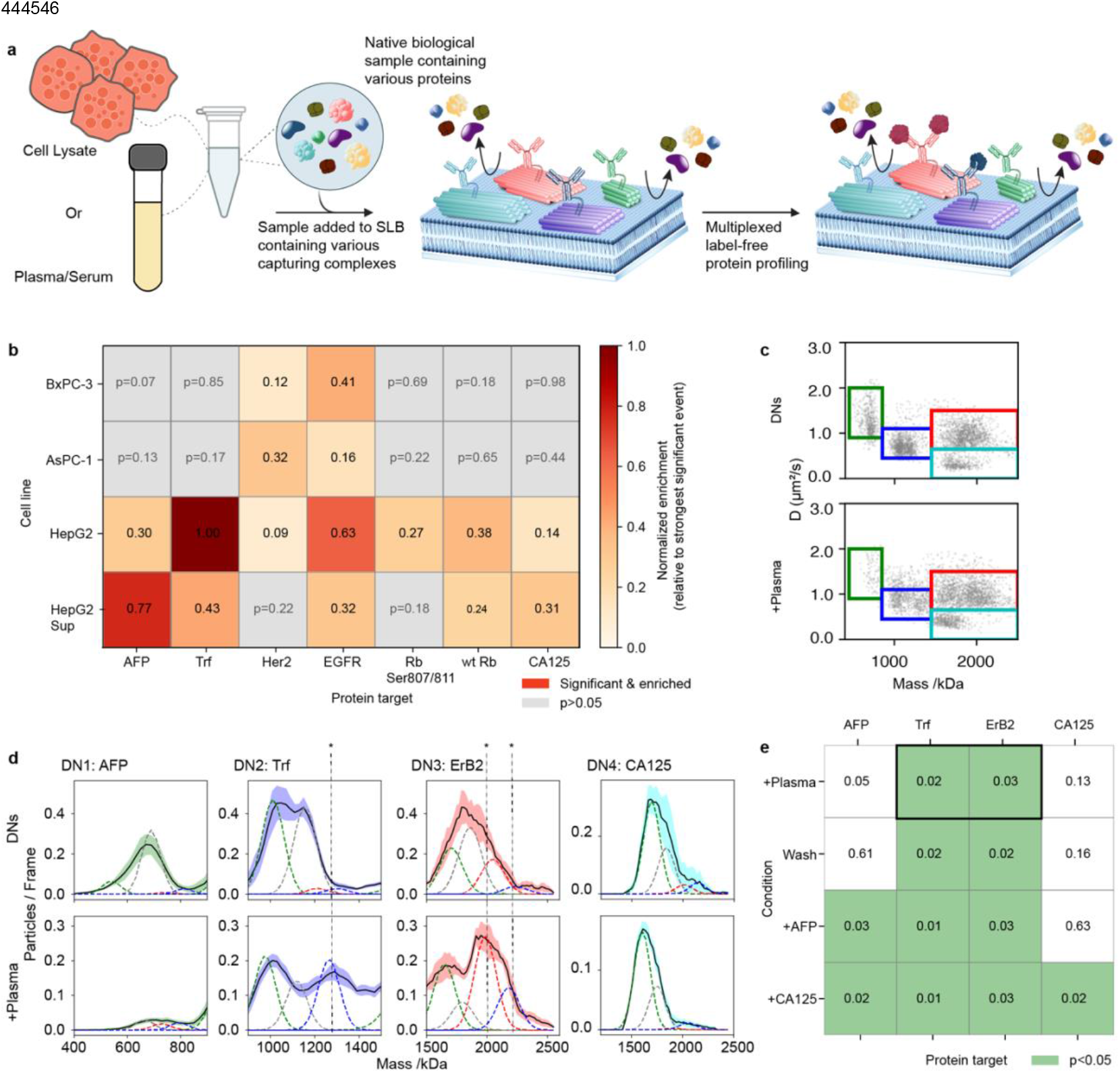
Biomarker protein profiling in human cancer cell lines and complex media. **a**, Schematic of the assay. **b**, Heatmap showing p-values from two-tailed unpaired t-tests comparing ROI occupancies between antibody-only and lysate-treated conditions (n = 3 technical replicates). Statistically significant enrichments (p < 0.05) reflect differential expression of protein targets across lysates from AsPc-1 and BxPc-3 (pancreatic), HepG2 (liver), and HepG2-SupN (secreted fraction). **c**, Mass and mobility multiplexing for a 4-panel assay. **d**, ROI-based detection of native transferrin (Trf) and spiked ErbB2 (400 pM) in 1% human plasma. Mass histograms show Trf binding in the DN2 ROI (∼1205/1280 kDa, singly/doubly bound) and ErbB2 binding in the DN3 ROI (∼2000/2200 kDa), marked by *, while AFP (DN1) and CA125 (DN4) channels show no enrichment. **e**, p-value matrix from pairwise t-tests across sequential stages, showing Trf and ErbB2 enrichment in plasma, while AFP and CA125 remained non-significant until their sequential addition.

To quantify the resulting signals, we employed a trajectory-weighted fractional occupancy metric, enabling derivation of enrichment ratios and statistical evaluation by two-tailed t-tests across replicates (SI Section II, Methods 12-15). Distinct proteome signatures emerged across the four lysates (Fig. 5b and Fig. S12-23). In HepG2 cells, Trf and AFP—both liver-associated secreted biomarkers—were clearly detected in lysate and supernatant, reflecting hepatic secretion activity. CA125 was also present and enriched in the supernatant, consistent with MUC16 ectodomain shedding^48^, whereas AsPC-1 and BxPC-3 showed no CA125 signal, in line with their negligible MUC16 expression^49^. Her2 was confined to HepG2 lysates, reflecting membrane localisation and limited shedding, while EGFR was detected across all lysates, consistent with its broad expression. Phosphorylated Rb (pSer807/811) was likewise dominant in HepG2 lysates compared to the non-phosphorylated, consistent with CDK-mediated inactivation in proliferating carcinoma cells, but was absent in the pancreatic lines, in line with Rb1 loss or deregulation typical of this cancer type^50^. Trace Rb detected in the supernatant likely reflects low-level release from apoptosis or membrane leakage. Together, these results highlight the platform’s ability to resolve tissue-specific expression profiles with high specificity and minimal sample requirements. Validation by western blotting (SI Fig. S12, S13 and S24) confirmed the MP-based detection, underscoring the platform’s capacity for low-input, label-free analysis with high information content and sensitivity to low-abundance targets in complex samples.

To enable clinically relevant validation, we evaluated multiplexed detection directly in complex biological media. Human plasma was chosen as a representative background containing abundant nonspecific proteins, into which defined biomarkers were spiked (movie S7). Distinct shifts were observed in the DN2-Trf channel (native) and in the DN3-ErbB2 channel upon spiking, while DN1-AFP and DN4-CA125 remained unchanged, confirming target specificity (Fig. 5d and Fig S25, S26). A transient reduction in the DN1-AFP signal was noted during plasma exposure, likely reflecting reduced detection of fast-diffusing, low-mass structures, but it recovered after buffer wash (SI. Fig S25). Statistical analysis (Fig. 5e) confirmed significant differences in the Trf and ErbB2 channels, whereas AFP and CA125 remained non-significant until their sequential addition that yielded further mass shifts in their respective channels, completing the four-target panel. Despite increased background, ROI-resolved trajectories enabled robust separation of bound and unbound populations, demonstrating that multiplexed, label-free detection can be performed directly in clinically relevant samples without prior purification, washing, or sample processing.

## Conclusions

We have introduced a programmable single-molecule platform that quantifies protein binding and resolves stoichiometric and oligomeric states directly in native and complex media by combining DNA nanostructures with single molecule mass photometry. We have demonstrated that mass and diffusion can serve as orthogonal encoding dimensions, enabling modular reprogramming to distinct protein targets and multiplexed detection within the same measurement environment. These results show that nanoscale DNA structures can decouple biochemical recognition from single-molecule readout and that mass-resolved trajectories provide a direct and label-free route to quantify affinity, stoichiometry, and target identity.

A defining strength of our platform is its speed and modularity. Once folded, nanostructures can be reprogrammed within minutes by exchanging capture strands, yielding multiplexed, label-free results in under two hours. Together with the growing availability of commercial origami, its large scale production^51^ and accessible design tools^52–56^, the platform enables rapid, on-demand assembly of bespoke biomarker panels without specialised instrumentation. Expanding origami geometries and anchor configurations could further broaden the accessible mass–diffusion space and increase multiplexing density.

Current capabilities already enable affinity determination across a broad dynamic range and direct operation in serum and plasma without washing or sample processing, and could be extended through incorporating fluorescence-based barcoding or logic elements, expanding the available design space for multiplexing and improving tolerance to high-background matrices^17^. Integration of alternative affinity reagents (nanobodies, aptamers, covalent binders) and improved surface passivation^57,58^ could further widen biochemical scope and support kinetic measurements at higher matrix loadings.

Beyond sensing, the combined ability to resolve stoichiometry, oligomeric state and abundance under native solution conditions opens a third direction: mechanistic interrogation, in which programmable architectures provide a tunable physical context for studying molecular interactions, assembly pathways and conformational equilibria without immobilisation or amplification. The modularity of our platform provides a general route to interrogate reaction mechanisms, binding cooperativity and state distributions in situ.

Three application domains emerge: (i) translational proteomics, where state-resolved measurements can illuminate protein regulation and signalling dynamics; (ii) diagnostic multiplexing, where biomarker panels can be assembled and read out from small sample volumes; and (iii) mechanistic analysis, where programmable single-molecule architectures enable native biochemical interrogation. Taken together, by bridging DNA nanotechnology with single-molecule mass photometry, this platform establishes a general framework for programmable, wash-free and multiplexed protein analysis with relevance across proteomics, nanotechnology and molecular diagnostics.

## Authors Contribution

**S.H**. and **P.K**. conceptualized and designed research; **S.H**. designed and prepared the origami nanostructures, performed the MP experiments, **M.M**., **D.W**., and **S.S**. provided cells; **M.M**. performed western blots; **S.H**., **R.A**., **J.C.T**, and **R.v.W**. analysed data; **S.H**., **R.A**., **J.C.T**., and **R.v.W**. provided code for the analysis; **D.W**., **M.M**., **C.V.R**., **K.C.Z**. and **J. L.P.B**. contributed new reagents and insightful discussions; **S.H**. and **P.K**. wrote the manuscript; **S.H**., **P.K**., **R.A**., **J.C.T**., **R.v.W**., **M.M**., **D.W**., **S.S**., **K.C.Z**., **J.L.P.B**., and **C.V.R**. revised and edited the manuscript; and all authors discussed results and approved the final manuscript.

## Acknowledgements

This research was funded in whole or in part by ERC PHOTOMASS 819593, EPSRC (EP/T03419X/1 and EP/W001055/1), and BBSRC Transformative Research Technologies (UKRI1877). **S.H**. is supported by ERC (Photomass 8195), EPSRC (EP/T03419X/1) and BBSRC Transformable Research Technologies Fund (UKRI1877). **R.A**. is supported by the EMBO (ALTF-198-2020) and EPSRC (EP/T03419X/1). **M.M**. is supported by MRC (MR/V028839/1). **J.C.T** is supported by EPSRC (EP/W001055/1) and a Schmidt AI Fellowship. **D.W**. is supported by Wellcome Trust grant number (221795/Z/20/Z). **R.v.W**. is supported by the Wellcome Trust Grant Number (218514/Z/19/Z). **S.S**. is funded by CSC– University of Oxford Scholarship. **K.Z**. is supported by the Wellcome Trust LEAP programme NanoQuest. **J.L.P.B**. is supported by the BBSRC sLoLa (BB/W00349X/1). **C.V.R** is supported by MRC (MR/V028839/1) and Wellcome Trust grant number (221795/Z/20/Z). **P.K**. is supported by EPSRC (EP/T03419X/1 and EP/W001055/1) and ERC (Photomass 819593). For the purpose of Open Access, the author has applied a CC BY public copyright licence to any Author Accepted Manuscript (AAM) version arising from this submission.

## Competing interests

P.K. is an academic founder, shareholder, and non-executive director to Refeyn Ltd. J.L.P.B. is an academic founder and shareholder of and advisor to Refeyn Ltd. CVR is a cofounder and scientific advisor to OMass therapeutics. All other authors declare that they have no competing interests.

## Data and code availability

The raw data and code required to reproduce all of the manuscript figures will be deposited in the University of Oxford Research Archive upon manuscript acceptance.

