## Supporting Information for "Modular DNA nanosensors enable state-aware multiplexed single-molecule mass photometry in complex media"

#### I. Materials

**Stocks and reagents:**

1,2-dioleoyl-sn-glycero-3-phosphoethanolamine-N-[methoxy(polyethylene glycol)-2000] (ammonium salt) (DOPE-PEG2k, 880130P) and 1,2-dioleoyl-sn-glycero-3-phosphocholine (DOPC, 850375P) were purchased from Avanti Polar Lipids. UltraPure 10% SDS (15553-035) was obtained from Invitrogen. Silicone gaskets (GBL103280) were purchased from Grace Bio-Labs. Glass coverslips (24 × 50 mm, Menzel Gläser, VWR 630-2603) and isopropanol (24137-M) were obtained from VWR. SiteClick Antibody Azido Modification Kit (S20026) was purchased from Thermo Fisher. HEPES (H3375), magnesium chloride hexahydrate (M2670), and KCl (P9541) were purchased from Merck Life Science UK Limited. Amicon Ultra centrifugal filters (UFC5050) were obtained from Millipore. DPBS (14040133) was from Thermo Fisher.

**Scaffold and enzymes**

Single-stranded DNA scaffold p8064 was purchased from Tilibit, and the restriction enzymes BsaBI (R0537) and XmnI (R0194) were obtained from NEB.

**Recombinant proteins and antibodies used for MP experiments** were obtained from the following suppliers:

EGF (236-EG, R&D Systems);
EGFR antibody (MABF120, Merck);
p53 mAb (DO-1, sc-126, Santa Cruz Biotechnology);
HER2 (trastuzumab) human IgG1 (her2tra-mab1, InvivoGen);
Phospho-Rb (Ser807/811) antibody (41359SF, Cell Signaling Technology);
Rb (4H1) mAb (61121SF, Cell Signaling Technology);
Anti-human transferrin (100440_1 mg, Medix Biochemica);
Anti-human CA125 (100598_1 mg, Medix Biochemica);
Anti-AFP [AFP-01] (ab3980, Abcam);
Anti-EGFR clone 225 (MABF120, Merck);
Human EGF Antibody Pair (ab241876, Abcam);
Recombinant ErbB2/HER2 protein (ab168896, Abcam);
Recombinant human ErbB2 Fc chimera (1129-ER-050, R&D Systems);
Anti-ErbB2/HER2 [EPR19547-12] (ab222482, Abcam);

Recombinant p53 (81091, Active Motif via Cambridge Bioscience);
Human apo-transferrin protein (3188-AT, R&D Systems).

**Biological media** included human plasma (P9523-1ML, Merck) and fetal bovine serum (26140-087, Thermo Fisher; 12103C-100ML, Merck)

**Cell lines**
AsPC-1 cells (ATCC CRL-1682)
BxPC-3 cells (ATCC CRL-1687)
HepG2 cells (ATCC HB-8065)

**Lysis and sample preparation**
Pierce IP Lysis Buffer (Thermo Scientific, 87788)
Amicon Ultra 50 kDa centrifugal filters (Millipore)

**SDS–PAGE gels and running buffer**
NuPAGE 4–12% Bis-Tris Mini Protein Gel (Invitrogen, NP0321BOX)
NuPAGE MES SDS Running Buffer (Invitrogen, NP0002)
**Western blot membrane and transfer**
PVDF membrane (Millipore IPVH00010)
Trans-Blot transfer system (Bio-Rad)

**Blocking and washing reagents**
Non-fat dry milk
TBST (Tris-buffered saline with Tween-20)

**Primary antibodies for Western blot**
Anti-AFP (SC-8399, Santa Cruz)
Anti-HER2 (SC-08, Santa Cruz)
Anti-EGFR (AHR5072, Thermo Fisher)
Anti-Transferrin (100440, Medix Biochemica)
Anti-CA125 (SC-365002, Santa Cruz)
Anti-p53 (9282S, Cell Signaling Technology)
Anti-Rb (SC-102, Santa Cruz)
Anti-GAPDH (2118S, Cell Signaling Technology)

**Secondary antibodies**
HRP-conjugated secondary antibodies (Abcam or Life Technologies)

**Detection reagents**
ECL chemiluminescence solution (Thermo Fisher)
iBright 1500 Imaging System (Invitrogen/Thermo Fisher)

**DNA Oligos**
All DNA oligonucleotides were purchased from Integrated DNA Technologies (IDT), except for the 3’ Cholesterol oligo was purchased from Biomers.

### II. Methods:

#### 1. Scaffold digestion:

Different-length scaffold fragments (Fig. 1, S1, and S11 a,b) were prepared using the following protocol, adapted from a previously published method and modified for the present study[1]; all scaffold fragments and digestion oligo sequences are listed in Section IV (Sequences).

1.1. 1496-nt fragment: To generate the 1496-nt fragment from p8064, a 200 µL double-digestion reaction was prepared containing p8064 scaffold at a final concentration of 50 nM, digestion-staple oligonucleotides added at twenty-fold molar excess (1 µM; fragment-specific staple set), 1× rCutSmart buffer, 10 µL XmnI (NEB R0194), and nuclease-free water to 200 µL. The reaction was divided into two 100 µL aliquots, heated to 85 °C for 5 min, and incubated at 37 °C for 90 min. Each aliquot then received 5 µL BsaBI (NEB R0537) and was incubated at 60 °C for 90 min, followed by enzyme inactivation at 80 °C for 20 min. In this design, XmnI cuts at position 2988 and BsaBI at 1492.

1.2. The 796-nt fragment was generated using the same procedure and the corresponding digestion-staple set, with XmnI cutting at position 700 and BsaBI at 1492.

1.3. The 2288-nt fragment used its respective staple set and XmnI only, cutting at positions 700 and 2988.

1.4. The 2824-nt fragment used the appropriate staple set and combined XmnI cleavage at position 2988 with BsaBI cleavage at positions 1492 and 4316.

No purification of the scaffold fragments was required. The digestion mixtures were stored at −20 °C and used directly for the assembly reactions of the different origami structures.

#### 2. DNA origami design

Four DNA origami designs were used in this study (Fig. S11 c-I, S28-31). In all cases, staples on the top layer carried 3′ extensions encoding distinct capture sequences, enabling programmable target recognition through hybridisation to complementary mAb–DNA conjugates. Staples on the bottom layer carried 5′ extensions complementary to a universal 3′ cholesterol-modified oligonucleotide for SLB anchoring. The number of 5′ extensions defined the number of SLB linkages and thus tuned the diffusion of each structure.

2.1. The first design consisted of two stacked layers of six parallel helices arranged on a square lattice and was folded from a 1496-nt custom scaffold fragment, yielding a structure measuring ~43 × 4 × 16 nm (Fig. S28). This design incorporated ten 5′ SLB-linker staples (Fig. S28, green).

2.2. The second design, folded from the 792-nt scaffold fragment, comprised three layers of four helices arranged on a square lattice and measured ~21 × 8 × 6 nm (Fig. S29). It incorporated six SLB-linker staples.

2.3. The third design, folded from the 2288-nt scaffold fragment, consisted of two layers measuring ~50 × 4 × 16 nm (Fig. S30) and featured twenty-eight SLB-linker staples.

2.4. The fourth design, folded from the 2824-nt scaffold fragment, formed a four-layer rectangular structure measuring ~54 × 8 × 10 nm and included eight SLB-linker staples (Fig. S31).

3. **DNA origami assembly:**

All origami structures were assembled by combining 20 nM of the scaffold-digestion mixture containing the appropriate scaffold fragment with each staple oligonucleotide at 200 nM in folding buffer (10 mM Tris, 1 mM EDTA, pH 8.0, 16 mM $MgCl_2$). The mixture was heated to 65 °C for 20 min and then cooled to 20 °C over 16 h, using a linear cooling ramp of 1 °C every 45 min. Each origami design was folded in a separate batch reaction.

No purification was performed. Following assembly, structures were mixed with the 3′-cholesterol-modified oligonucleotide at a 5:1 molar ratio of cholesterol oligo to origami.

4. **Agarose gel:**

The digestion reactions and resulting scaffold fragments were analysed on 2% agarose gels prepared in 1× TAE buffer (40 mM Tris, 40 mM acetic acid, 1 mM EDTA, pH 8) supplemented with 11 mM $MgCl_2$. Electrophoresis was performed at 60 V for 2 h on ice to.

5. **Antibody-DNA conjugation:**

The monoclonal antibodies were site-specifically functionalized with DNA following a two-steps procedure:

5.1. **Antibody-Azide functionalization:** Monoclonal antibodies were functionalised with azide groups using the SiteClick™ Antibody Azido Modification Kit according to the manufacturer's protocol. Briefly, 100–250 µg of antibody was concentrated to ~50 µL and buffer-exchanged into the supplied antibody preparation buffer. Terminal galactose residues on Fc N-linked glycans were removed by incubation with β-galactosidase at 37 °C overnight. Azido-modified GalNAz sugars were then added enzymatically by mixing the antibody with UDP-GalNAz, Tris buffer, buffer additive, and GalT enzyme, followed by overnight incubation at 30 °C. The azide-modified antibody was purified and concentrated using the provided 50 kDa filters and collected at 1–5 mg/mL (OD280). The azide-modified antibody (antibody-N3) was stored at 4 °C for subsequent DNA conjugation.

5.2. **Antibody–DNA conjugation:** azide-modified antibody was mixed with DBCO-modified DNA oligonucleotides at a 1:5 antibody-to-DNA molar ratio and incubated overnight at 25 °C. Conjugation mixtures were purified using 50 kDa Amicon Ultra centrifugal filters to remove unreacted oligonucleotide. Filters were pre-rinsed with 500 µL DPBS and centrifuged at 14,000 g for 5 min at 4 °C, and the flow-through was discarded. The reaction mixture was then added to the filter, the volume was adjusted to 500 µL with DPBS, and the sample was centrifuged again at 14,000 g for 5 min at 4 °C; the flow-through was discarded. A further 480 µL DPBS was added to the filter and centrifuged at 14,000 g for 5 min at 4 °C; the flow-through was discarded. This wash step was repeated four times. Conjugates were recovered by inverting the filter into a clean tube and centrifuging at 1,000 g for 3 min. Final mAb–DNA conjugates were stored at 4 °C, and concentrations were measured using a NanoDrop spectrophotometer and stored at 2µM stocks.

6. **Supported lipid bilayer (SLBs) preparation**

SLBs were prepared following procedures similar to those previously described[4]. Lipid mixtures were prepared from chloroform stocks to generate a $x10$ solution containing 0.07 mM DOPE-PEG2k and 4.93 mM DOPC (molar ratio 1.4:98.6). The stock was stored at −20 °C. For SLB formation, 50 µL of the lipid stock was diluted into 200 µL chloroform in a clean

glass tube (washed sequentially with Milli-Q water, 10% SDS, Milli-Q water, and acetone, then dried). Chloroform was removed by gentle nitrogen flow while rotating the tube.
The dried lipid film was rehydrated with 0.5 mL buffer (20 mM HEPES pH 7.4, 100 mM KCl) and subjected to two cycles of incubation at 45 °C for 20 min with brief vortexing between cycles. The hydrated lipids were transferred to a 1.5 mL microcentrifuge tube and tip-sonicated on ice using a 2 mm probe (30% amplitude, 1 s on / 3 s off, total sonication time 10 min). Sonicated vesicles were centrifuged at 21,130 g for 30 min at 4 °C, after which 400 µL of the supernatant was collected.
Glass coverslips (24 × 50 mm) were cleaned as for standard MP measurements using three steps of 5 min bath sonication in water (Milli-Q® water), 50% isopropanol in water and water again, and then dried under a stream of nitrogen gas. The cleaned glass coverslips were then treated with oxygen plasma for 3 min at 40% power and 0.6 mbar $O_2$ (Zepto plasma cleaner, Diener Electronic). Immediately after plasma cleaning, a silicone gasket was placed at the centre of the coverslip. A mixture of 30 µL buffer (20 mM Tris pH 7.8, 150 mM NaCl, 2 mM $MgCl_2$) and 20 µL lipids was added into the gasket and mixed thoroughly. After 20 min incubation at room temperature to allow bilayer formation, excess vesicles were removed by multiple washes with DPBS.

### 7. Dynamic MP measurements

Dynamic MP measurements were performed on a oneMP mass photometer (Refeyn Ltd.) using the large field-of-view mode (10.8 × 10.8 µm²) at a 360 Hz frame rate. Following SLB formation and removal of excess vesicles, 0.5 µL of each assembled, cholesterol-modified origami structure was introduced into ~80 µL buffer within the gasket. Surface density was adjusted by varying incubation time, after which the sample was washed eight times with 60 µL DPBS supplemented with 5 mM $MgCl_2$ to remove unbound material. Each dataset consisted of a 60 s acquisition. Although unpurified assembly mixtures containing residual digestion and folding components were used, only cholesterol-modified origami remained SLB-bound, as passivation excluded all non-lipid-anchored material.

7.1. DN assembly on the SLB: mAb–DNA (50 nM final concentration) was added to ~80 µL buffer in the gasket and incubated for 5 min with the microscope lid open to avoid continuous illumination. To limit evaporation, the sample was covered with a chamber lid containing a wet paper-towel insert. Excess antibody was removed by eight sequential 60 µL DPBS washes. To assemble the multiplexed-DN panel, the same procedure was followed, except that a mixture of mAb–DNA conjugates was added to the SLB containing the different origami nanostructures and incubated for 5 min before washing, analogous to the single-structure DN assembly.

7.2. Transferrin (Trf) detection on SLB (Fig. 1, 2 a-c and S5): recombinant Trf was added stepwise up to a final concentration of 400 pM, to the solution above the SLB (~80 µL) containing Trf-DNs, with 5 min incubation after each addition prior to acquisition.

7.3. p53 detection on SLB (Fig. 2 f-g and S7): recombinant p53 protein was added to a final concentration of 40 nM to the SLB containing p53-DNs and incubated for 5 min before acquisition.

7.4. AFP detection on SLB (Fig. 2 i-j and S8): recombinant AFP protein was added to a final concentration of 250 nM to the SLB containing AFP-DNs and incubated for 5 min before acquisition.

7.5. EGF detection (Fig. 3): EGF-DNs were assembled sequentially following the procedure in SI 8.1, using EGF–antibody–pair, each functionalized with a unique

DNA sequence. Recombinant EGF (2 nM final concentration) was then added to the solution above the assembled DNs (~80 µL) and incubated for 5 min prior to acquisition.

7.6. Serum experiment with Trf (Fig. 4 b-c): 1% fetal bovine serum (FBS; ~0.5 mg/mL background protein) was spiked with 200 pM Trf and added above the gasket, followed by a 5 min incubation. Wash recordings were obtained after eight washes with 60 µL DPBS + 5 mM $MgCl_2$.

7.7. Plasma experiment with Trf (Fig. 4b-c): 1% human plasma (~0.66 mg/mL background protein) was added above the DNs and incubated for 5 min prior to acquisition. Wash recordings were obtained after eight washes with 60 µL DPBS + 5 mM $MgCl_2$.

7.8. The multiplexed cell lysate experiments (Fig. 5b and Fig. S12-S23): cell lysates were added at ~ 20µg total protein concentration (5µl at ~ 4µg/µl concentration) to the SLB containing 4 structures DNs for specific targets and incubated for 10 min. Washing consisted of two exchanges with 60 µL DPBS + 1.5 M NaCl, followed by six exchanges with 60 µL DPBS + 5 mM $MgCl_2$, before acquisition.

7.9. Multiplexed detection in plasma (Fig. 5c–e and Fig. S25-S26 ): 1% human plasma (~0.66 mg/mL background protein) spiked with 0.5 nM ErbB2 was added above the DNs and incubated for 5 min prior to acquisition. Wash recordings were obtained after eight washes with 60 µL DPBS + 5 mM $MgCl_2$. AFP (200 nM) and CA125 (400 pM) were then added sequentially, each followed by a 5 min incubation before acquisition.

### 8. Landing assay MP-measurements

The landing assay experiments were carried out on a commercial mass photometer (TwoMP, Refeyn Ltd.) using the regular field of view (10.9 × 4.3 µm²) with an acquisition frame rate of 500 Hz, followed by 2× frame binning to yield an effective frame rate of 250 Hz.

All samples were measured in DPBS. Before each measurement, a gasket (GBL103250, Grace Bio-Labs) attached to a coverslip was filled with DPBS and placed on the microscope stage, and the focus was adjusted. The protein solution was then added to the buffer-filled gasket, and a 60 s video was recorded immediately to capture landing events of individual proteins on the coverslip surface.

8.1. Trf / Trf–mAb interaction assay (Fig. 2a (top) and S2): Trf–mAb was mixed with recombinant Trf at different molar ratios. In all experiments, Trf–mAb was kept at 200 nM and mixed either with 100 nM Trf (0.5:1 molar ratio) or with 300 nM Trf (1.5:1 molar ratio) and incubated for 5 min. The mixture was added to the gasket-contained DPBS, resulting in a tenfold dilution (20 µL final volume). Three technical replicates were acquired for each condition.

8.2. p53 / p53–mAb interaction assay (Fig. 2e and S2): p53–mAb (80 nM) was mixed with recombinant p53 (400 nM) to yield a 5:1 protein-to-antibody ratio and incubated for 5 min. The mixture was then added to the gasket for a tenfold dilution, and three technical replicates were acquired.

8.3. AFP / AFP–mAb interaction assay (Fig. 2h and S2): AFP–mAb (100 nM) was mixed with recombinant AFP (300 nM) to yield a 3:1 protein-to-antibody ratio and incubated for 5 min. The mixture was then added to the gasket for a tenfold dilution, and three technical replicates were acquired.

8.4. Individual recombinant proteins and mAbs were analysed separately prior to mixing. For these controls, protein or antibody was added to the gasket containing buffer to final concentration of 10 nM after a tenfold dilution (20 µL final volume).

For all measurements, the interval between sample loading and acquisition was <10 s.
Movie processing and event detection were performed using DiscoverMP (V2023 R1.2, Refeyn Ltd.), and movies were generated using a rolling window of 10 frames (25 ms integration time). Particle-detection thresholds were kept at their default values (threshold 1 = 1.5, threshold 2 = 0.2). Measured contrasts were converted to mass using a protein calibrant as described previously[2,3].

### 9. Dynamic MP Image analysis:

All movies were analysed using a custom-written Python script adapted from previously published dynamic MP analysis workflows[2,3], with modifications to reflect the acquisition parameters used in this study. For each dataset, raw pixel values were first converted to photoelectron counts and each frame was normalised to its total number of accumulated photoelectrons. Unlike earlier implementations that applied temporal binning, here all analyses were performed on the native 360 Hz data to preserve the full temporal resolution of particle motion.
To remove static background contributions arising from glass roughness, we employed a sliding-median ratiometric background subtraction. For each pixel at time $t$, a median value was computed over a 601-frame window (equivalent to ~1.67 s at 360 Hz, using 300 frames on either side of $t$). Each frame was then transformed by subtracting and dividing by this median value, which suppresses static or slowly varying background features while retaining the dynamic interferometric signals generated by freely diffusing particles. After temporal background correction, low-frequency spatial variations were reduced using a spatial median filter applied to each frame.
Particle detection proceeded by convolving each background-corrected frame with a Laplacian-of-Gaussian (LoG) kernel. Candidate detections were identified as pixels exceeding a fixed contrast threshold of 0.0012, and that simultaneously satisfied a local-maximum condition. These pixels were taken as provisional particle centres. For each detection, a 13 × 13-pixel region of interest (ROI) centred on the candidate position was extracted and fitted with a point-spread-function (PSF) model. The fitting procedure estimated the particle contrast by optimising the PSF amplitude together with its x–y position, using least-squares minimisation between the observed ROI and the model. This yielded frame-resolved particle localisations and contrast values suitable for subsequent trajectory linking and mass-diffusion analysis. The PSF model is described in detail in the Materials and Methods of ref 2.

### 10. Generating a trajectory from consecutive localisations

To connect individual successful and consecutive fitting events across adjacent frames into individual molecular trajectories of diffusing particles, similar to ref.[2,3], we used the linking function of the *trackpy* Python package (trackpy.link_df). For our use, there are three relevant parameters: (1) The maximum distance between two consecutive localisations that can be linked to the same particle. In our analysis, this value was set to 5 pixels. (2) A memory parameter, that allows gaps in individual trajectories to be filled owing to particles that transiently disappear below the detection limit or owing to fitting errors. Here, the memory was set to one frame. (3) A minimum localisation-quality threshold that determines which fitted positions are sent to the linking procedure. Localisations were retained only if the refined

position deviated by less than 2 pixels from the initial peak estimate and if their fitted contrast was below 0.0012.
These parameters were used to assemble the fitted positions into individual particle trajectories.

### 11. Extraction of Mass and Diffusion Coefficient

Detected trajectories and molecular segments were analysed similarly. Here we considered all trajectories that were longer than 139 ms (corresponding to 50 frames at an effective frame rate of 360 Hz) for the calculation of diffusion coefficients. For shorter trajectories, no mobility estimate was included. The molecular mass of each trajectory or segment was calculated as the median of its mass trace, which reduces sensitivity to local noise or fluctuations. For trajectories where a diffusion coefficient was determined, the assigned mass was corrected to account for motion blur, which lowers the apparent contrast of mobile particles due to their movement during exposure. The correction was applied automatically during analysis based on the measured diffusion coefficient. The mass of trajectories for which no diffusion coefficient was assigned was not corrected. The contrast was converted to mass using a calibration factor of $2.7 \times 10^{-5}$ per kDa.[4] This procedure allows for accurate estimation of both diffusion and mass, sufficient to resolve the expected oligomeric species in the dataset.

### 12. Plotting Mass Histograms and Calculating Surface Molar Fractions

To calculate the relative abundance of different oligomeric species, we generated weighted mass histograms from the trajectory dataset. To reduce the impact of noise at low masses, we considered only trajectories longer than 50 frames (139 ms at 360 Hz). The contribution of each trajectory or segment was weighted by its length (in frames) and normalized by the total number of frames in the movie. This produced a mass histogram in which the x-axis represents molecular mass, and the y-axis corresponds to the average number of detected particles per frame in each mass bin.
The histograms were then fitted using a four-component Gaussian model corresponding to the expected species: (1) origami only, (2) origami bound to antibody, (3) origami–antibody bound to one protein, and (4) origami–antibody bound to two proteins. The expected relative mass positions of these species were hold constant while the absolute mass was scaled by a single global factor, which was optimised during fitting. This ensured that the fitted mass of the origami-only population defined the absolute mass scale, while the positions of the remaining peaks were automatically adjusted according to predefined relative offsets.
Each Gaussian was assigned an independent amplitude and standard deviation (σ). The initial σ values were set to 5% of the expected origami mass and constrained to remain within a physically meaningful range. This approach ensured consistent and realistic peak widths across datasets. Fitting was performed by minimizing a chi-squared loss function comparing the observed histogram to the sum of Gaussian components.
The relative abundance of each species was obtained from the area under its corresponding Gaussian. The average protein occupancy per antibody-containing origami was calculated externally as:

$$\chi = \frac{\rho_1 + 2\rho_2}{\rho_0 + \rho_1 + \rho_2}$$

where $\rho_n$ is the abundance of origami–antibody complexes bound to $n$ proteins. This formulation provides the average number of proteins per antibody-containing origami structure. The origami-only population was excluded from this calculation.

### 13. Model-Based Extraction of Fractional Occupancies

To evaluate whether the additional two peaks corresponding to the singly and doubly bound states are statistically significant, we compared the fitting performance of two alternative models. The first model included only two mass peaks, representing the free origami structure and the origami bound to a single anti-transferrin antibody. The second model incorporated all four possible mass peaks, thereby accounting also for the molecular states in which one or two transferrin molecules are bound. Both models additionally included a constant background term to represent uniform noise contributions.

We calculated the Poisson likelihood chi-square values for each model as a function of the transferrin concentration in solution (Fig. 2 and S5), using the expression[5]:

$$\chi^2_{\lambda,p} = 2\sum_{i=1}^{Nbins} y_i - n_i + n_i \log\frac{n_i}{y_i}$$

Here, $y_i$ and $n_i$ are the modelled and experimental values of the i-th histogram bin. Fig. 2b, S1 e, show that while the likelihood of the two-peak model decreases rapidly with increasing transferrin concentration, the likelihood of the four-peak model remains constant across the full range of concentrations. Based on these results, we define a detection threshold for transferrin as the concentration at which the two curves diverge. Here, we set the detection limit to be 60 pM.

### 14. Fitting data of multiplexing

Fitting of the mass histograms from the multiplexing experiments was performed as follows. Prior to each set of measurements, the diffusing origami structures were measured in their reference state in buffer solution. The two-dimensional mass–diffusion histograms of the origami structures were divided into two ROIs: one containing structures 1–3 (with masses of 550, 990, and 1850 kDa) and a second ROI containing the large, slowly diffusing origami at 1550 kDa (see Fig. S11, S16-26). The resulting mass histograms were fitted to four Gaussian functions, providing reference masses and peak line shapes for subsequent analysis, described by the Gaussian function:

$$P_i(\mu_i, \sigma_i) = A_i G(\mu_i, \sigma_i)$$

Where $P_i$ is the peak line shape of the i-th origami structure and $\mu_i, \sigma_i$ and $A_i$ are its mean mass, standard deviation and amplitude, respectively. Following the fitting of the reference measurements, the masses ($\mu_i$) and standard deviations ($\sigma_i$) were fixed.

To fit the mass histograms of the experiment described in Fig. 5 and Fig. S11-S26, the mass histograms of the two ROIs were fitted to the following sums of calibrated Gaussian functions:

$$PD_{ROI_1}(\{A_{i,j}\}, s) = \sum_{i=1}^{3}\sum_{j=1}^{4} A_{i,j} G(s \cdot \mu_{i,j}, s \cdot \sigma_i)$$

$$PD_{ROI_2}(\{A_{4,j}\}, s) = \sum_{j=1}^{4} A_{4,j} G(s \cdot \mu_{4,j}, s \cdot \sigma_4)$$

Here, $\mu_{i,j}$ are the masses associated with the i-th origami structure in one of its four molecular states: free (j=1), bound to one antibody (j=2), bound to an antibody with one bound antigen (j=3), or bound to an antibody with 2 bound antigens (j =4). The masses of the antibodies and antigens were determined using a standard mass photometry assay and are listed in Table 1. The $\sigma_i$ are the calibrated standard deviations of the origami mass peaks, The parameters $A_{i,j}$ are the amplitudes of the peaks corresponding to the different molecular species, and $s$ is a global mass scaling factor that accounts for variation in the adjusted focus position between measurements, which results in slight differences in the optical-contrast to mass conversion. The fitting parameters are the set of amplitudes, $A_{i,j}$ and the global scale, $s$. The scale factor $s$ was constrained to vary between 0.95 and 1.05, corresponding to a maximum allowed deviation of 5% in the contrast-to-mass conversion. The optimal parameters were obtained by minimising:

$$f = \frac{1}{N_{bins}} \sum_{n=1}^{N_{bins}} (PD_{ROI}(\{A\}, s)_n - Data_n)^2$$

where, $Data_n$ is the measured occurrence of particles in the n-th bin of the mass histogram, and $PD_{ROI}(\{A\}, s)_n$ is the corresponding modelled value. The minimisation was performed using *scipy.optimize.minimize* python package.

### 15. Quantification of Target Enrichment and Heat Map Generation

To quantify target engagement across proteins and cell lysates, we calculated enrichment values based on changes in fractional occupancy before and after lysate exposure. For each protein–lysate pair and technical replicate, the fractional occupancies of singly and doubly bound species were summed and normalized to the total antibody population, yielding a combined binding fraction. The enrichment ratio was defined as the mean post-lysate fraction divided by the mean pre-lysate fraction across three technical replicates, providing a measure of effect size.

To assess statistical significance, we applied a two-tailed t-test to compare the pre- and post-lysate binding fractions across replicates. That is, for each protein–lysate pair, the three replicate values measured before lysate exposure were compared to the three values measured after exposure. Each matrix entry was therefore characterized by an enrichment ratio and a corresponding p-value. Pairs with an enrichment ratio greater than 1 and a p-value below 0.05 were considered significantly enriched (Table 2).

Only these significant enrichments were used to define a global normalization constant, corresponding to the highest enrichment ratio observed among all significantly enriched protein–lysate pairs. Each significant enrichment value was then divided by this maximum to yield a normalized score between 0 and 1, which determined the color intensity in the heat map. Entries that did not meet the enrichment threshold (enrichment ratio less than 1) or the statistical significance criterion (p-value greater than or equal to 0.05) were shown in grey, to indicate that they were tested but not enriched.

In some cases, background signals in the DNA-nanosensors (DN) movies prior to lysate exposure could be spuriously fitted as species resembling antibody–protein complexes, particularly in the mass range corresponding to antibody bound to one or two proteins. These artefacts can distort the pre-lysate baseline and affect the outcome of the t-test, potentially masking real enrichment. To address this, we incorporated both statistical significance and enrichment value in our criteria, ensuring that enrichment was only reported when both metrics supported a genuine increase in binding upon lysate addition.

By considering both the magnitude of enrichment and its statistical significance, this analysis minimized false positives that could result from background noise—such as particles incorrectly assigned to the expected mass range of protein-bound species despite while not representing genuine complexes—and ensured that only reproducible, meaningful changes in binding were interpreted as specific target engagement.

**16. immunoblotting assay:**

Cells were purchased as AsPC-1 (CRL-1682), BxPC-3 (CRL-1687) and HepG2 (HB-8065). Cells were cultured in T75 flasks and allowed to reach at least 85% coverage before harvesting. Cells were harvested, centrifuged at 600 g for 5’, and 100 µL lysis buffer was added to lyse the cells. For HepG2 cells, supernatant was collected and concentrated from 12 mL to 1 mL with a 50 kDa centrifugal filter. About 40 µg of total protein (measured by nanodrop) was loaded on a gradient 4–12% Bis-Tris mini protein gel, in MES-SDS running buffer, resolved at 200 V and electroblotted to activated polyvinylidene fluoride (PVDF) membrane using a transfer system. The membranes were blocked with 3% non-fat milk solution in Tris-buffered saline and Tween 20 (TBST) for 1 hour at room temperature. Membranes were incubated overnight at 4 °C with either anti-AFP (1:400), HER2 (1:400), EGFR (1:400), Trf (1:400), CA125 (1:400), p53 (1:400), Rb (1:400), or GAPDH (1:400) primary antibody diluted in blocking solution. Membranes were washed and then incubated with peroxidase-conjugated secondary antibodies (1:10,000) diluted in blocking solution for 1 hour at room temperature. The membranes were washed with TBST and proteins were visualized using enhanced chemiluminescence (ECL) solution on a chemiluminescence imaging system.

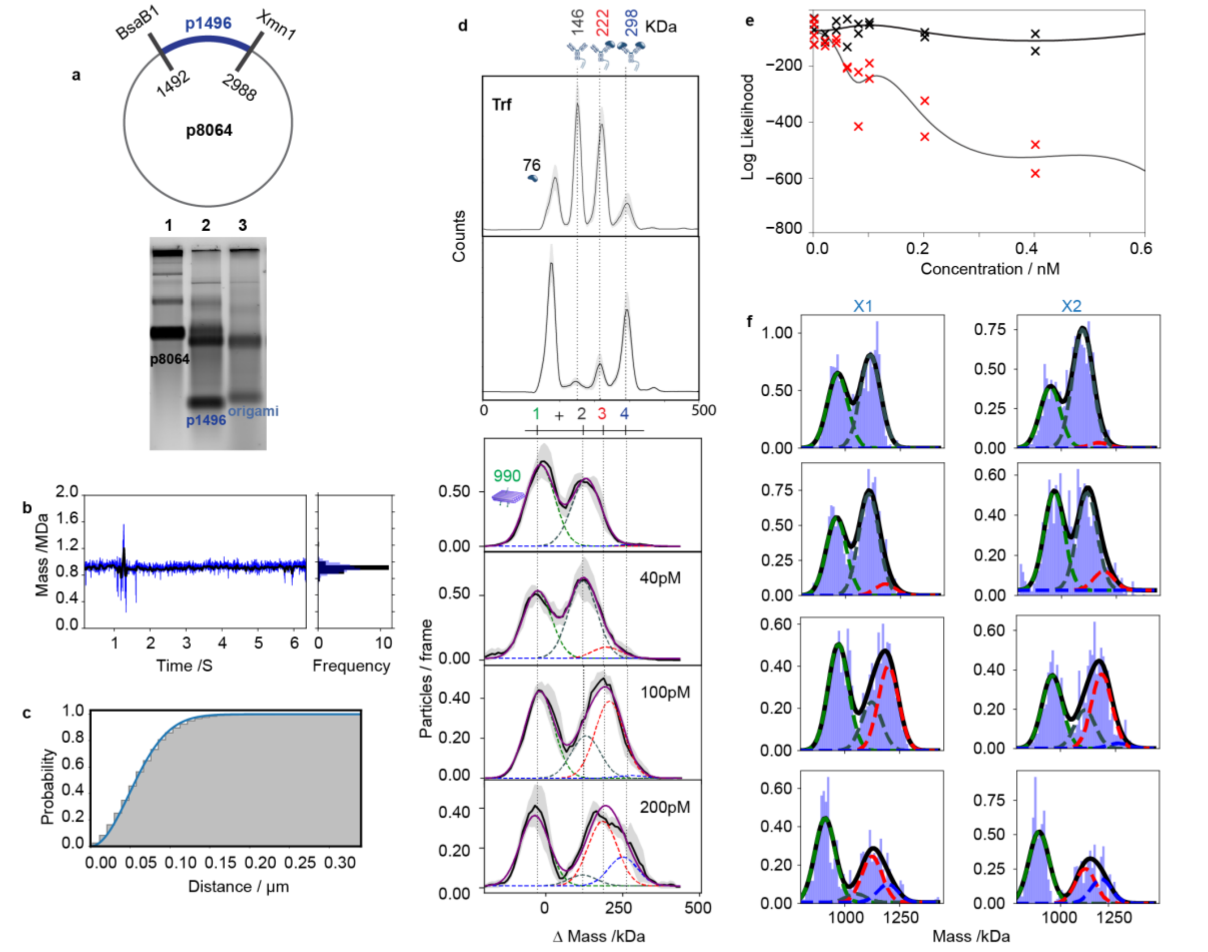

***Fig. S1: Engineering a membrane-tethered DNA-origami nanosensor for single-molecule mass photometry.***
***a, Scaffold preparation and origami folding.*** *Data correspond to Fig. 1. Top: Schematic of p8064 digestion using BsaBI and XmnI to excise the 1,496-nt scaffold fragment. Bottom: Agarose gel electrophoresis verifying successful digestion (lane 2) and assembly of the ~1 MDa DNA-origami nanostructure (lane 3).*
***b, Dynamic MP mass trace.*** *A representative single-particle mass trace is recorded at 360 Hz (blue), with a corresponding mass histogram displayed after binning of 10 (28 ms) in black (data correspond to Fig. 1c).*
***c, Diffusion analysis****. Cumulative probability distribution of the distance travelled by a single origami nanostructure during a single frame within its measured trajectory. The corresponding time interval for particle displacement was 2.8 ms. The blue curve corresponds to the best fitted model used to extract the diffusion coefficient.*
***d, Stoichiometric assignment of Trf binding.*** *Top: Reference landing assay resolving free transferrin (Trf, 75 kDa), unbound DNA–mAb conjugate (146 kDa), and Trf-bound antibody conjugates carrying one (221 kDa) or two (298 kDa) proteins, at two protein-to-antibody ratios (0.5:1 and 1.5:1 Trf:mAb), across three technical repeats. Bottom: Mass distributions of SLB-tethered DNA origami before and after antibody conjugation and Trf binding following a sequential increase in the titrated-Trf concentration (data correspond to Fig.1e). Gaussian components correspond to the unmodified origami (green), antibody-bound origami (DN, grey), and DN bound to one Trf protein (red) and DN bound to two Trf protein (blue) Trf molecules, across two technical repeats. All peaks were assigned relative to the reference origami peak (see Section II Methods no. 14).*
***e, Log-likelihood analysis of binding models.*** *The fitting comparison is shown in which black and red symbols indicate the converged value of the log-likelihood function that was minimised during the fit for the four-peaks model and two-peaks model, respectively. The likelihood function was taken as a Poisson distribution. Black curves show the interpolation through the scattered symbols and serve as a guide to the eye.*
***f, Technical replicates underlying the bilayer histogram.*** *Mass histograms from two independent SLB recordings (X1, X2) that were used to generate the combined distributions shown in panel d of the SLB measurement.*

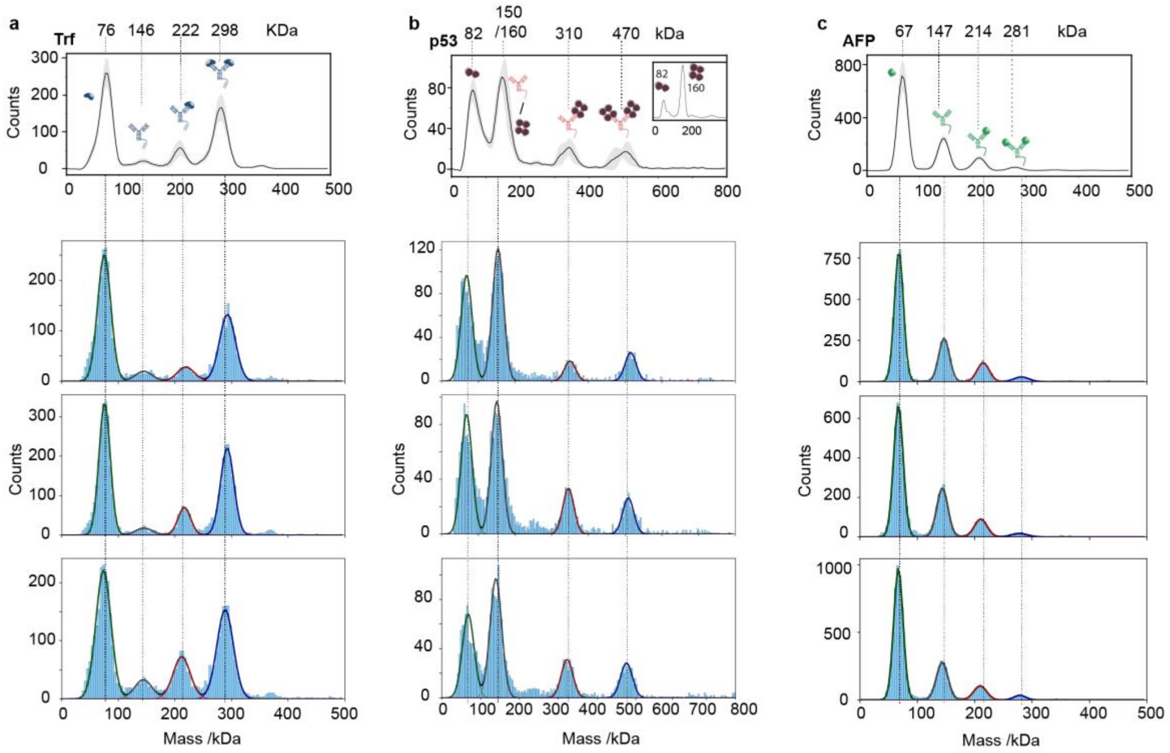

***Fig. S2: Replicate solution landing-assay measurements for protein–antibody mixtures.***
***a–c***, *Technical replicates of solution-phase landing assays for transferrin (Trf) (a), p53 (b), and alpha-fetoprotein (AFP) (c), performed under identical conditions: Trf at 1.5:1 Trf:Trf-IgG, p53 at 5:1 p53:p53-IgG, and AFP at 3:1 AFP:AFP-IgG. For each target, three independent replicates are shown. Histograms display detected particle masses with constrained Gaussian fits corresponding to unbound and successively bound states; dashed lines indicate expected masses. Data correspond to Fig. 2 (a (Top), e, and h).*

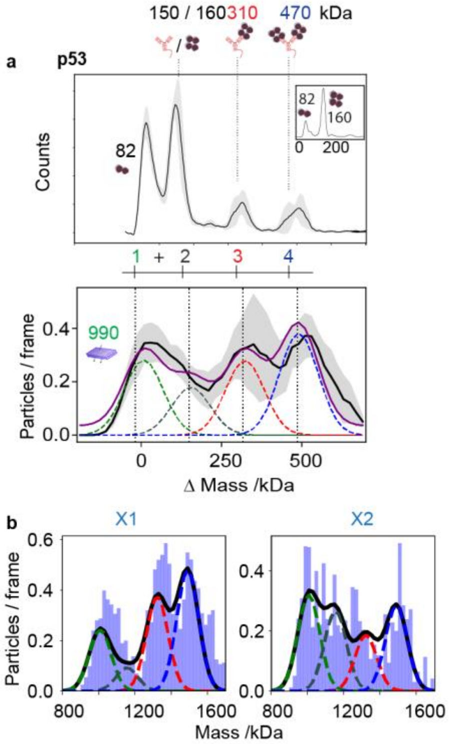

***Fig. S3: Sensor modularity and Detection of oligomeric p53.***

***a, Solution-phase calibration and SLB-based p53 detection. Top:*** *Reference MP landing assay of recombinant p53 resolving the dimer (82 kDa) and tetramer (160 kDa) populations (inset). Upon mixing with the DNA–mAb conjugate (146 kDa), new species corresponding to single-bound (310 kDa) and double-bound (470 kDa) tetramers appear, shown across three technical repeats.*

***Bottom:*** *Mass distributions of SLB-tethered DN functionalised with the p53-specific DNA–mAb conjugate. Gaussian components represent the unmodified origami (green), the antibody-bound origami (grey), and origami bound to one (red) or two (blue) p53 tetramers (data correspond to Fig. 2 e-g)**.***

***b, Technical replicates.*** *Mass histograms from two independent SLB measurements (X1, X2), that were used to generate the combined SLB distribution shown in panel a (bottom), each resolving the same four species—unmodified origami, antibody-bound origami, and one- and two-tetramer-bound complexes.*

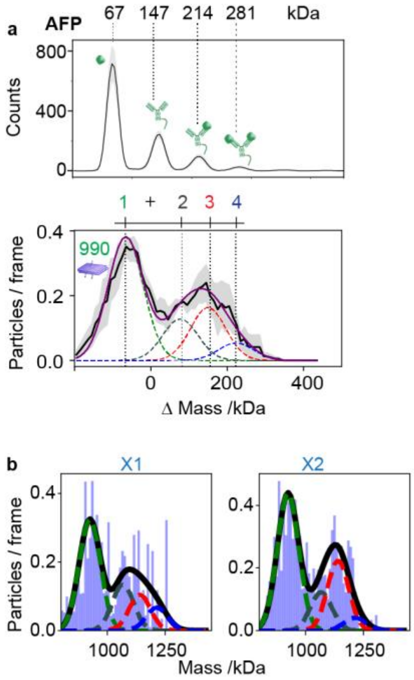

***Fig. S4: Sensor modularity and Detection of AFP.***

***a, Solution-phase calibration and SLB-based AFP detection. Top:*** *Reference landing assay resolving free AFP (AFP, 67 kDa), unbound DNA–mAb conjugate (147 kDa), and AFP-bound antibody conjugates carrying one (214 kDa) or two (281 kDa) proteins, at protein-to-antibody ratios (3:1 AFP:mAb),across three technical repeats (shown in Fig. S2).*

***Bottom:*** *Mass distributions of SLB-tethered DN functionalised with the AFP-specific DNA–mAb conjugate. Gaussian components represent the unmodified origami (green), the antibody-bound origami (grey), and origami bound to one (red) or two (blue) AFP (data correspond to Fig.2 h-j)*.

***b, Technical replicates.*** *Mass histograms from two independent SLB measurements (X1, X2), that were used to generate the combined SLB distribution shown in panel a (bottom), each resolving the same four species—unmodified origami, antibody-bound origami, and one- and two-AFP-bound complexes.*

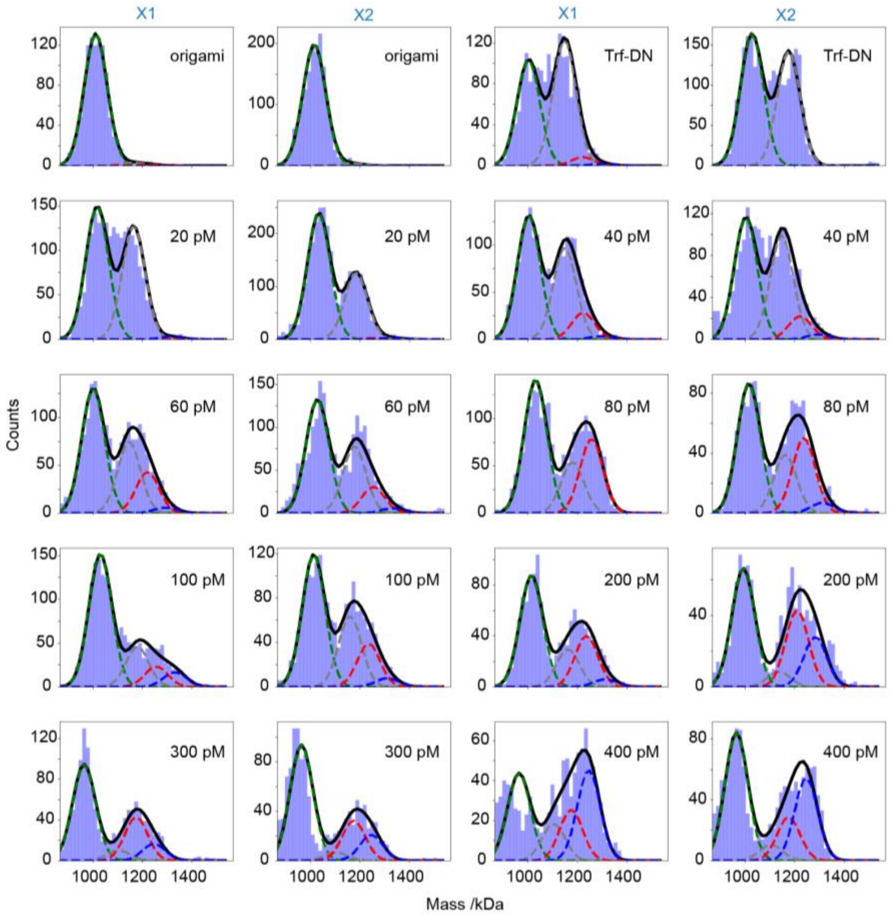

***Fig. S5: SLB-based titration of Trf.***

*Mass histograms from SLB-tethered Trf-DN across a Trf concentration series, shown for two independent measurements (**X1** and **X2**). Top row shows control measurements of unfunctionalised origami and Trf-functionalised origami in the absence of protein. Subsequent rows show increasing Trf concentrations (20–400 pM). Histograms are fitted with constrained Gaussian components corresponding to unbound origami (green), antibody-bound origami (grey), and origami bound to one (red) or two (blue) Trf proteins. Data correspond to Fig. 2a–c.*

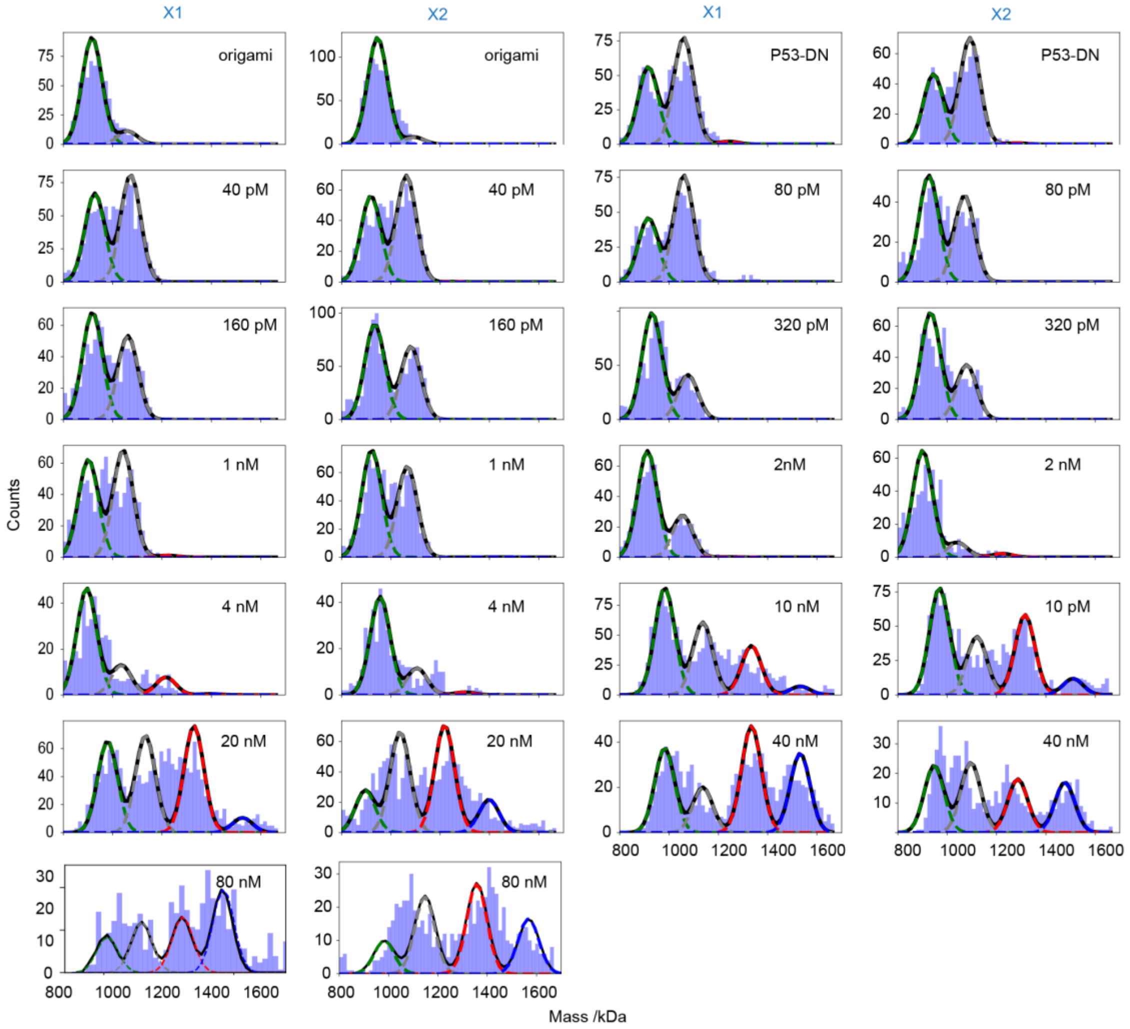

***Fig. S6: SLB-based titration of p53.***

*Mass histograms from SLB-tethered p53-DN across a p53 concentration series, shown for two independent measurements (X1 and X2). The top row shows control measurements of unfunctionalised origami and p53-functionalised origami in the absence of protein. Subsequent rows show increasing p53 concentrations (40 pM–80 nM). Histograms are fitted with constrained Gaussian components corresponding to unbound origami (green), antibody-bound origami (grey), and origami bound to one (red) or two (blue) p53 tetramers. Data correspond to Fig. 2e–g.*

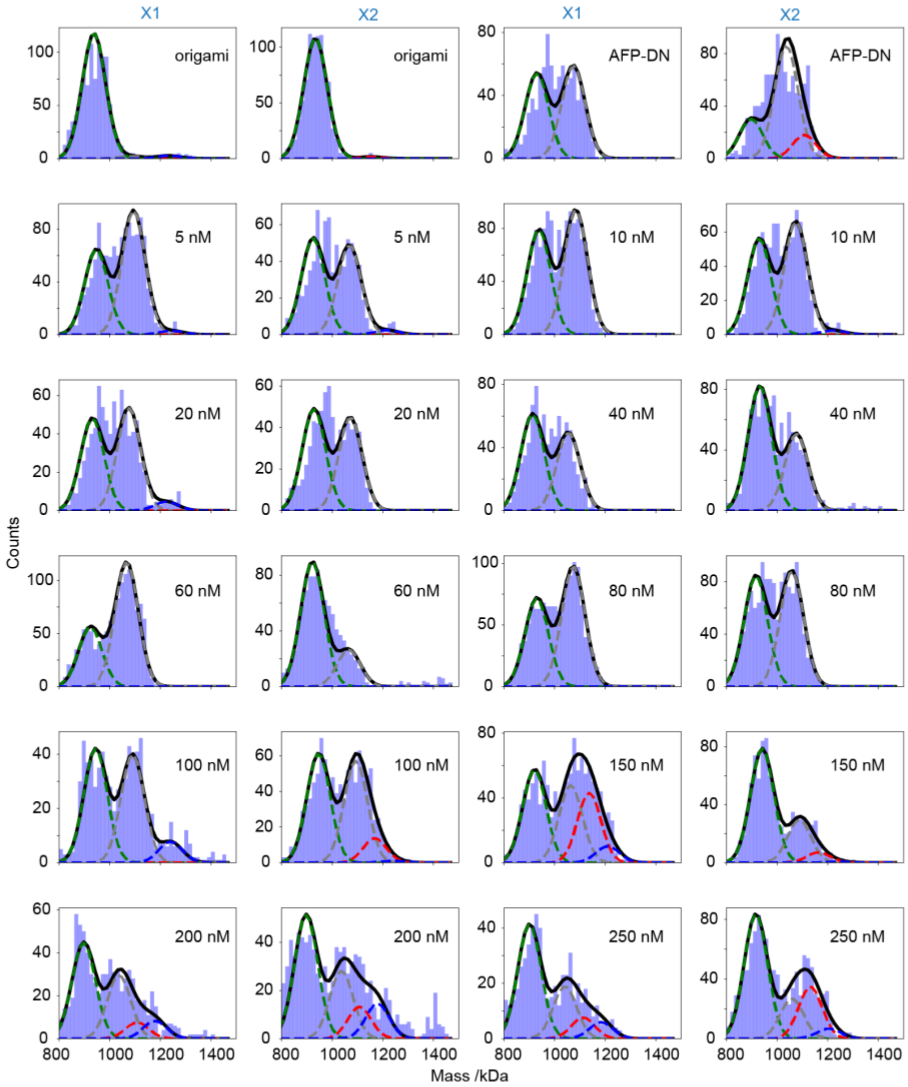

***Fig. S7: SLB-based titration of AFP.***

*Mass histograms from SLB-tethered AFP-DN across an AFP concentration series, shown for two independent measurements (X1 and X2). The top row shows control measurements of unfunctionalised origami and AFP-functionalised origami in the absence of protein. Subsequent rows show increasing AFP concentrations (5–250 nM). Histograms are fitted with constrained Gaussian components corresponding to unbound origami (green), antibody-bound origami (grey), and origami bound to one (red) or two (blue) AFP proteins. Data correspond to Fig. 2h–j.*

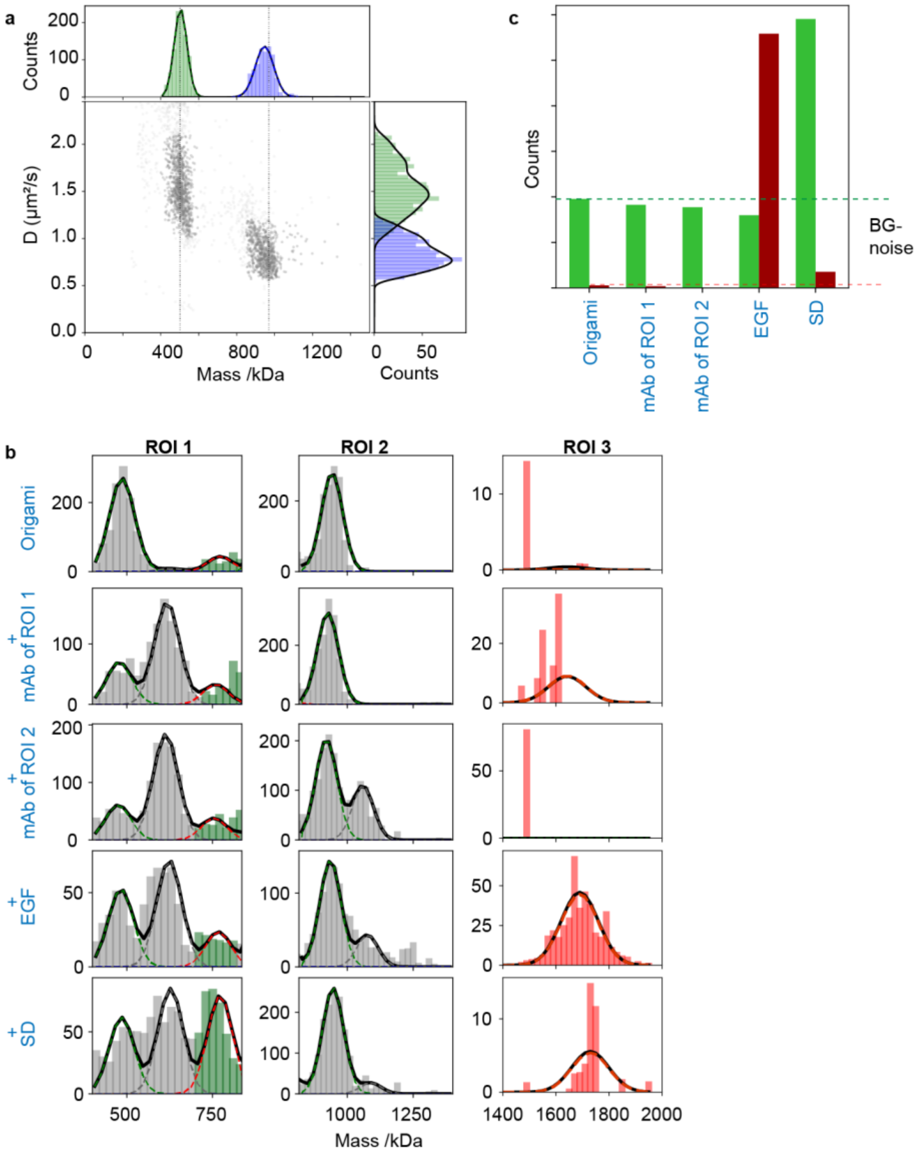

***Fig. S8: Mass–diffusion–resolved detection of EGF and programmable nanosensor resetting (data correspond to Fig. 3).***

***a, Mass–diffusion identification of the two encoded nanosensors.***

*Two-dimensional mass–diffusion plot showing the two encoded DNA-origami structures that form the basis of the DNs used in the EGF detection assay (see Fig. S11 for details of scaffold digestion). Their distinct mass–diffusion signatures define ROI1 and ROI2, which are different from the detection ROI (ROI3)**,** which emerges only upon EGF addition when the two structures are crosslinked, providing a unique mass–diffusion signature for the crosslinked complex.*

***b, ROI-resolved mass histograms across the five sequential stages of the assay.*** *Mass histograms for ROI1–ROI3 across the five steps of the EGF assay: unmodified structures; addition of the ROI1-specific mAb; addition of the ROI2-specific mAb; addition of 2 nM EGF; and strand displacement (SD).*

*The sequential mAb additions establish that each nanosensor responds only to its intended sequence-specific antibody, producing antibody-bound peaks uniquely in ROI1 or ROI2 and confirming independent programmability. Prior to EGF addition, ROI3 shows negligible signal. Upon addition of EGF, a new peak emerges in ROI3 with the combined mass of both nanosensors and reduced diffusion (as defined in panel a), consistent with EGF-induced crosslinking. SD, designed to remove the ROI2 antibody, eliminates the crosslinked species in ROI3 and generates the expected singly tethered SD product in ROI1. Gaussian components represent unmodified structures (green), antibody-bound structures (grey), and the EGF-crosslinked complex (red).*

***c, Quantification of crosslinked and strand-displaced species.*** *Quantitative analysis of the crosslinked species (red peak) and the SD product (green peak) across the pre-EGF, post-EGF, and post-SD stages confirms the fidelity of EGF detection and demonstrates programmable topological switching and signal reconfiguration.*

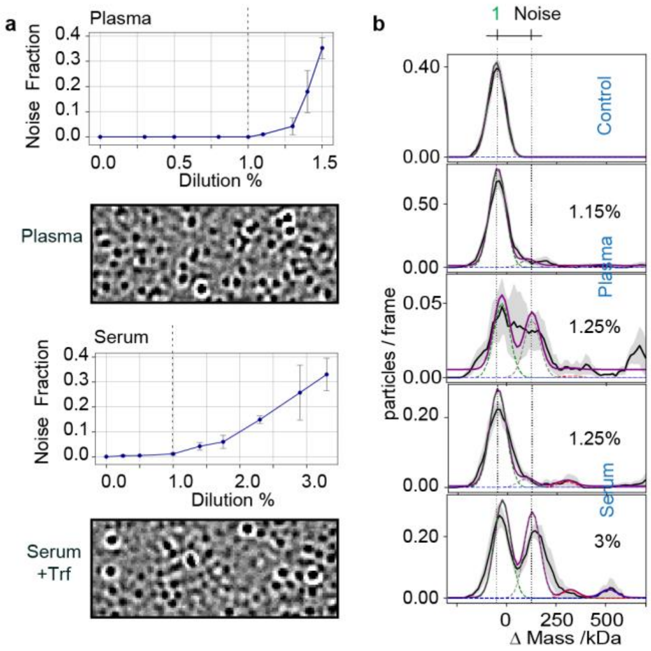

***Fig. S9: Definition and quantification of noise fraction in serum and plasma.***
***a,*** *Noise fraction extracted from bare (non-functionalised) origami as a function of serum and plasma dilution on supported lipid bilayers (SLBs), quantified as the fitted mass contribution above the origami peak. Dilution series were performed stepwise on the same bilayer for each matrix (serum and plasma in separate experiments). Representative MP images are shown below each plot.*
***b,*** *Representative Δ-mass distributions illustrating the definition of the noise fraction. Control measurements (top) show only the origami peak, while increasing serum or plasma content introduces additional mass contributions attributed to non-specific protein adsorption ("noise"). Dashed vertical lines indicate the origami reference peak; fitted components correspond to origami and noise contributions. Data correspond to figure 4a.*

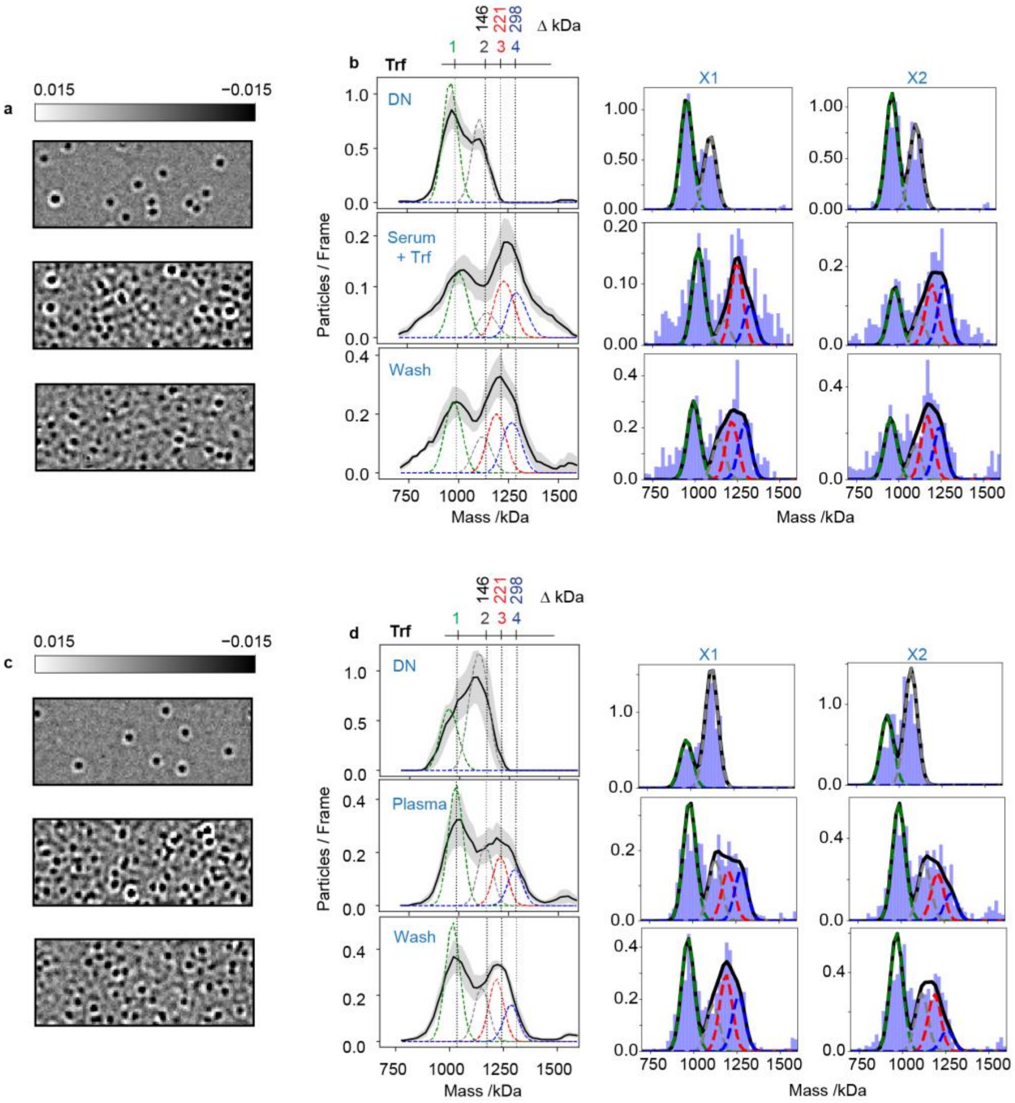

***Fig. S10: Evaluation of DNA-nanosensor performance in complex media (data correspond to Fig. 4).***

***a, Dynamic MP images for serum measurements.*** *Representative MP images of DNs in buffer (reference), after addition of 1% serum spiked with 200 pM Trf, and after washing.*

***b, Mass histograms for serum measurements and corresponding technical replicates.*** *Left: Mass histograms of DN in buffer, in 1% serum + 200 pM Trf, and after wash. Gaussian components represent unmodified DN (green), antibody-bound DN (grey), and DN carrying one (red) or two (blue) Trf molecules. Trf-dependent peaks remain detectable in 1% serum. Right: corresponding technical replicates (X1, X2) for the 1% serum + Trf condition.*

***c, Dynamic MP images for plasma measurements.*** *Representative dynamic MP frames of SLB-tethered DNA nanostructures recorded in buffer (reference), after addition of 1% plasma, and after washing. No Trf was added; detection arises from native plasma transferrin.*

***d, Mass histograms for plasma measurements and corresponding technical replicates.*** *Left: Mass histograms of DN in buffer, in 1% plasma, and after wash. Gaussian components represent unmodified DN (green), antibody-bound DN (grey), and DN carrying one (red) or two (blue) Trf molecules originating from endogenous plasma transferrin. Washing removes nonspecific plasma species while preserving Trf-dependent peaks. Right: corresponding technical replicates (X1, X2) for the 1% plasma condition.*

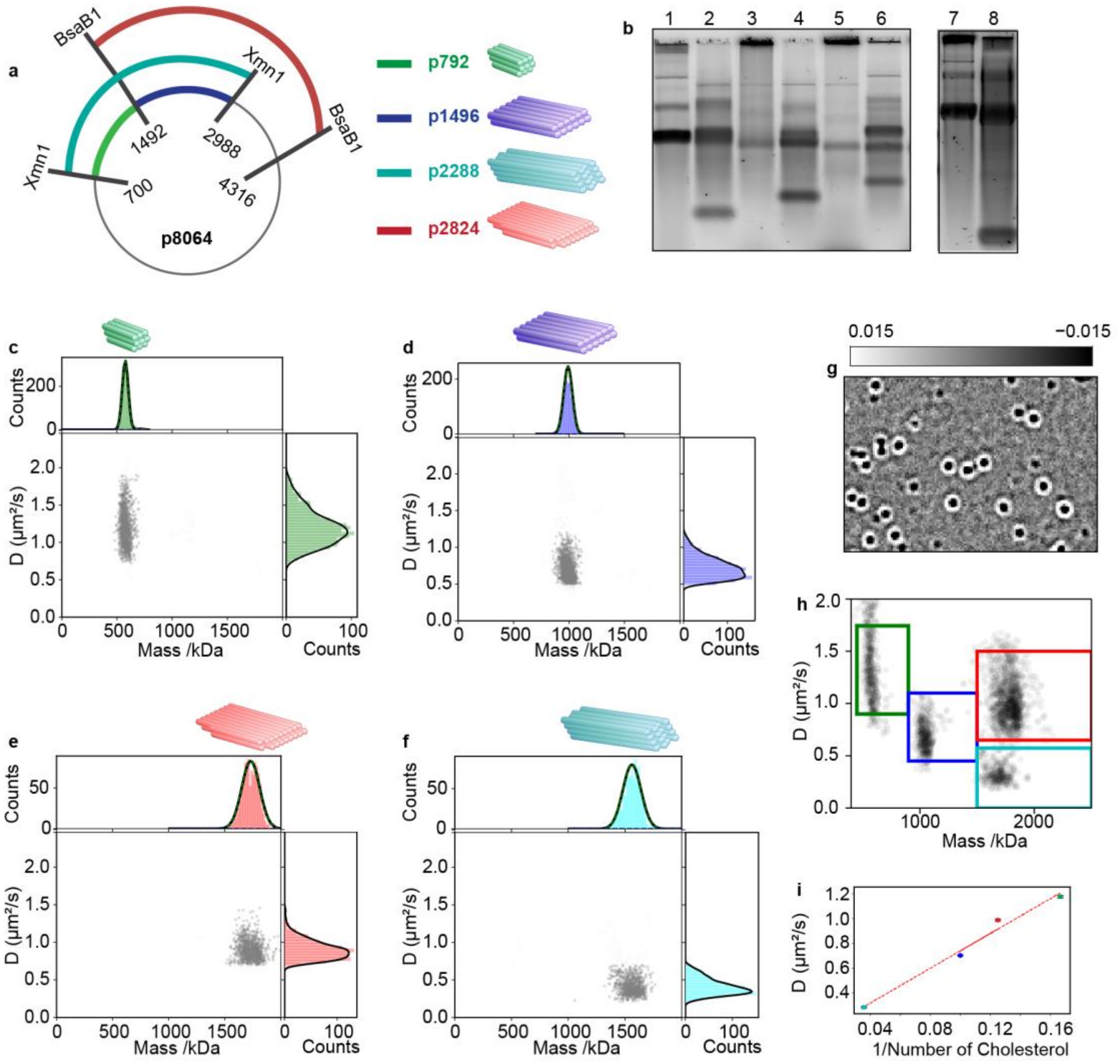

***Fig. S11: Creating a multiplexed panel with defined mass–diffusion signatures.***

***a, Scaffold digestion scheme.*** *Schematic showing the positions of BsaBI and XmnI digestion sites on the p8064 scaffold used to generate four scaffold fragments (p792, p1496, p2288, p2824) for folding the corresponding origami nanostructures.*

***b, Agarose gel validation of scaffold fragments.*** *Lanes 1, 3, 5, and 7 show the p8064 scaffold. Lanes 2, 4, 6, and 8 show the digestion reaction products for the four scaffold fragments, p1496, p2288, p2824, and p792, respectively.*

***c–f, Mass–diffusion signatures of the four origami designs.*** *Two-dimensional mass–diffusion plots for each nanostructure (p792, p1496, p2288, p2824). Each design exhibits a well-defined mass–diffusion signature, enabling mass- and diffusion-based encoding.*

***g, Mixed-panel dynamic MP image.*** *Representative dynamic MP image of a mixed panel of the four origami designs bound to an SLB via cholesterol anchors, illustrating simultaneous detection of the encoded structures.*

***h, ROI definition for the four-structure panel.*** *Combined mass–diffusion plot showing the non-overlapping mass–diffusion clusters for each origami design* ***observed in panel g****. The coloured boxes represent the ROIs used for structure-specific assignment in multiplexed experiments.*

***i, Diffusion vs. cholesterol anchor number.*** *Plot of diffusion coefficient versus the inverse number of cholesterol anchors used for SLB tethering. The observed trend demonstrates controllable tuning of mobility through the number of membrane anchors appended to the origami.*

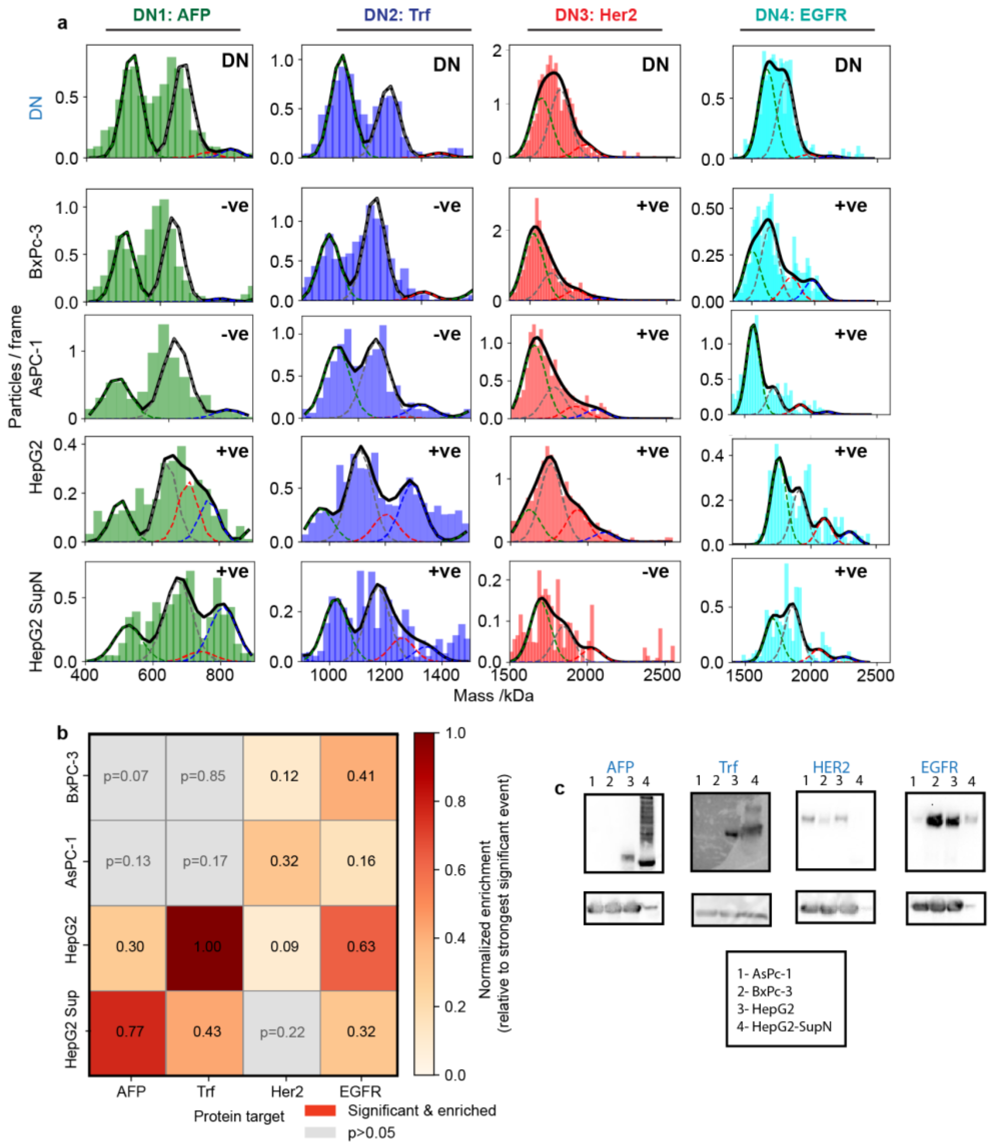

***Fig. S12: Multiplexed detection of protein targets across diverse cancer cell lysates (data correspond to Fig. 5b).***

***a, ROI-resolved mass histograms for four DNA nanosensors (DNs) across four cell lysates.*** *Mass histograms from combined results of three replicates for the four mass- and diffusion-encoded nanosensors DN1–DN4, each functionalised for a distinct protein target (DN1: AFP, DN2: Trf, DN3: Her2, DN4: EGFR), measured on SLBs exposed to lysates from BxPC-3, AsPC-1, HepG2, and HepG2-SupN cells. For each nanosensor, Gaussian components represent unmodified DN (black), antibody-bound DN (grey), and specific target-bound complexes (DN carrying one (red) or two (blue) target molecules). "+ve" and "–ve" annotations indicate the presence or absence of a detectable target-specific peak above background that showed statistical significance across the three replicates. These measurements show target-specific binding patterns across the four lysate conditions within a single multiplexed experiment.*

***b, Normalised enrichment heat map.*** *Quantification of target-specific binding events for each nanosensor–lysate pair shown in **panel a**. Enrichment values are normalised to the strongest significant event in the panel, and coloured-squares denote statistically significant enrichment ($p < 0.05$). Grey squares correspond to "–ve" results, in which no statistically significant enrichment was obtained. The heat map highlights distinct proteomic signatures across the four cancer cell lysates.*

***c, Western blot validation.*** *Western blots for AFP, Her2, EGFR, and Trf across the same four lysates (1: AsPC-1; 2: BxPC-3; 3: HepG2; 4: HepG2-SupN). The observed band intensities closely corroborate the nanosensor-derived enrichment profiles shown in panels **a** and **b**.*

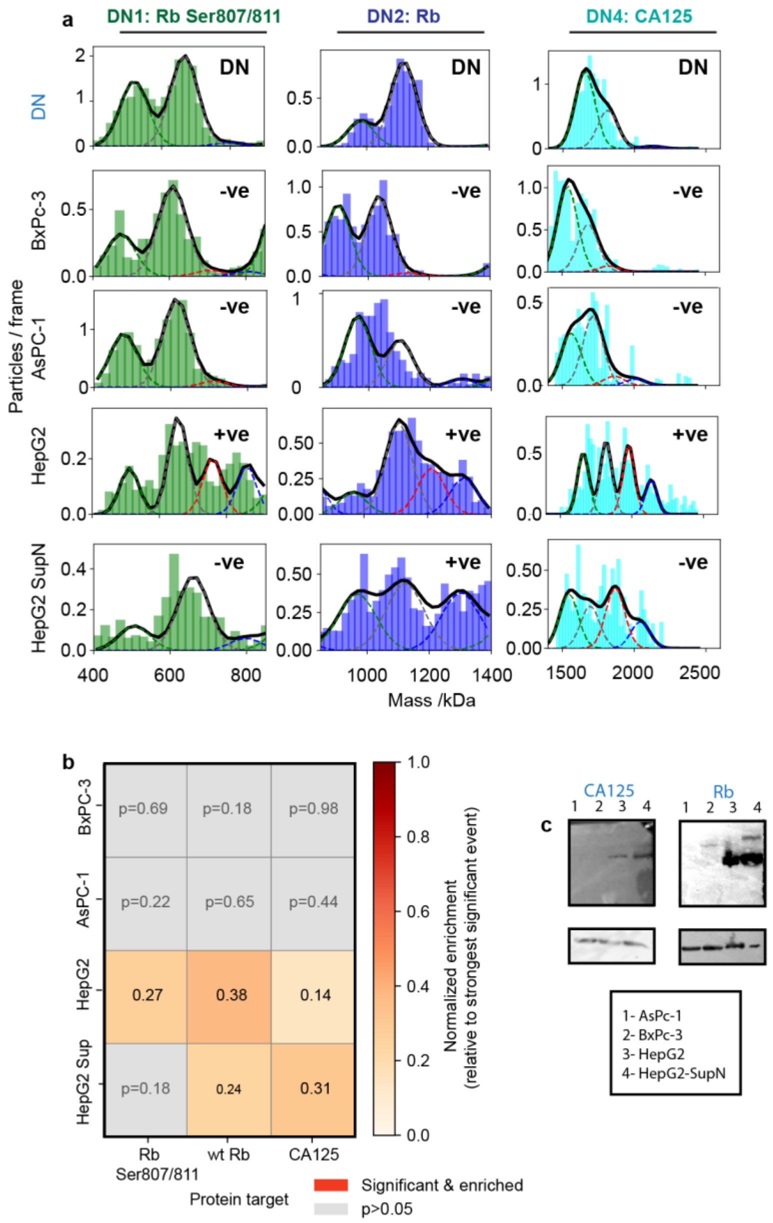

***Fig. S13: Multiplexed detection of PTM & CA125 across diverse cancer cell lysates (data correspond to Fig. 5b).***

***a, ROI-resolved mass histograms for three DNA nanosensors (DNs) across four cell lysates.*** *Mass histograms from combined results of three replicates for the three mass- and diffusion-encoded nanosensors DN1–DN3, each functionalised for a distinct protein target (DN1: phosphorylated Rb Ser807/811, DN2: total Rb, DN3: CA125), measured on SLBs exposed to lysates from BxPC-3, AsPC-1, HepG2, and HepG2-SupN cells. For each nanosensor, Gaussian components represent unmodified DN (black), antibody-bound DN (grey), and specific target-bound complexes (DN carrying one (red) or two (blue) target molecules). "+ve" and "–ve" annotations indicate the presence or absence of a detectable target-specific peak above background that showed statistical significance across the three replicates.*

***b, Normalised enrichment heat map.*** *Quantification of target-specific binding events for each nanosensor–lysate pair shown in **panel a**. Enrichment values are normalised to the strongest significant event in the panel, and coloured-squares denote statistically significant enrichment ($p < 0.05$). Grey squares correspond to "–ve" results, in which no statistically significant enrichment was obtained. The heat map highlights distinct proteomic signatures across the four cancer cell lysates.*

***c, Western blot validation.*** *Western blots for CA125 and Rb (phosphorylated and total) across the same four lysates (1: AsPC-1; 2: BxPC-3; 3: HepG2; 4: HepG2-SupN). The observed band intensities closely corroborate the nanosensor-derived enrichment profiles shown in panels **a** and **b**.*

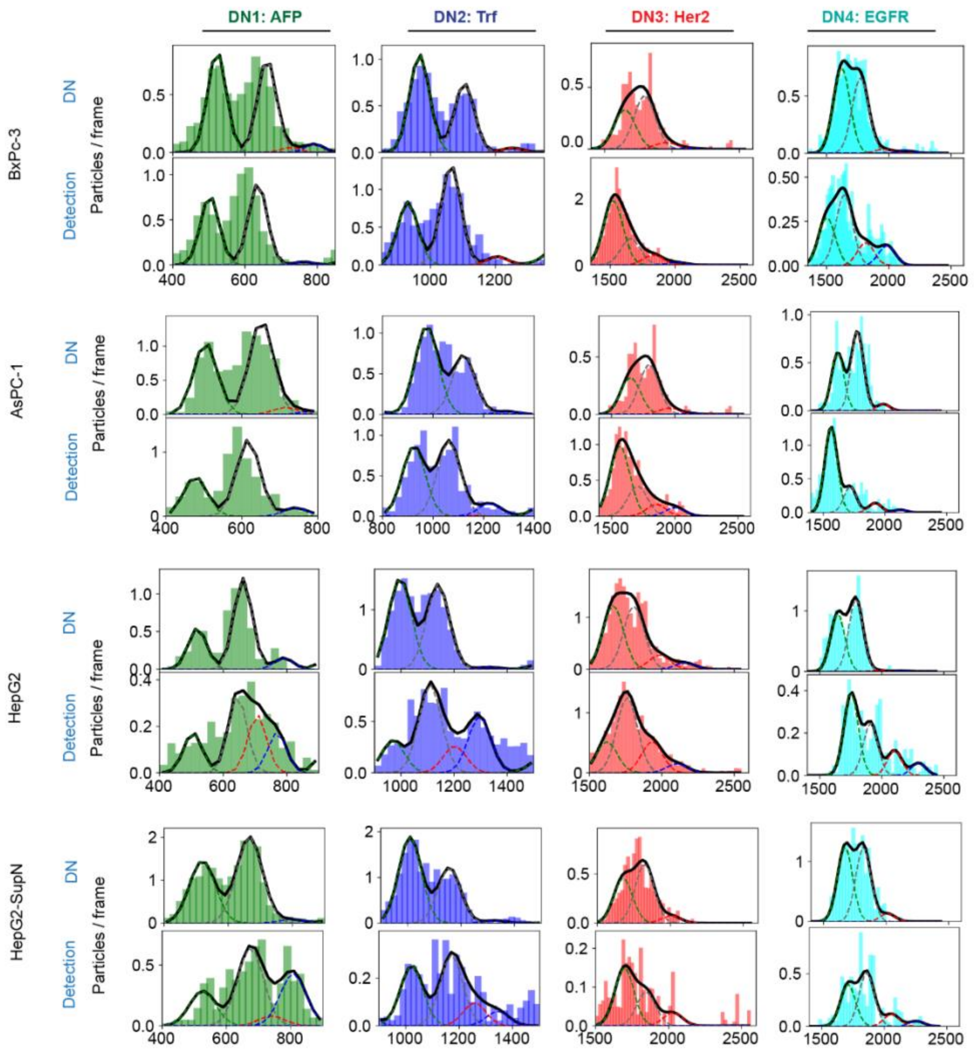

***Fig. S14: DN characterisation and corresponding detection profiles across four cancer cell lysates (combined three replicates; data correspond to Fig. 5b and S12).***

*For each nanosensor (DN1: AFP, DN2: Trf, DN3: Her2, DN4: EGFR) and each lysate (BxPC-3, AsPC-1, HepG2, HepG2-SupN), the* ***top row*** *shows the DN-only mass distributions obtained from the combined three replicates used in that experiment. The* ***bottom row*** *shows the corresponding detection distributions—also the combined three replicates—measured after exposing the SLB-tethered DNs to the same lysate.*

*Gaussian components represent unmodified DN (black), antibody-bound DN (grey), and target-bound complexes (DN carrying one (red) or two (blue) target molecules).*

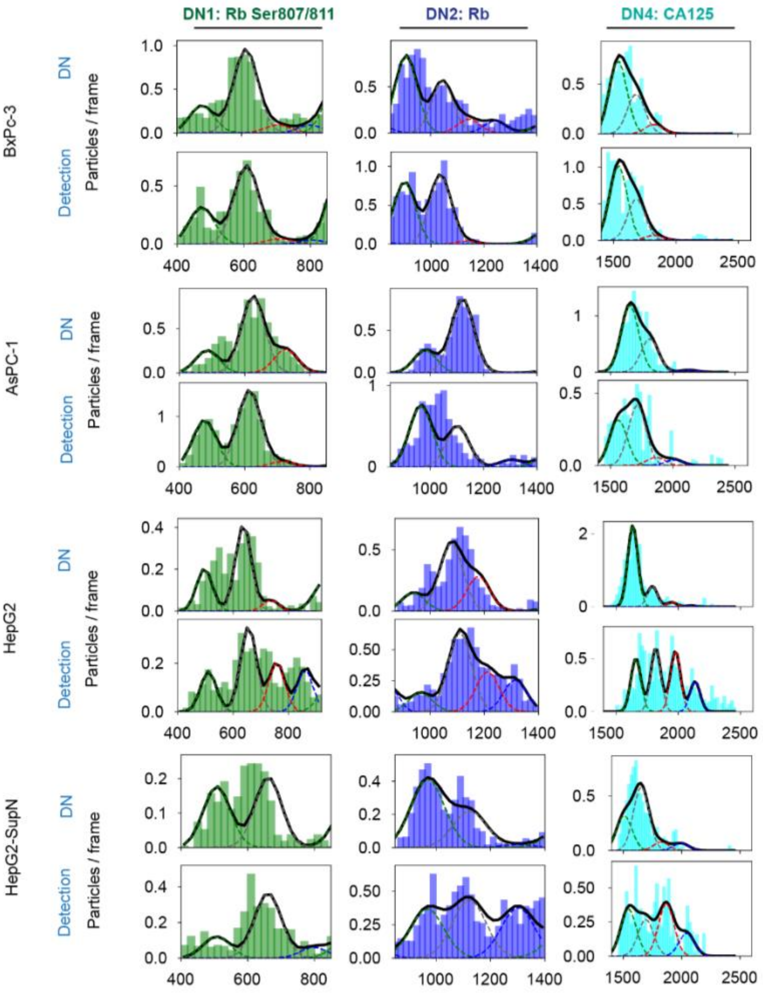

***Fig. S15: DN characterisation and corresponding detection profiles across four cancer cell lysates of PTM and CA125 (combined three replicates; data correspond to Fig. 5b and S13).***

*For each nanosensor (DN1: phosphorylated Rb Ser807/811, DN2: total Rb, DN3: CA125) and each lysate (BxPC-3, AsPC-1, HepG2, HepG2-SupN), the **top row** shows the DN-only mass distributions obtained from the combined three replicates used in that experiment. The **bottom row** shows the corresponding detection distributions—also the combined three replicates—measured after exposing the SLB-tethered DNs to the same lysate. Gaussian components represent unmodified DN (black), antibody-bound DN (grey), and target-bound complexes (DN carrying one (red) or two (blue) target molecules).*

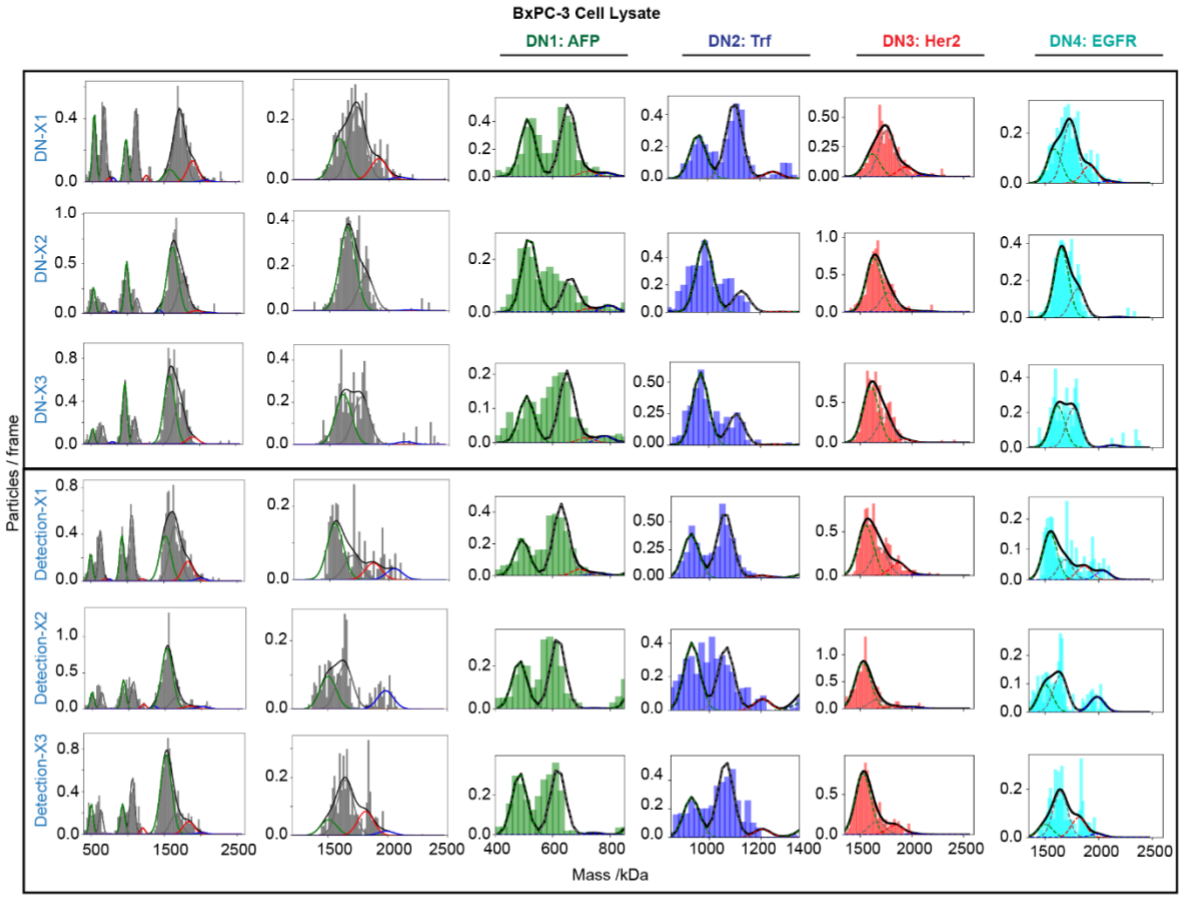

***Fig. S16: Replicate-level DN characterisation and detection in BxPC-3 lysate (data correspond to Fig. 5b and Fig. S12, S14).***

*Representative mass histograms for all three replicates (DN-X1, DN-X2, DN-X3) for the four mass- and diffusion-encoded nanosensors (DN1: AFP, DN2: Trf, DN3: Her2, DN4: EGFR) measured in BxPC-3 cell lysate. For each nanosensor, the top three rows show the DN-only measurements and the bottom three rows show the corresponding detection measurements performed on the same DN populations after exposure to lysate.*

*Grey bars show the measured mass distributions of individual replicates. Overlaid curves show the Gaussian components used for ROI-based quantification: unmodified DN (black), antibody-bound DN (grey), DN carrying one target molecule (red), and DN carrying two target molecules (blue).*

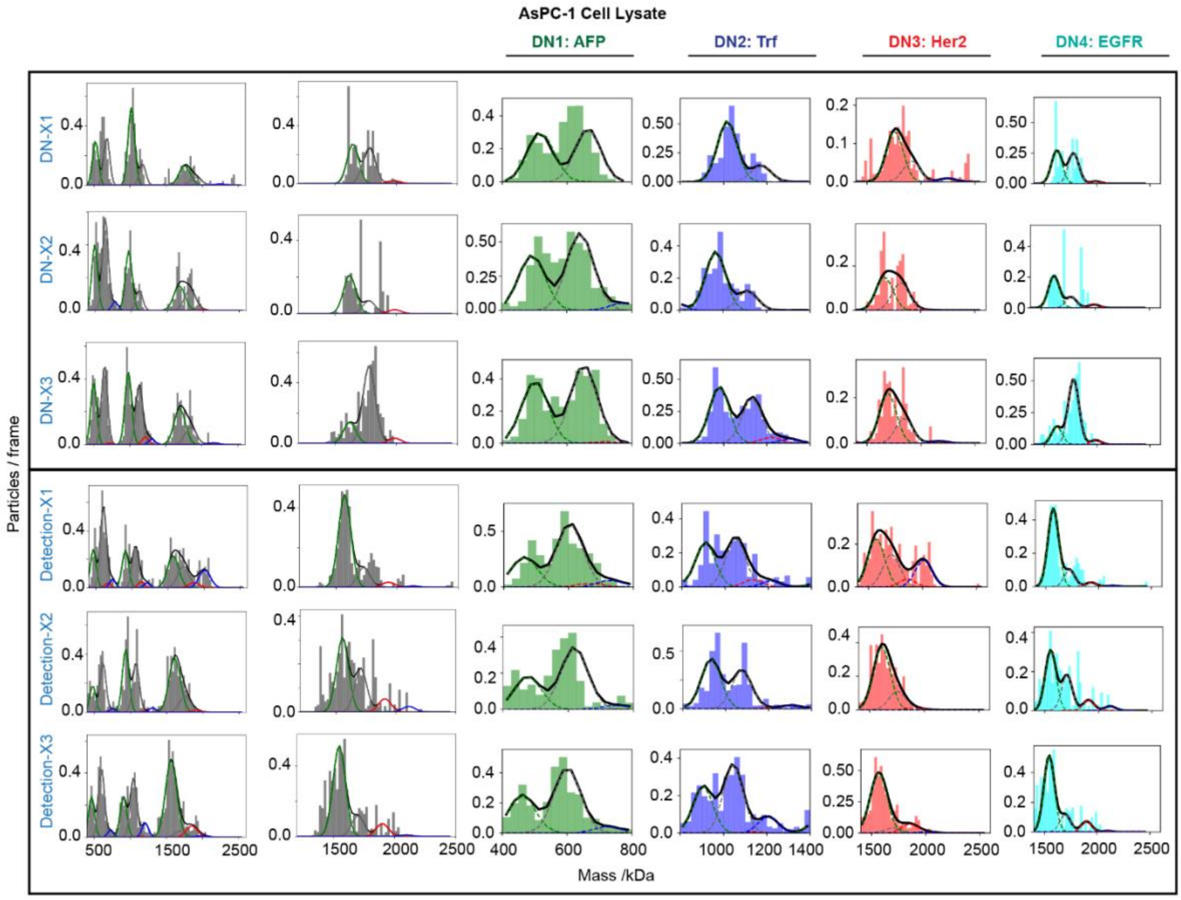

***Fig. S17: Replicate-level DN characterisation and detection in AsPC-1 lysate (data correspond to Fig. 5b and Fig. S12, S14).***

*Representative mass histograms for all three replicates (DN-X1, DN-X2, DN-X3) for the four mass- and diffusion-encoded nanosensors (DN1: AFP, DN2: Trf, DN3: Her2, DN4: EGFR) measured in AsPc-1 cell lysate. For each nanosensor, the top three rows show the DN-only measurements and the bottom three rows show the corresponding detection measurements performed on the same DN populations after exposure to lysate.*

*Grey bars show the measured mass distributions of individual replicates. Overlaid curves show the Gaussian components used for ROI-based quantification: unmodified DN (black), antibody-bound DN (grey), DN carrying one target molecule (red), and DN carrying two target molecules (blue).*

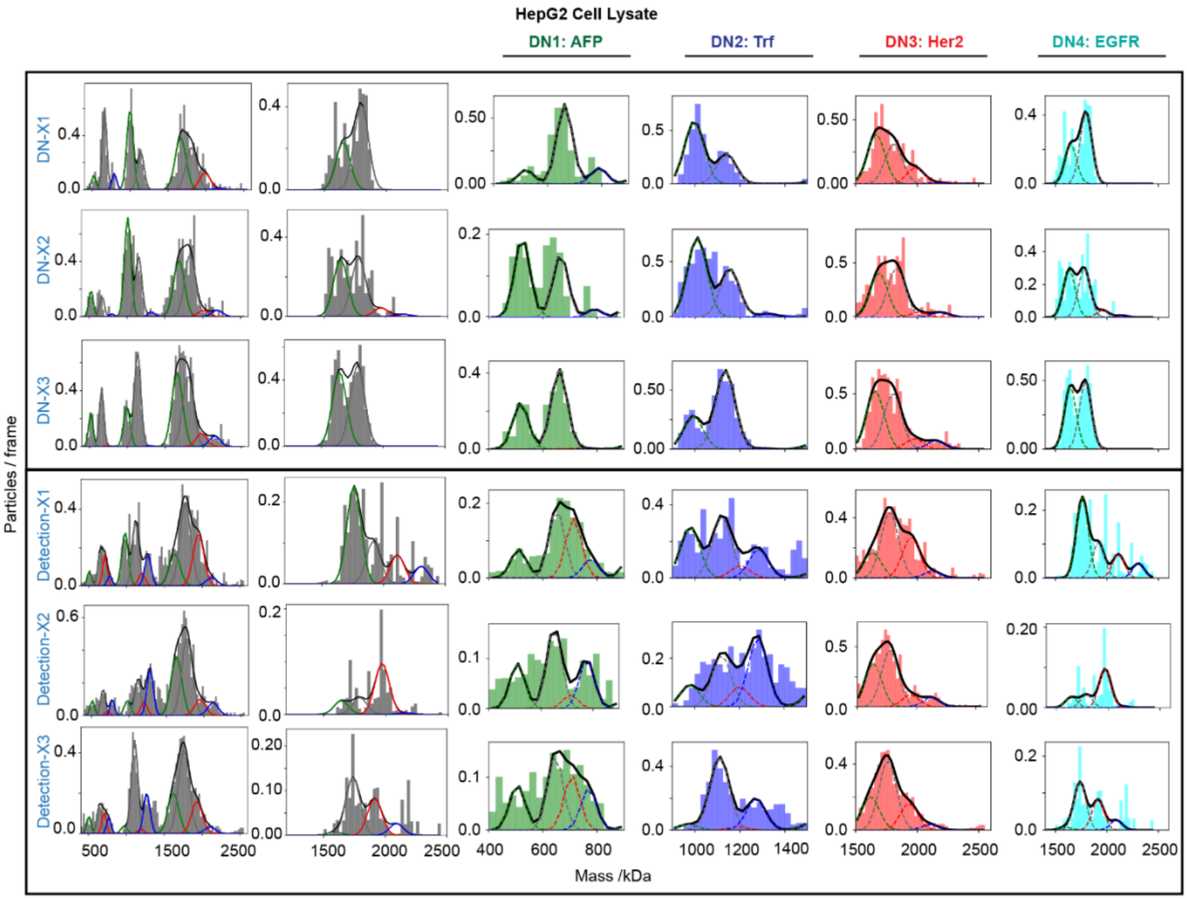

***Fig. S18: Replicate-level DN characterisation and detection in HepG2 lysate (data correspond to Fig. 5b and Fig. S12, S14).***

*Representative mass histograms for all three replicates (DN-X1, DN-X2, DN-X3) for the four mass- and diffusion-encoded nanosensors (DN1: AFP, DN2: Trf, DN3: Her2, DN4: EGFR) measured in HepG2 cell lysate. For each nanosensor, the top three rows show the DN-only measurements and the bottom three rows show the corresponding detection measurements performed on the same DN populations after exposure to lysate.*

*Grey bars show the measured mass distributions of individual replicates. Overlaid curves show the Gaussian components used for ROI-based quantification: unmodified DN (black), antibody-bound DN (grey), DN carrying one target molecule (red), and DN carrying two target molecules (blue).*

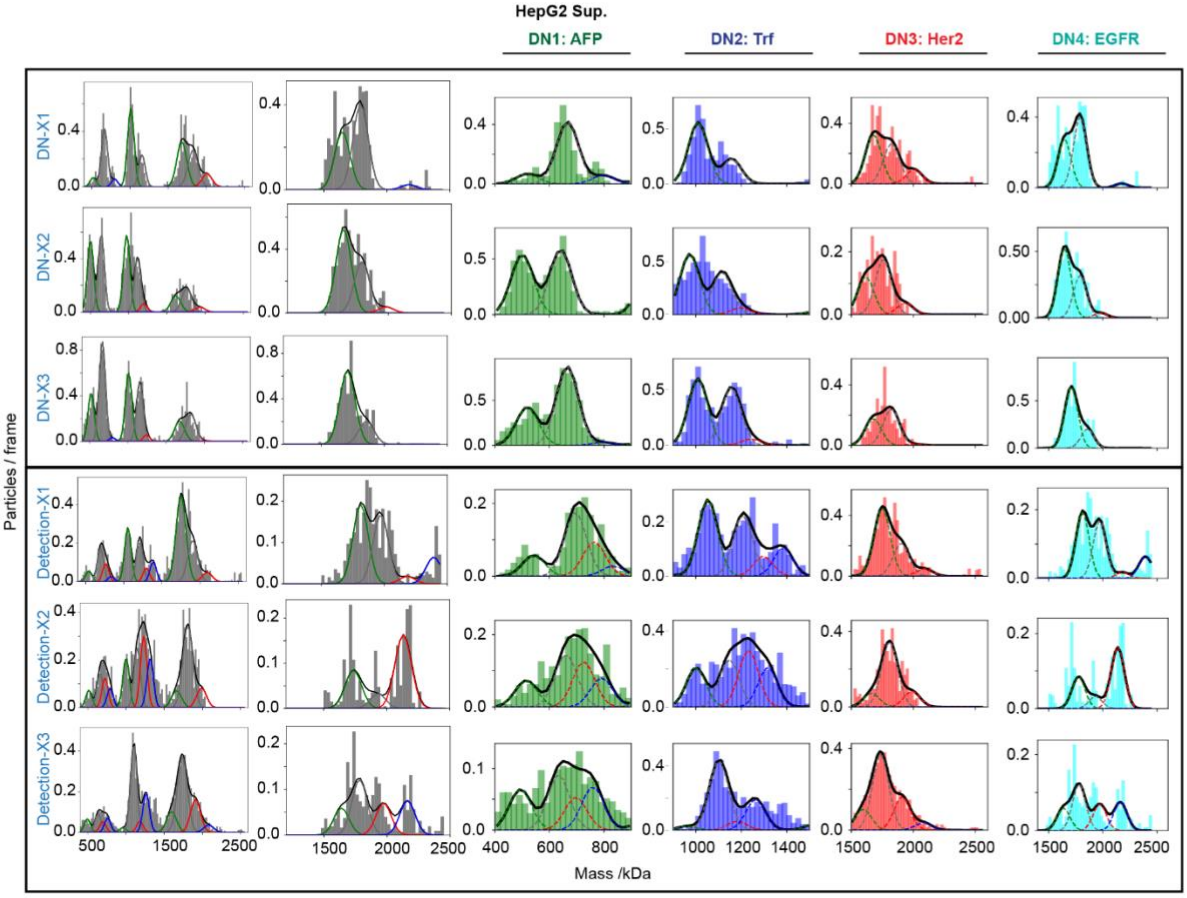

***Fig. S19: Replicate-level DN characterisation and detection in HepG2-SupN (Data correspond to Fig. 5b and Fig. S12, S14).***

*Representative mass histograms for all three replicates (DN-X1, DN-X2, DN-X3) for the four mass- and diffusion-encoded nanosensors (DN1: AFP, DN2: Trf, DN3: Her2, DN4: EGFR) measured in HepG2-SupN. For each nanosensor, the top three rows show the DN-only measurements and the bottom three rows show the corresponding detection measurements performed on the same DN populations after exposure to lysate.*

*Grey bars show the measured mass distributions of individual replicates. Overlaid curves show the Gaussian components used for ROI-based quantification: unmodified DN (black), antibody-bound DN (grey), DN carrying one target molecule (red), and DN carrying two target molecules (blue).*

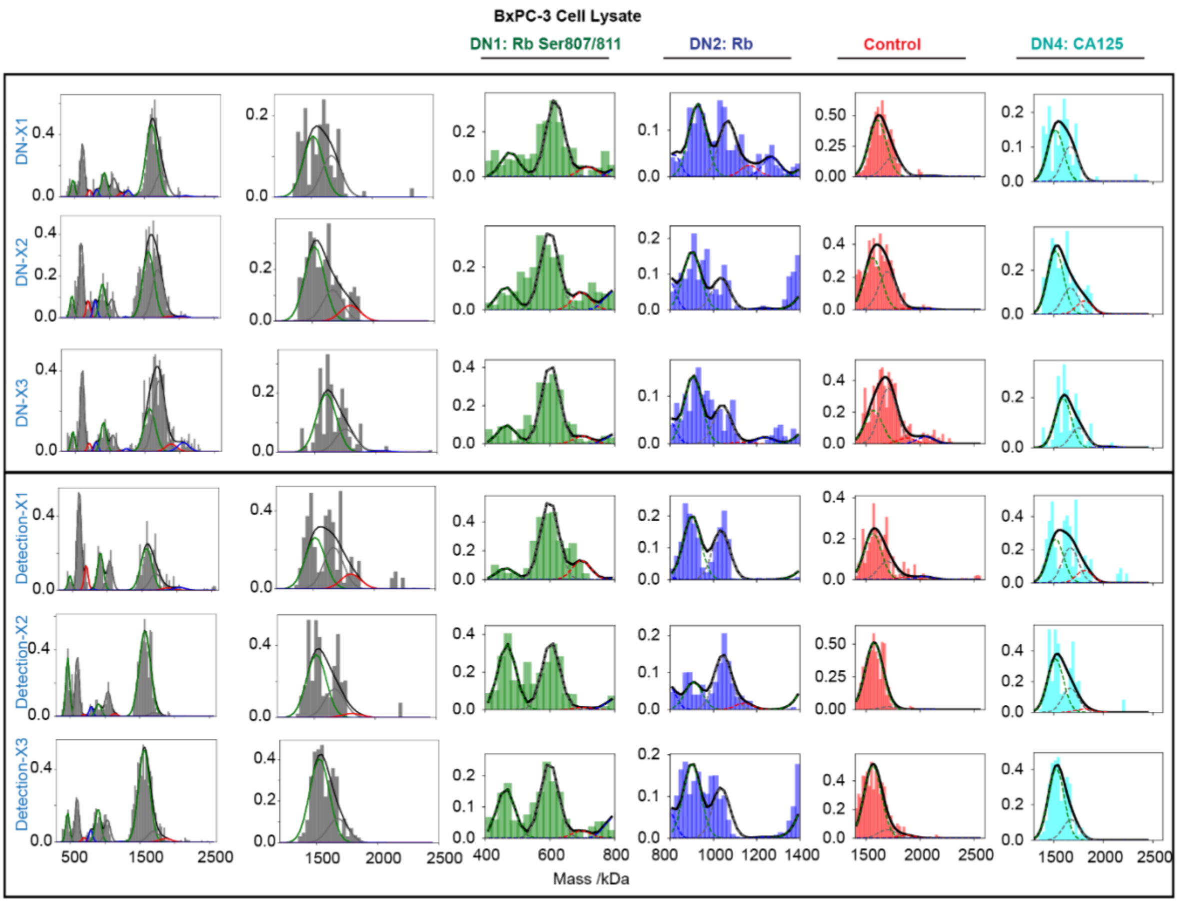

***Fig. S20: Replicate-level DN characterisation and detection for PTM-resolved nanosensors & CA125 in BxPC-3 lysate (Data correspond to Fig. 5b and Fig. S13, S15)***

*Representative mass histograms for all three replicates (DN-X1, DN-X2, DN-X3) for the four mass- and diffusion-encoded nanosensors (DN1: phosphorylated Rb Ser807/811, DN2: total Rb, DN3: control origami without antibody, DN4: CA125) measured in BxPC-3 cell lysate. For each nanosensor, the top three rows show the DN-only measurements and the bottom three rows show the corresponding detection measurements performed on the same DN populations after exposure to lysate.*

*Grey bars show the measured mass distributions of individual replicates. Overlaid curves show the Gaussian components used for ROI-based quantification: unmodified DN (black), antibody-bound DN (grey), DN carrying one target molecule (red), and DN carrying two target molecules (blue). DN3 (control) contains **no antibody** and therefore reports only the intrinsic background of the system; occasional high-mass species in this control channel represent rare background noise. In downstream enrichment analysis, only peaks exceeding this control-defined noise floor are considered target-specific.*

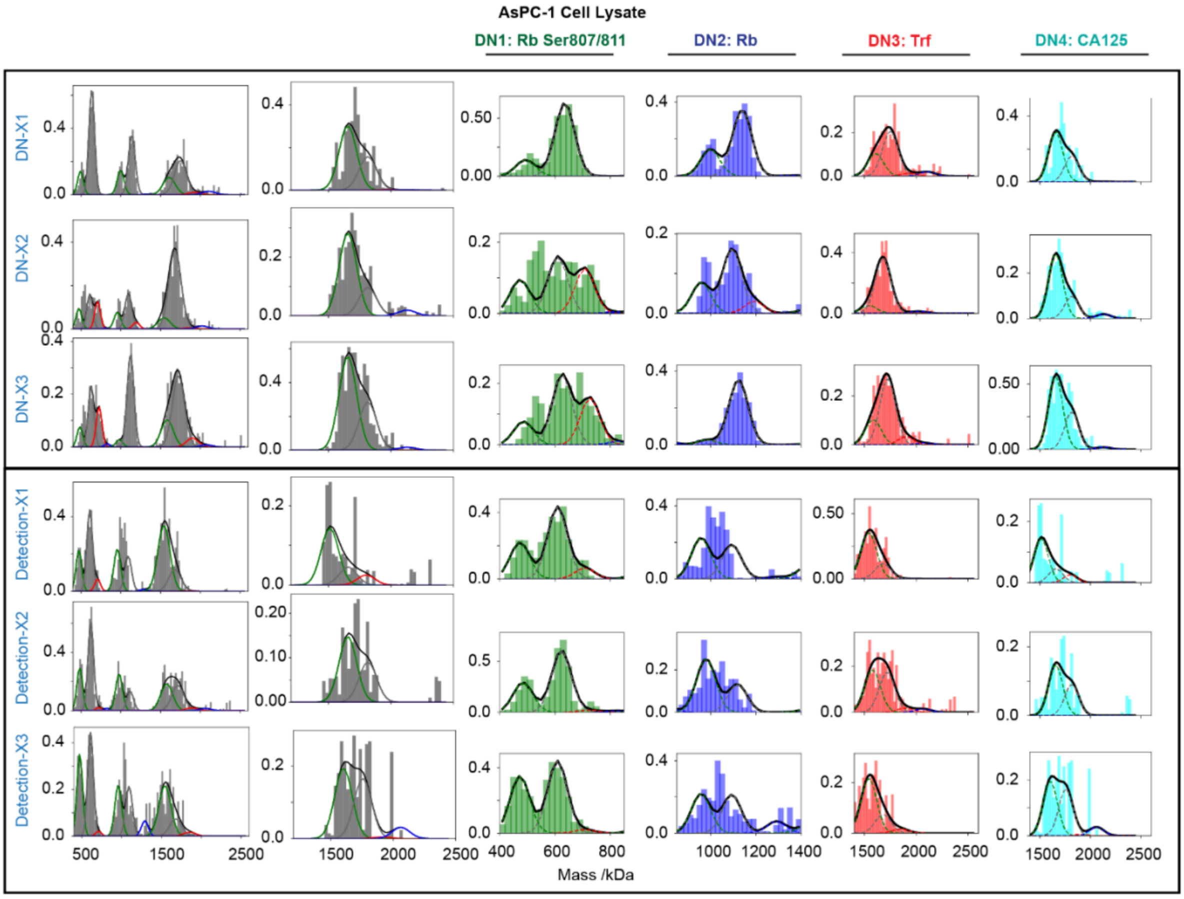

***Fig. S21: Replicate-level DN characterisation and detection for PTM-resolved nanosensors & CA125 in AsPC-1 lysate (data correspond to Fig. 5b and Fig. S13, S15).***

*Representative mass histograms for all three replicates (DN-X1, DN-X2, DN-X3) for the four mass- and diffusion-encoded nanosensors (DN1: phosphorylated Rb Ser807/811, DN2: total Rb, DN3: Trf, DN4: CA125) measured in* ***AsPC-1*** *cell lysate. For each nanosensor, the top three rows show the DN-only measurements and the bottom three rows show the corresponding detection measurements performed on the same DN populations after exposure to lysate.*

*Grey bars show the measured mass distributions of individual replicates. Overlaid curves show the Gaussian components used for ROI-based quantification: unmodified DN (black), antibody-bound DN (grey), DN carrying one target molecule (red), and DN carrying two target molecules (blue). Importantly, DN3 (Trf) corresponds to a different origami design from the Trf-specific DN used in the canonical-protein multiplexing experiment (Fig. S10–S13), yet it again shows no Trf-specific peak in AsPC-1 lysate. This confirms both the absence of detectable Trf in this lysate and the robustness of the platform, demonstrating that nanosensor identity can be reassigned to new targets without producing false-positive signals.*

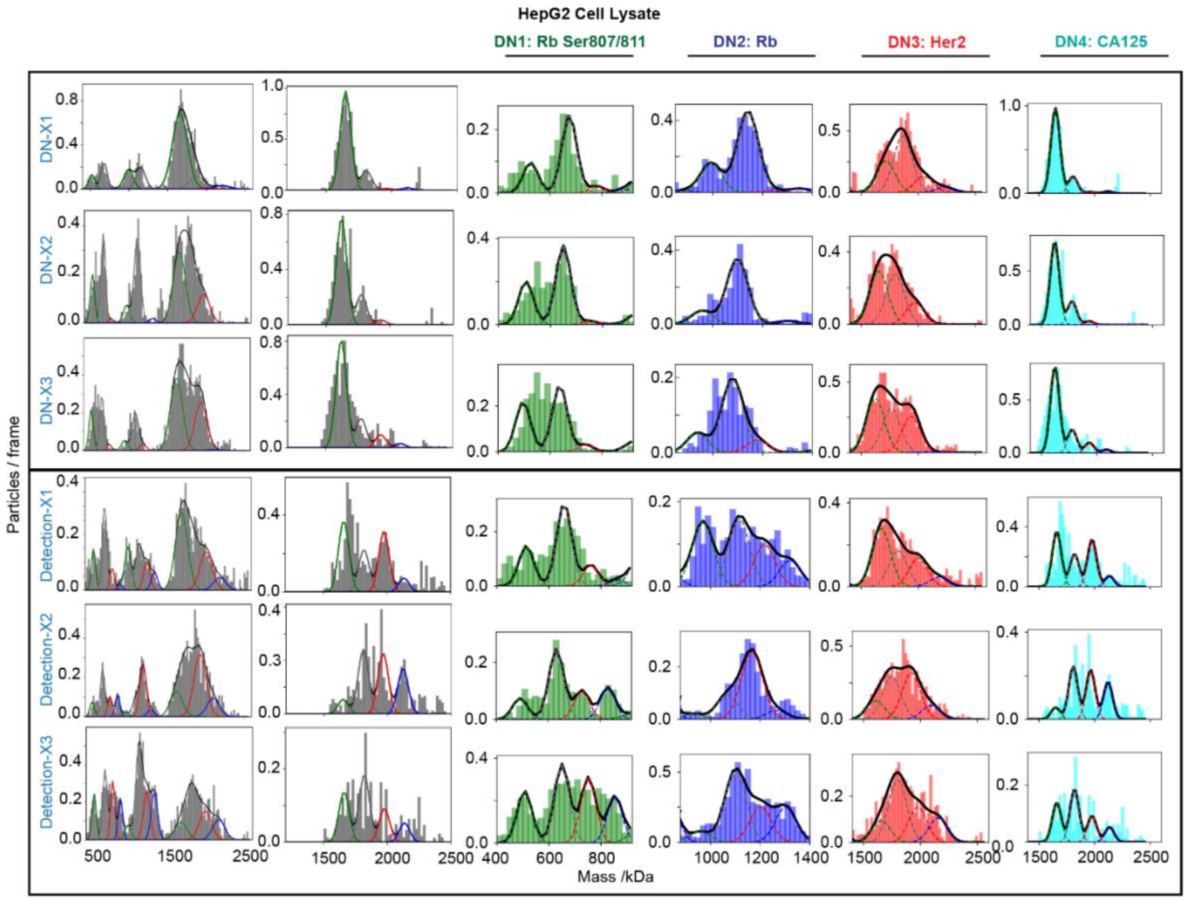

***Fig. S22: Replicate-level DN characterisation and detection for PTM-resolved nanosensors & CA125 in HepG2 lysate (data correspond to Fig. 5b and Fig. S13, S15).***

*Representative mass histograms for all three replicates (DN-X1, DN-X2, DN-X3) for the four mass- and diffusion-encoded nanosensors (DN1: phosphorylated Rb Ser807/811, DN2: total Rb, DN3: Her2, DN4: CA125) measured in HepG2 cell lysate. For each nanosensor, the top three rows show the DN-only measurements and the bottom three rows show the corresponding detection measurements performed on the same DN populations after exposure to lysate.*

*Grey bars show the measured mass distributions of individual replicates. Overlaid curves show the Gaussian components used for ROI-based quantification: unmodified DN (black), antibody-bound DN (grey), DN carrying one target molecule (red), and DN carrying two target molecules (blue). DN3 (Her2) shows a clear and reproducible Her2-specific peak in all three replicates. Importantly, these independent measurements again yield positive Her2 detection with comparable enrichment in HepG2 lysate.*

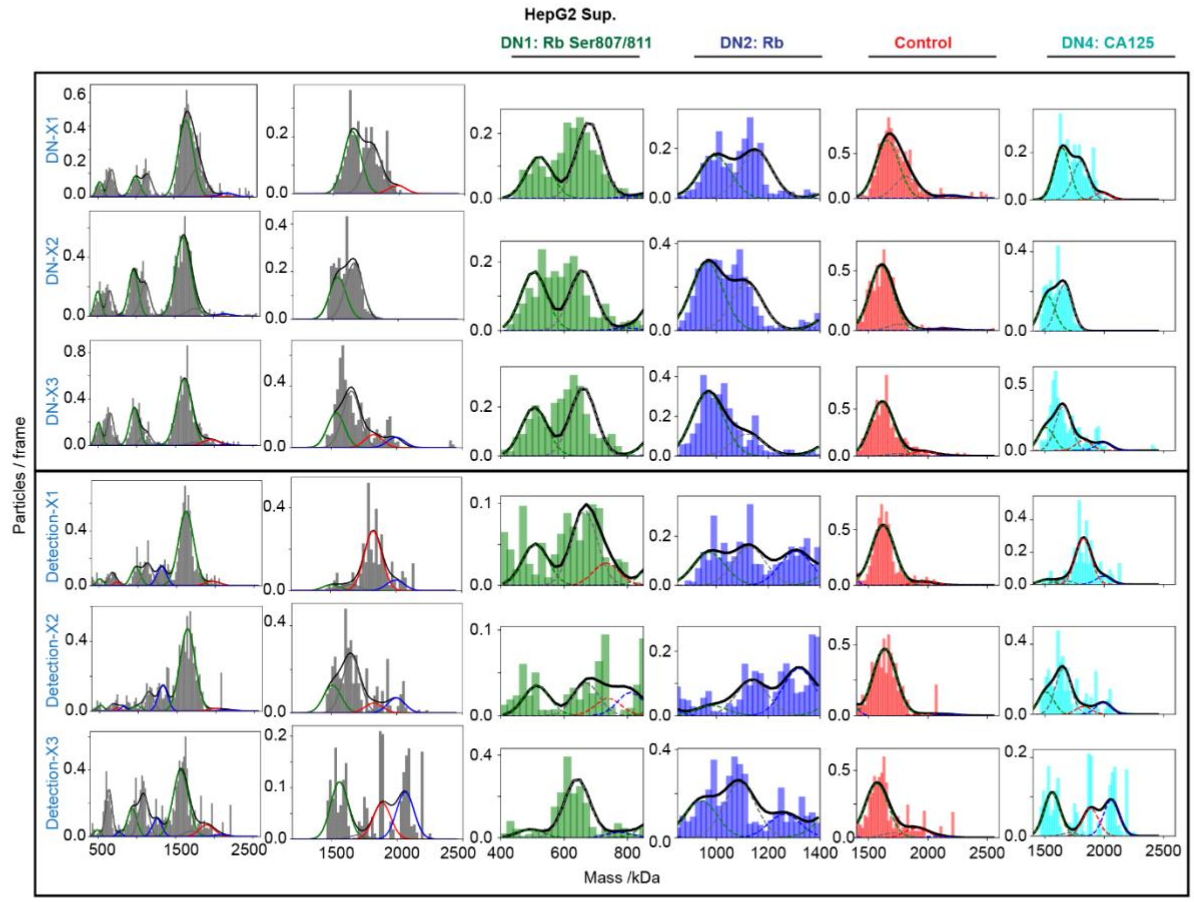

***Fig. S23: Replicate-level DN characterisation and detection for PTM-resolved nanosensors & CA125 in HepG2-SupN lysate (data correspond to Fig. 5b and Fig. S13, S15).***

*Representative mass histograms for all three replicates (DN-X1, DN-X2, DN-X3) for the four mass- and diffusion-encoded nanosensors (DN1: phosphorylated Rb Ser807/811, DN2: total Rb, DN3: control origami without antibody, DN4: CA125) measured in HepG2-SupN lysate. For each nanosensor, the top three rows show the DN-only measurements and the bottom three rows show the corresponding detection measurements performed on the same DN populations after exposure to lysate.*

*Grey bars show the measured mass distributions of individual replicates. Overlaid curves show the Gaussian components used for ROI-based quantification: unmodified DN (black), antibody-bound DN (grey), DN carrying one target molecule (red), and DN carrying two target molecules (blue). DN3 (control) contains no antibody and therefore reports only the intrinsic background of the system; rare high-mass events in this channel reflect background noise. In downstream enrichment analysis, only peaks exceeding this control-defined noise floor are considered target-specific.*

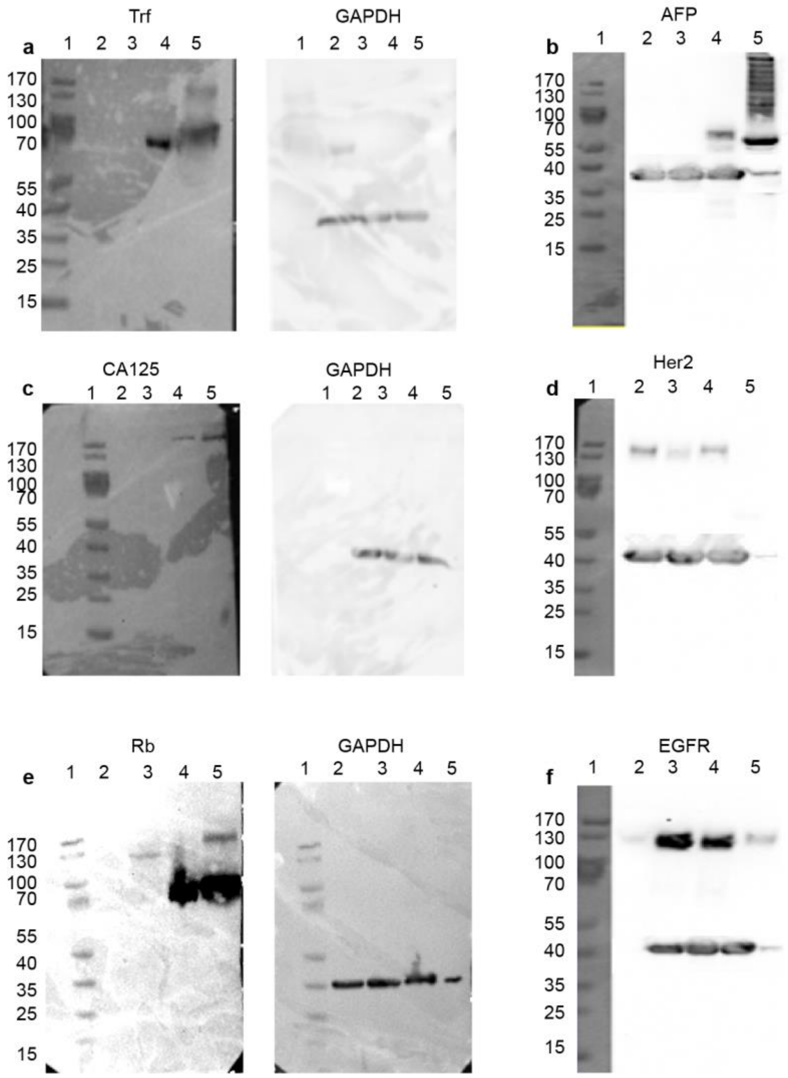

***Fig. S24: Western blot validation of protein expression across the lysates used in this study.*** *Representative western blots for all targets analysed. Lane order in all panels: 1 = molecular-weight ladder, 2 = BxPC-3, 3 = AsPC-1, 4 = HepG2, 5 = HepG2-SupN. For each target, the corresponding GAPDH loading control is shown from the same membrane.* ***a****, Transferrin (Trf) and GAPDH.* ***b****, AFP and GAPDH.* ***c****, CA125 and GAPDH.* ***d****, Her2 and GAPDH.* ***e****, Rb and GAPDH.* ***f****, EGFR and GAPDH. In* ***b****,* ***d****, and* ***f****, the GAPDH is overlaid in the same image. Band patterns across the four lysates are consistent with the nanosensor-derived enrichment profiles.*

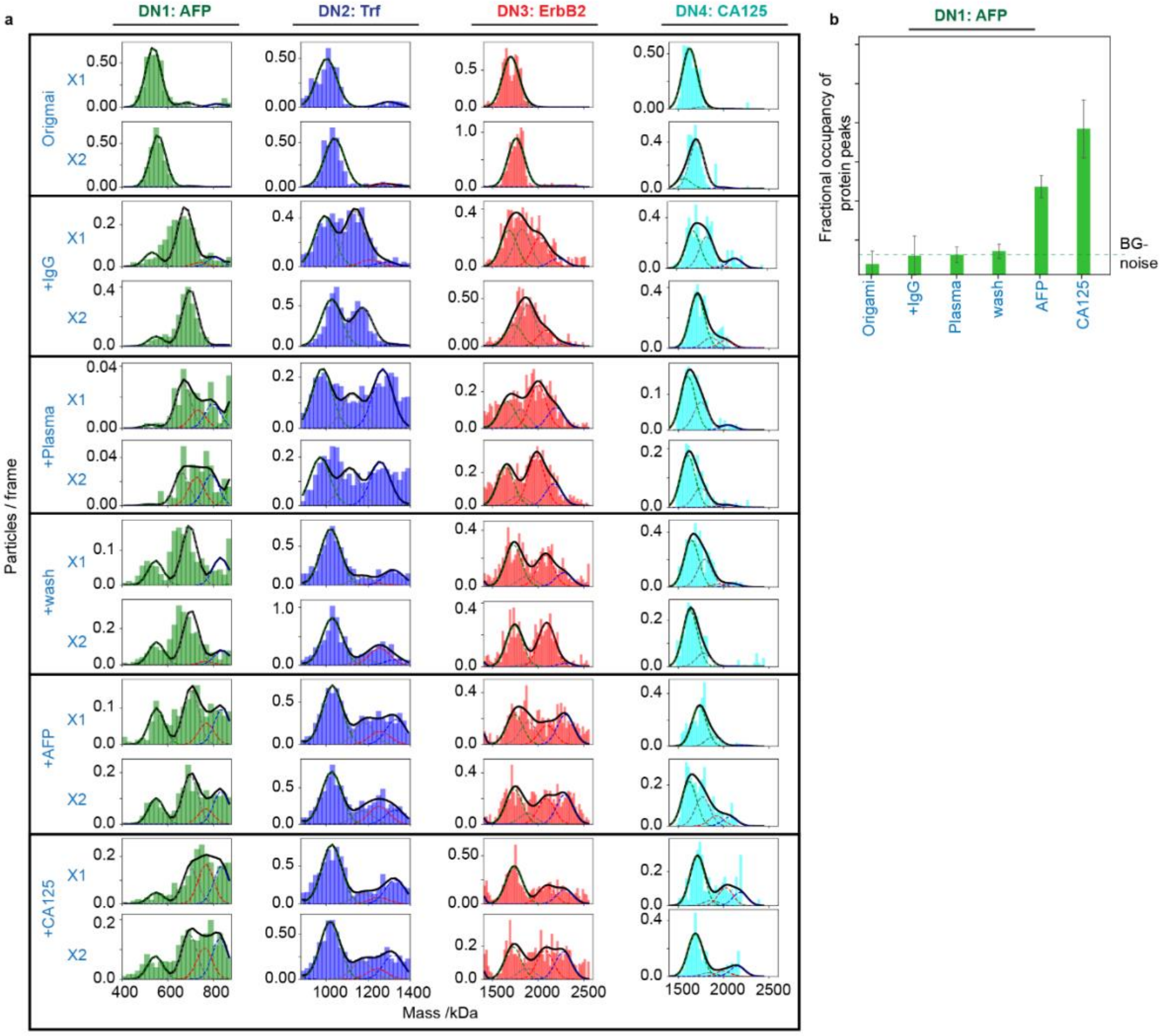

***Fig. S25: Multiplexed protein detection in complex media, plasma (data correspond to Fig. 5c–e).*** *a, ROI-resolved mass histograms for four mass- and diffusion-encoded nanosensors across the sequential plasma assay, across two technical repeats. The four nanosensors (DN1: AFP, DN2: Trf, DN3: ErbB2, DN4: CA125) were first recorded as origami-only baselines and then functionalised with their respective antibodies, yielding four distinct DNs (grey antibody peaks in all channels). Upon exposure to 1% human plasma containing native Trf and spiked ErbB2, only the corresponding nanosensors (DN2 for Trf and DN3 for ErbB2) generated protein-bound peaks (red = one protein, blue = two proteins) within their ROIs, confirming selective detection in a complex background.*

*DN1 (AFP) exhibits a marked loss of contrast in plasma, with broadened distributions arising from its lower mass and higher diffusion, which reduce resolvability against the plasma background. Importantly, following the wash step, DN1 returns to its original baseline and shows no increase above the noise level, confirming the absence of AFP signal in plasma.*

*Sequential protein addition then demonstrates programmable, reconfigurable detection: addition of AFP produces a strong AFP-specific peak in DN1, and a final addition of CA125 yields a clear CA125 peak exclusively in DN4. Because AFP and CA125 additions were performed sequentially without an intermediate wash, the AFP peak increases further due to the prolonged incubation time. Gaussian components denote unmodified DN (black), antibody-bound DN (grey), and target-bound species carrying one (red) or two (blue) protein molecules.*

***b*, *Average fractional occupancy analysis for DN1 (AFP).*** *Quantification of AFP-specific signal across all assay stages. Each bar represents the mean fractional occupancy of the protein-bound species (sum of +1 and +2 protein states) across the two technical repeats, with error bars indicating the inter-repeat deviation. The dashed line denotes the background-noise threshold defined from the origami and DN baselines. AFP addition produces a clear increase above this threshold, confirming specific detection, whereas plasma exposure alone yields no AFP signal after washing, consistent with non-detection in plasma.*

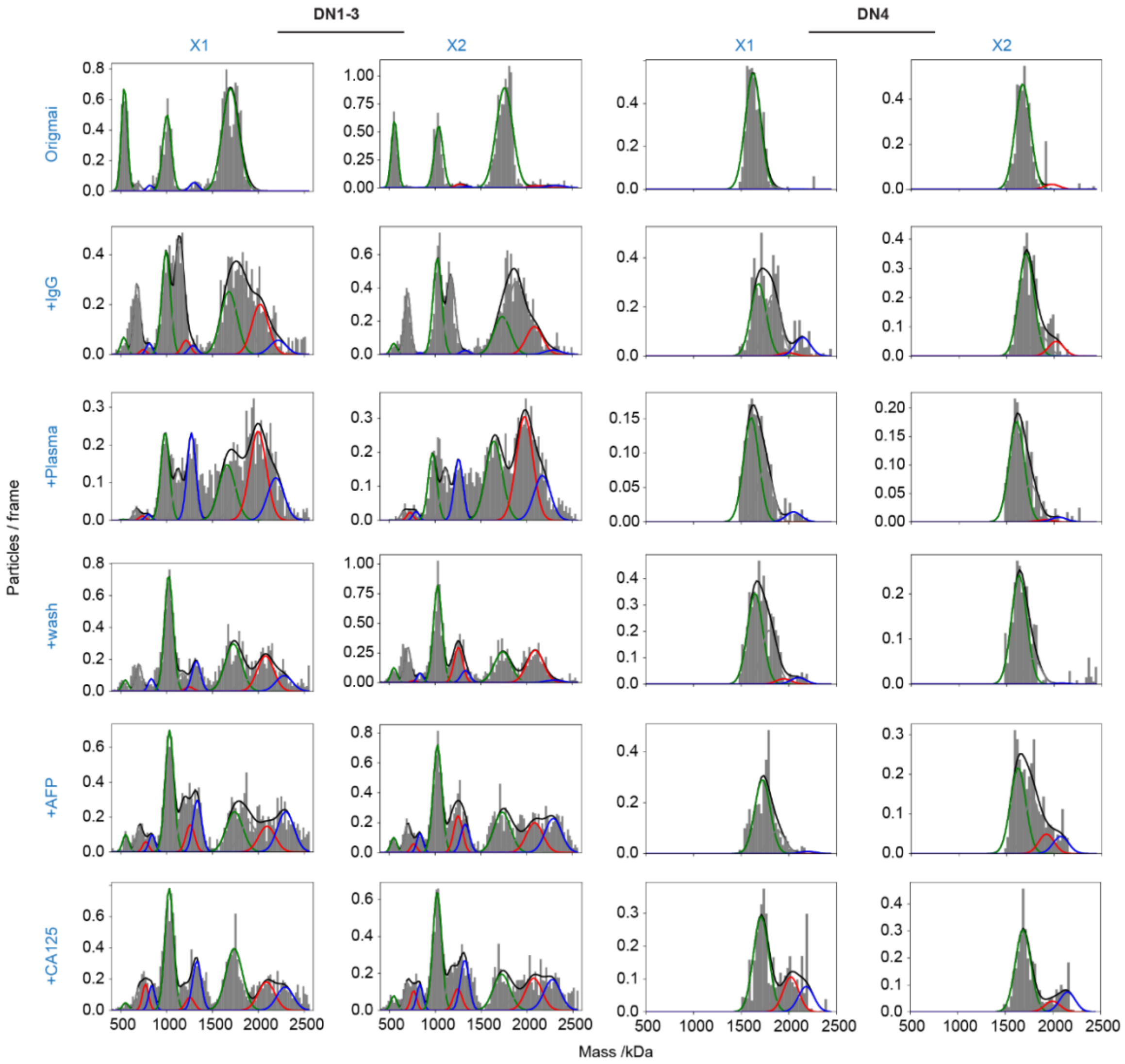

***Fig. S26: Technical-repeat mass histograms for the plasma multiplexing assay (data correspond to Fig. 5c–e and Fig. S25a).*** *Representative mass histograms from the sequential plasma assay for the multiplexed panel. The left block (D1–D3) shows the three nanosensors DN1–DN3 (AFP, Trf, ErbB2), while the right block (D4) shows the CA125 nanosensor (DN4), all measured together on the same SLB in two technical repeats (X1, X2). Rows correspond to the six stages of the assay: origami baseline, antibody functionalisation (+IgG), exposure to 1% human plasma, wash, AFP addition, and CA125 addition. Grey bars show the measured mass distributions; overlaid Gaussian components denote unmodified DN (black), antibody-bound DN (grey), and target-bound species carrying one (red) or two (blue) protein molecules.*

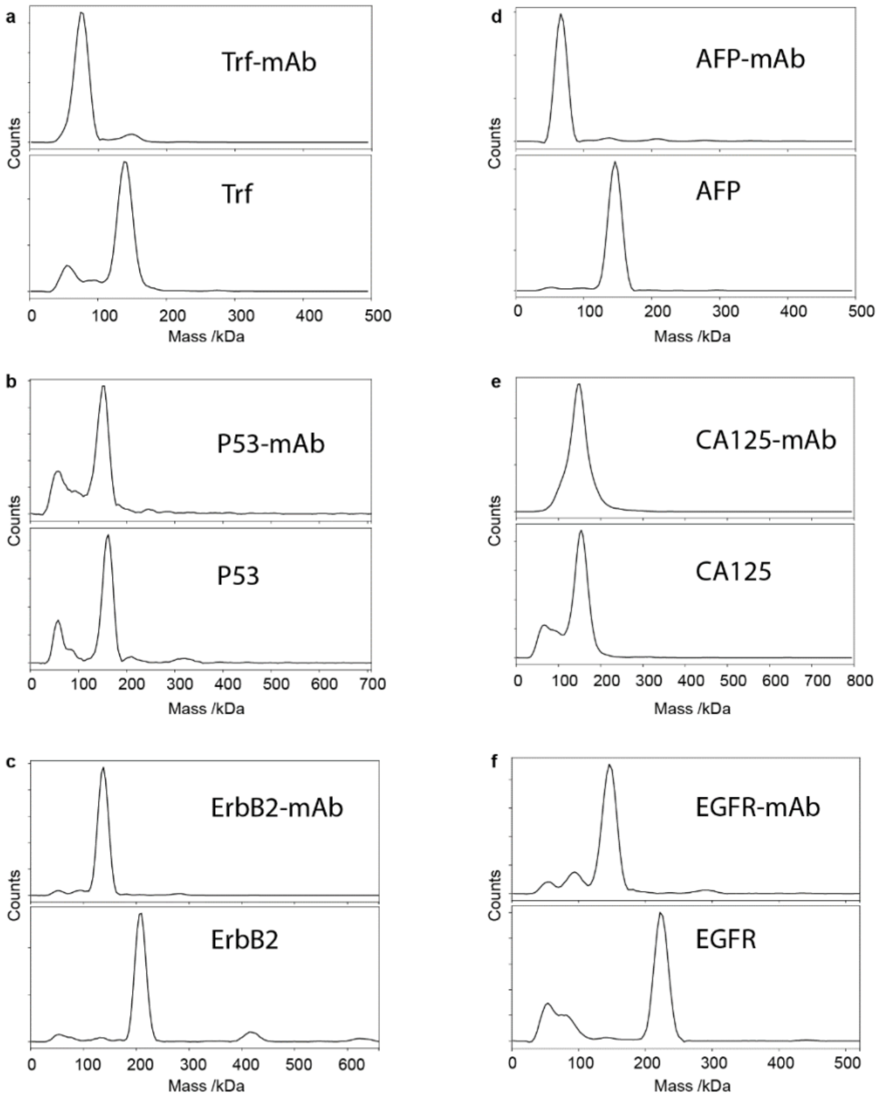

***Fig. S27: Solution-phase mass photometry (landing assay) reference for all protein targets.***
*Representative 10 nM MP landing assays for each monoclonal antibody (top panels) and the corresponding target protein (bottom panels). Measured single-molecule masses were: Trf-mAb (146 kDa) and Trf (76 kDa) (a); p53-mAb (150 kDa) and p53 (82 dimer and 160 tetramer kDa) (b); ErbB2-mAb (146 kDa) and ErbB2 (208 kDa) (c); AFP-mAb (147 kDa) and AFP (67 kDa) (d); CA125-mAb (148 kDa) and CA125 ( 153 kDa) (e); EGFR-mAb (146 kDa) and EGFR (223 kDa) (f).These solution-phase masses were used to initialise and constrain Gaussian components in all SLB-based nanosensor fits.*

| Ligands | Mass |
|---|---|
| **Trf** | 76 kDa |
| **p53** | 160 kDa (tetramer) |
| **EGF** | 6 kDa |
| **AFP** | 67 kDA |
| **ErbB2 (Her2)** | 200 kDa |
| **CA125** | 153 kDa |
| **EGFR** | 223 kDa |
| **Rb** | 100 kDa |
| **mAb** | 146 kDa |

***Table 1.*** *Mass references summary*

| | **BxPC-1** | **AsPC-1** | **HepG2** | **HepG2_Sup** |
|---|---|---|---|---|
| **AFP** | P=0.07 | P=0.13 | 5.13 ± 0.08 | 13.3 ± 0.13 |
| **Trf** | P=0.85 | P=0.17 | 17.36 ± 0.14 | 7.4 ± 0.15 |
| **Her2** | 2 ± 0.08 | 5.6 ± 0.18 | 1.63 ± 0.07 | P=0.22 |
| **EGFR** | 7.1 ± 0.09 | 2.83 ± 0.09 | 11 ± 0.14 | 5.5 ± 0.18 |
| **p807/811 Rb** | P=0.69 | P=0.22 | 4.74 ± 0.14 | P=0.18 |
| **Rb** | P=0.18 | P=0.65 | 6.64 ± 0.13 | 4.2 ± 0.17 |
| **CA125** | P=0.98 | P=0.44 | 2.5 ± 0.14 | 5.44 ± 0.22 |

***Table 2.*** *Protein enrichment values and associated standard errors across the three replicates prior to normalisation for the cell-lysate experiments. The listed entries represent statistically significant enrichments ($p < 0.05$), reflecting differential expression of protein targets across lysates from AsPc-1 and BxPc-3 (pancreatic), HepG2 (liver), and HepG2-SupN (secreted fraction). Entries shown in grey denote cases without statistical significance ($p > 0.05$).*

### III. Origami Designs:

#### a. 1496 Scaffold fragment-based origami

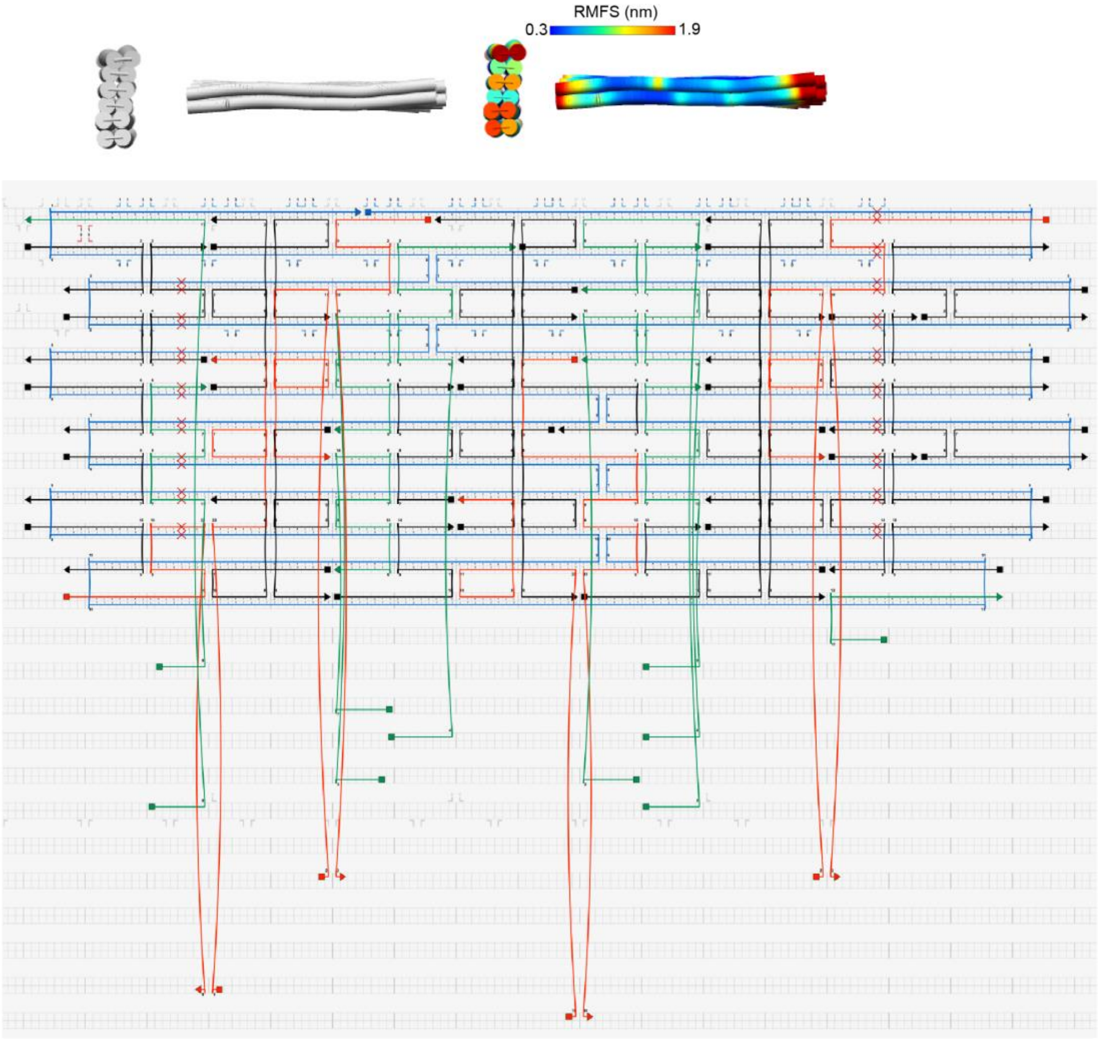

***Fig. S28. 1496-Scaffold based origami design.***
*CanDo6 3D structure and local flexibility predictions are shown as a heat map indicating the local root-mean-square fluctuations (RMSFs) of the origami (top). The corresponding Cadnano strand diagram for the two-layer DNA-origami design, routed on a square lattice, is shown below. The scaffold routing is depicted in dark blue and the staple strands in black. Staples on the bottom layer carry 5' extensions (green; 10 in this design) complementary to a universal 3' cholesterol-modified oligonucleotide used for SLB anchoring. Four positions on the top layer carry capture strands (red; four in this design), implemented as 3' staple extensions complementary to the oligonucleotide conjugated to the antibody of interest, thereby defining a programmable DNA nanosensor whose protein target is specified by the antibody. To improve antibody attachment and elevate the recognition element above the origami surface, a 6-nt spacer was introduced between the staple and the capture sequence. These 6 nt are complementary to a 6-nt extension on the neighbouring 5' oligo at the same break point, providing additional stabilisation through base stacking Scaffold and oligonucleotide sequences are provided in section II Sequences.*

### b. 792 Scaffold fragment-based origami

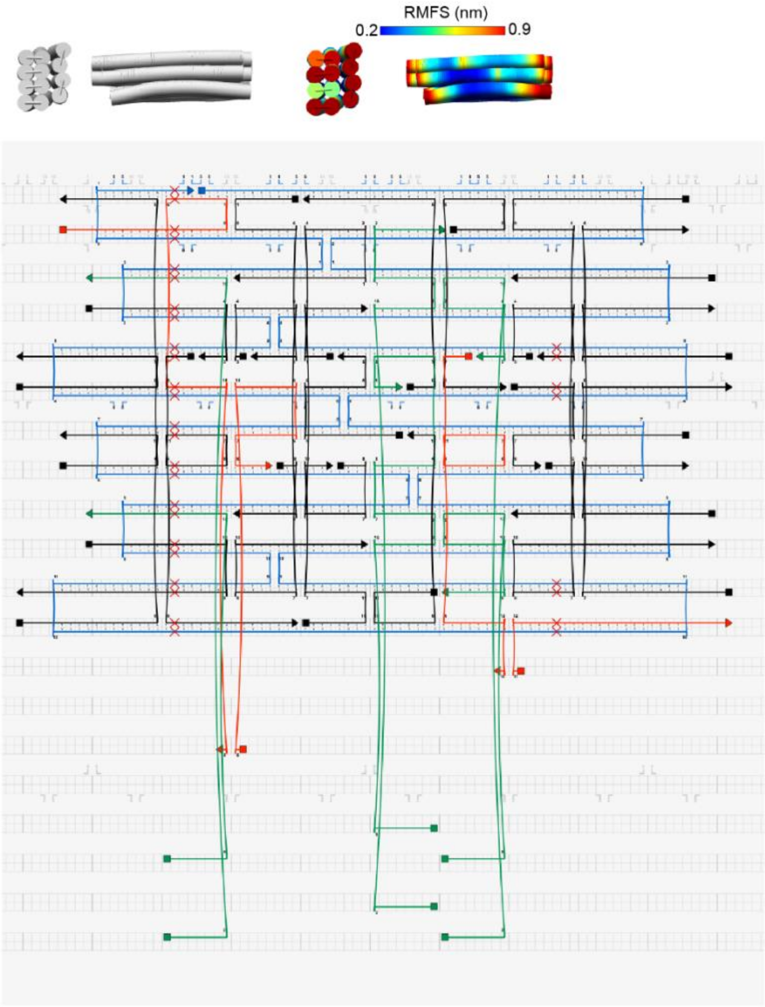

***Fig. S29. 792-Scaffold based origami design.***

*CanDo 3D structure and local flexibility predictions are shown as a heat map indicating the local root-mean-square fluctuations (RMSFs) of the origami (top). The corresponding Cadnano strand diagram for the three-layer DNA-origami design, routed on a square lattice, is shown below. The scaffold routing is depicted in dark blue and the staple strands in black. Staples on the bottom layer carry 5' extensions (green; six in this design) complementary to a universal 3' cholesterol-modified oligonucleotide used for SLB anchoring. Two positions on the top layer carry capture strands (red; two in this design), implemented as 3' staple extensions complementary to the oligonucleotide conjugated to the antibody of interest, thereby defining a programmable DNA nanosensor whose protein target is specified by the antibody. To improve antibody attachment and elevate the recognition element above the origami surface, a 6-nt spacer was introduced between the staple and the capture sequence. These 6 nt are complementary to a 6-nt extension on the neighbouring 5' oligo at the same break point, providing additional stabilisation through base stacking. Scaffold and oligonucleotide sequences are provided in section II Sequences.*

### c. 2288 Scaffold fragment-based origami

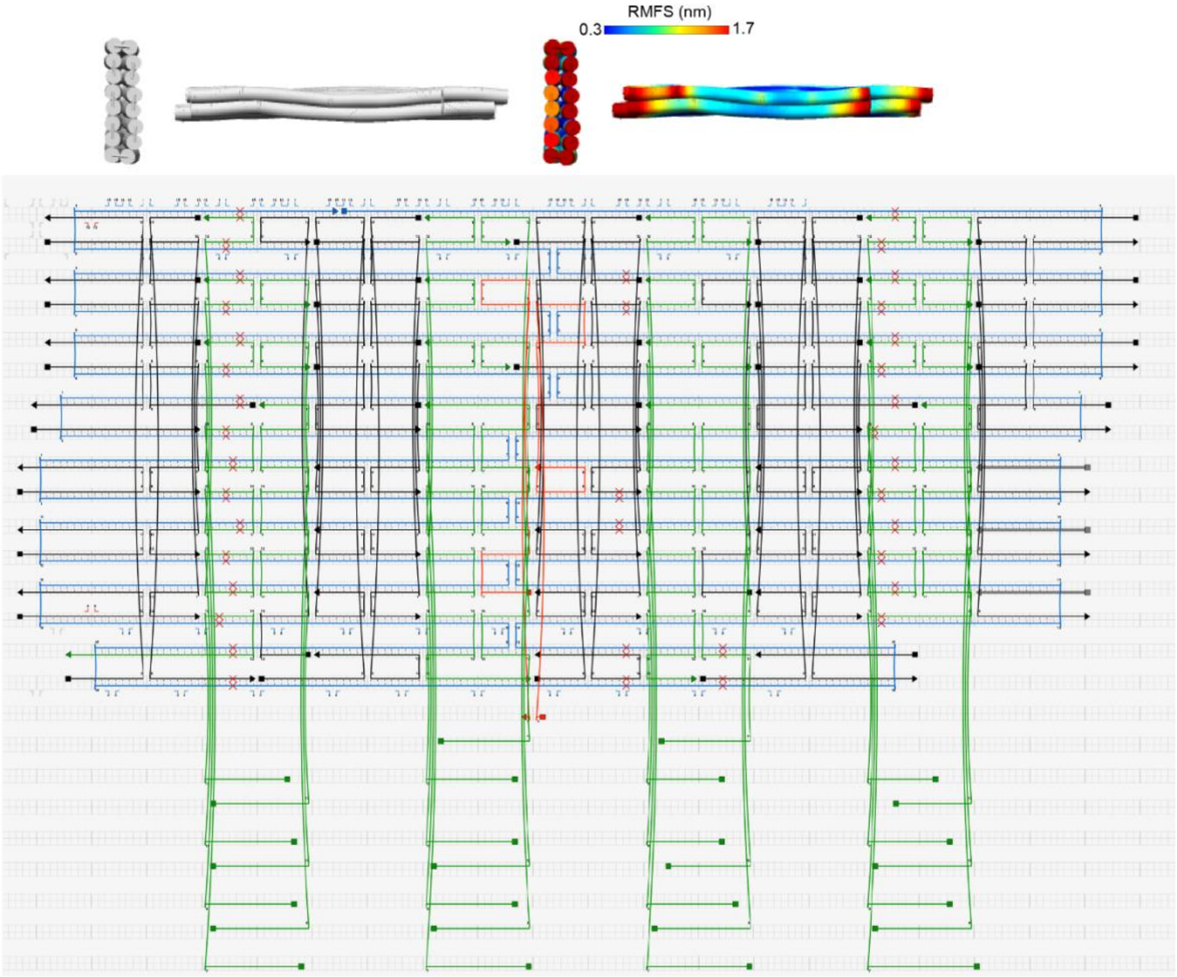

***Fig. S30. 2288-Scaffold based origami design.***

*CanDo 3D structure and local flexibility predictions are shown as a heat map indicating the local root-mean-square fluctuations (RMSFs) of the origami (top). The corresponding Cadnano strand diagram for the two-layer DNA-origami design, routed on a square lattice, is shown below. The scaffold routing is depicted in dark blue and the staple strands in black. Staples on the bottom layer carry 5' extensions (green; twenty-eight in this design) complementary to a universal 3' cholesterol-modified oligonucleotide used for SLB anchoring. Two positions on the top layer carry capture strands (red; one in this design), implemented as 3' staple extensions complementary to the oligonucleotide conjugated to the antibody of interest, thereby defining a programmable DNA nanosensor whose protein target is specified by the antibody. To improve antibody attachment and elevate the recognition element above the origami surface, a 6-nt spacer was introduced between the staple and the capture sequence. These 6 nt are complementary to a 6-nt extension on the neighbouring 5' oligo at the same break point, providing additional stabilisation through base stacking. Scaffold and oligonucleotide sequences are provided in section II Sequences.*

### d. 2288 Scaffold fragment-based origami

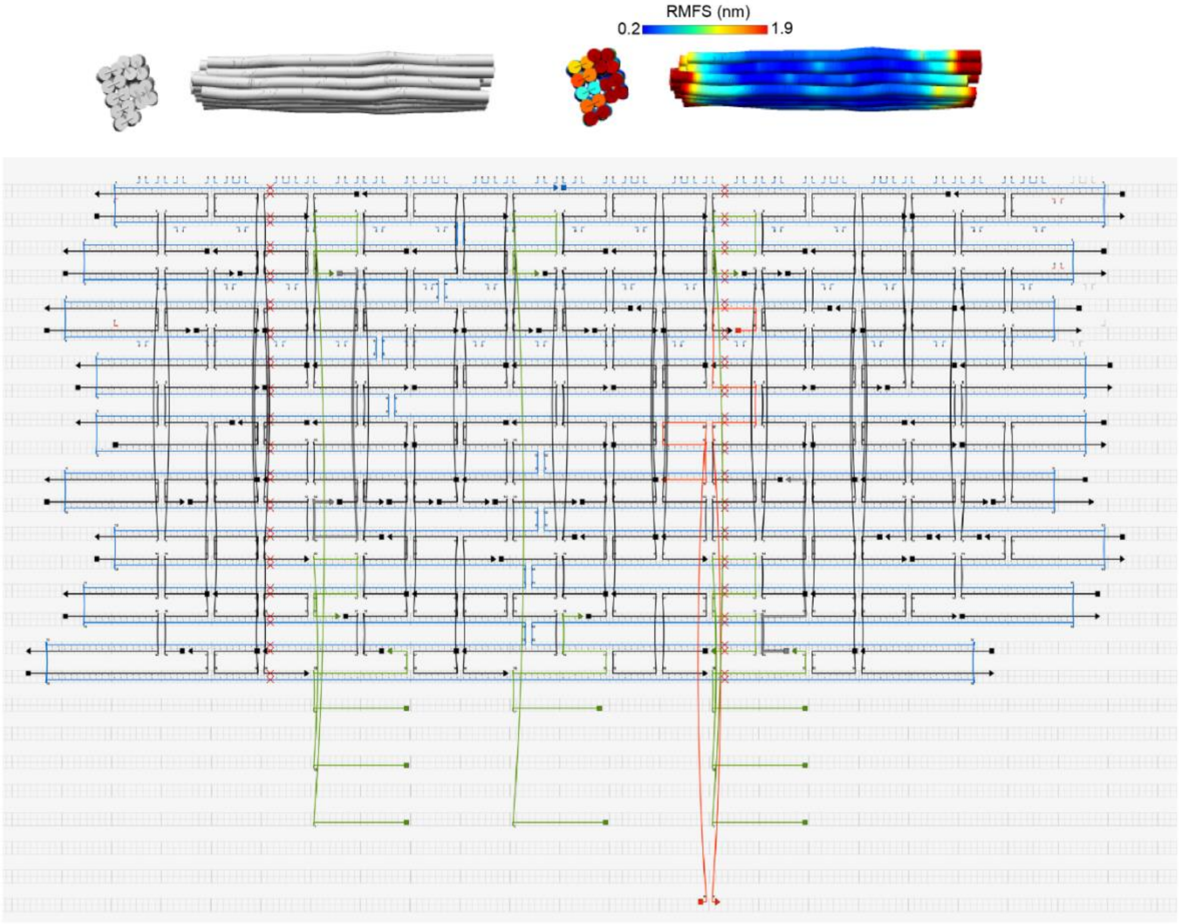

***Fig. S31. 2824-Scaffold based origami design.***

*CanDo 3D structure and local flexibility predictions are shown as a heat map indicating the local root-mean-square fluctuations (RMSFs) of the origami (top). The corresponding Cadnano strand diagram for the four-layer DNA-origami design, routed on a square lattice, is shown below. The scaffold routing is depicted in dark blue and the staple strands in black. Staples on the bottom layer carry 5′ extensions (green;*

*eight in this design) complementary to a universal 3′ cholesterol-modified oligonucleotide used for SLB anchoring. Two positions on the top layer carry capture strands (red; one in this design), implemented as 3′ staple extensions complementary to the oligonucleotide conjugated to the antibody of interest, thereby defining a programmable DNA nanosensor whose protein target is specified by the antibody. To improve antibody attachment and elevate the recognition element above the origami surface, a 6-nt spacer was introduced between the staple and the capture sequence. These 6 nt are complementary to a 6-nt extension on the neighbouring 5′ oligo at the same break point, providing additional stabilisation through base stacking. Scaffold and oligonucleotide sequences are provided in section II Sequences.*

### IV. Sequences:

#### 1. 1496 scaffold fragment:

AAATCTCCGTTGTACTTTGTTTCGCGCTTGGTATAATCGCTGGGGGTCAAAGATGAGTGTTTTAG
TGTATTCTTTTGCCTCTTTCGTTTTAGGTTGGTGCCTTCGTAGTGGCATTACGTATTTTACCCGTTT
AATGGAAACTTCCTCATGAAAAAGTCTTTAGTCCTCAAAGCCTCTGTAGCCGTTGCTACCCTCGTT
CCGATGCTGTCTTTCGCTGCTGAGGGTGACGATCCCGCAAAAGCGGCCTTTAACTCCCTGCAAG
CCTCAGCGACCGAATATATCGGTTATGCGTGGGCGATGGTTGTTGTCATTGTCGGCGCAACTAT
CGGTATCAAGCTGTTTAAGAAATTCACCTCGAAAGCAAGCTGATAAACCGATACAATTAAAGGCT
CCTTTTGGAGCCTTTTTTTTGGAGATTTTCAACGTGAAAAAATTATTATTCGCAATTCCTTTAGTTG
TTCCTTTCTATTCTCACTCCGCTGAAACTGTTGAAAGTTGTTTAGCAAAATCCCATACAGAAAATT
CATTTACTAACGTCTGGAAAGACGACAAAACTTTAGATCGTTACGCTAACTATGAGGGCTGTCTG
TGGAATGCTACAGGCGTTGTAGTTTGTACTGGTGACGAAACTCAGTGTTACGGTACATGGGTTCC
TATTGGGCTTGCTATCCCTGAAAATGAGGGTGGTGGCTCTGAGGGTGGCGGTTCTGAGGGTGG
CGGTTCTGAGGGTGGCGGTACTAAACCTCCTGAGTACGGTGATACACCTATTCCGGGCTATACT
TATATCAACCCTCTCGACGGCACTTATCCGCCTGGTACTGAGCAAAACCCCGCTAATCCTAATCC
TTCTCTTGAGGAGTCTCAGCCTCTTAATACTTTCATGTTTCAGAATAATAGGTTCCGAAATAGGCA
GGGGGCATTAACTGTTTATACGGGCACTGTTACTCAAGGCACTGACCCCGTTAAAACTTATTACC
AGTACACTCCTGTATCATCAAAAGCCATGTATGACGCTTACTGGAACGGTAAATTCAGAGACTGC
GCTTTCCATTCTGGCTTTAATGAGGATTTATTTGTTTGTGAATATCAAGGCCAATCGTCTGACCTG
CCTCAACCTCCTGTCAATGCTGGCGGCGGCTCTGGTGGTGGTTCTGGTGGCGGCTCTGAGGGT
GGTGGCTCTGAGGGTGGCGGTTCTGAGGGTGGCGGCTCTGAGGGAGGCGGTTCCGGTGGTGG
CTCTGGTTCCGGTGATTTTGATTATGAAAAGATGGCAAACGCTAATAAGGGGGCTATGACCGAAA
ATGCCGATGAAAACGCGCTACAGTCTGACGCTAAAGGCAAACTTGATTCTGTCGCTACTGATTAC
GGTGCTGCTATCGATGGTTTCATTGGTGACGTTTCCGGCCTTGCTAATGGTAATGGTGCTACTGG
TGATTTTGCTGGCTCTAATTCCCAAATGGCTCAAGTCGGTGACGGTGATAATTCACCTTTAATGAA
TA

#### 2. 792 scaffold fragment:

CTTTCGGGCTTCCTCTTAATCTTTTTGATGCAATCCGCTTTGCTTCTGACTATAATAGTCAGGGTA
AAGACCTGATTTTTGATTTATGGTCATTCTCGTTTTCTGAACTGTTTAAAGCATTTGAGGGGGATT
CAATGAATATTTATGACGATTCCGCAGTATTGGACGCTATCCAGTCTAAACATTTTACTATTACCC
CCTCTGGCAAAACTTCTTTTGCAAAAGCCTCTCGCTATTTTGGTTTTTATCGTCGTCTGGTAAACG
AGGGTTATGATAGTGTTGCTCTTACTATGCCTCGTAATTCCTTTTGGCGTTATGTATCTGCATTAG
TTGAATGTGGTATTCCTAAATCTCAACTGATGAATCTTTCTACCTGTAATAATGTTGTTCCGTTAGT
TCGTTTTATTAACGTAGATTTTTCTTCCCAACGTCCTGACTGGTATAATGAGCCAGTTCTTAAAATC
GCATAAGGTAATTCACAATGATTAAAGTTGAAATTAAACCATCTCAAGCCCAATTTACTACTCGTT
CTGGTGTTTCTCGTCAGGGCAAGCCTTATTCACTGAATGAGCAGCTTTGTTACGTTGATTTGGGT
AATGAATATCCGGTTCTTGTCAAGATTACTCTTGATGAAGGTCAGCCAGCCTATGCGCCTGGTCT
GTACACCGTTCATCTGTCCTCTTTCAAAGTTGGTCAGTTCGGTTCCCTTATGATTGACCGTCTGC
GCCTCGTTCCGGCTAAGTAACATGGAGCAGGTCGCGGATTTCGACACAATTTATCAGGCGATGA
TAC

#### 3. 2288 scaffold fragment:

CTTTCGGGCTTCCTCTTAATCTTTTTGATGCAATCCGCTTTGCTTCTGACTATAATAGTCAGGGTA
AAGACCTGATTTTTGATTTATGGTCATTCTCGTTTTCTGAACTGTTTAAAGCATTTGAGGGGGATT
CAATGAATATTTATGACGATTCCGCAGTATTGGACGCTATCCAGTCTAAACATTTTACTATTACCC
CCTCTGGCAAAACTTCTTTTGCAAAAGCCTCTCGCTATTTTGGTTTTTATCGTCGTCTGGTAAACG
AGGGTTATGATAGTGTTGCTCTTACTATGCCTCGTAATTCCTTTTGGCGTTATGTATCTGCATTAG
TTGAATGTGGTATTCCTAAATCTCAACTGATGAATCTTTCTACCTGTAATAATGTTGTTCCGTTAGT

TCGTTTTATTAACGTAGATTTTTCTTCCCAACGTCCTGACTGGTATAATGAGCCAGTTCTTAAAATC
GCATAAGGTAATTCACAATGATTAAAGTTGAAATTAAACCATCTCAAGCCCAATTTACTACTCGTT
CTGGTGTTTCTCGTCAGGGCAAGCCTTATTCACTGAATGAGCAGCTTTGTTACGTTGATTTGGGT
AATGAATATCCGGTTCTTGTCAAGATTACTCTTGATGAAGGTCAGCCAGCCTATGCGCCTGGTCT
GTACACCGTTCATCTGTCCTCTTTCAAAGTTGGTCAGTTCGGTTCCCTTATGATTGACCGTCTGC
GCCTCGTTCCGGCTAAGTAACATGGAGCAGGTCGCGGATTTCGACACAATTTATCAGGCGATGA
TACAAATCTCCGTTGTACTTTGTTTCGCGCTTGGTATAATCGCTGGGGGTCAAAGATGAGTGTTTT
AGTGTATTCTTTTGCCTCTTTCGTTTTAGGTTGGTGCCTTCGTAGTGGCATTACGTATTTTACCCG
TTTAATGGAAACTTCCTCATGAAAAAGTCTTTAGTCCTCAAAGCCTCTGTAGCCGTTGCTACCCTC
GTTCCGATGCTGTCTTTCGCTGCTGAGGGTGACGATCCCGCAAAAGCGGCCTTTAACTCCCTGC
AAGCCTCAGCGACCGAATATATCGGTTATGCGTGGGCGATGGTTGTTGTCATTGTCGGCGCAAC
TATCGGTATCAAGCTGTTTAAGAAATTCACCTCGAAAGCAAGCTGATAAACCGATACAATTAAAGG
CTCCTTTTGGAGCCTTTTTTTTGGAGATTTTCAACGTGAAAAAATTATTATTCGCAATTCCTTTAGT
TGTTCCTTTCTATTCTCACTCCGCTGAAACTGTTGAAAGTTGTTTAGCAAAATCCCATACAGAAAA
TTCATTTACTAACGTCTGGAAAGACGACAAAACTTTAGATCGTTACGCTAACTATGAGGGCTGTCT
GTGGAATGCTACAGGCGTTGTAGTTTGTACTGGTGACGAAACTCAGTGTTACGGTACATGGGTTC
CTATTGGGCTTGCTATCCCTGAAAATGAGGGTGGTGGCTCTGAGGGTGGCGGTTCTGAGGGTG
GCGGTTCTGAGGGTGGCGGTACTAAACCTCCTGAGTACGGTGATACACCTATTCCGGGCTATAC
TTATATCAACCCTCTCGACGGCACTTATCCGCCTGGTACTGAGCAAAACCCCGCTAATCCTAATC
CTTCTCTTGAGGAGTCTCAGCCTCTTAATACTTTCATGTTTCAGAATAATAGGTTCCGAAATAGGC
AGGGGGCATTAACTGTTTATACGGGCACTGTTACTCAAGGCACTGACCCCGTTAAAACTTATTAC
CAGTACACTCCTGTATCATCAAAAGCCATGTATGACGCTTACTGGAACGGTAAATTCAGAGACTG
CGCTTTCCATTCTGGCTTTAATGAGGATTTATTTGTTTGTGAATATCAAGGCCAATCGTCTGACCT
GCCTCAACCTCCTGTCAATGCTGGCGGCGGCTCTGGTGGTGGTTCTGGTGGCGGCTCTGAGGG
TGGTGGCTCTGAGGGTGGCGGTTCTGAGGGTGGCGGCTCTGAGGGAGGCGGTTCCGGTGGTG
GCTCTGGTTCCGGTGATTTTGATTATGAAAAGATGGCAAACGCTAATAAGGGGGCTATGACCGAA
AATGCCGATGAAAACGCGCTACAGTCTGACGCTAAAGGCAAACTTGATTCTGTCGCTACTGATTA
CGGTGCTGCTATCGATGGTTTCATTGGTGACGTTTCCGGCCTTGCTAATGGTAATGGTGCTACTG
GTGATTTTGCTGGCTCTAATTCCCAAATGGCTCAAGTCGGTGACGGTGATAATTCACCTTTAATG
AATA

4. **2824 scaffold fragment:**

AAATCTCCGTTGTACTTTGTTTCGCGCTTGGTATAATCGCTGGGGGTCAAAGATGAGTGTTTTAG
TGTATTCTTTTGCCTCTTTCGTTTTAGGTTGGTGCCTTCGTAGTGGCATTACGTATTTTACCCGTTT
AATGGAAACTTCCTCATGAAAAAGTCTTTAGTCCTCAAAGCCTCTGTAGCCGTTGCTACCCTCGTT
CCGATGCTGTCTTTCGCTGCTGAGGGTGACGATCCCGCAAAAGCGGCCTTTAACTCCCTGCAAG
CCTCAGCGACCGAATATATCGGTTATGCGTGGGCGATGGTTGTTGTCATTGTCGGCGCAACTAT
CGGTATCAAGCTGTTTAAGAAATTCACCTCGAAAGCAAGCTGATAAACCGATACAATTAAAGGCT
CCTTTTGGAGCCTTTTTTTTGGAGATTTTCAACGTGAAAAAATTATTATTCGCAATTCCTTTAGTTG
TTCCTTTCTATTCTCACTCCGCTGAAACTGTTGAAAGTTGTTTAGCAAAATCCCATACAGAAAATT
CATTTACTAACGTCTGGAAAGACGACAAAACTTTAGATCGTTACGCTAACTATGAGGGCTGTCTG
TGGAATGCTACAGGCGTTGTAGTTTGTACTGGTGACGAAACTCAGTGTTACGGTACATGGGTTCC
TATTGGGCTTGCTATCCCTGAAAATGAGGGTGGTGGCTCTGAGGGTGGCGGTTCTGAGGGTGG
CGGTTCTGAGGGTGGCGGTACTAAACCTCCTGAGTACGGTGATACACCTATTCCGGGCTATACT
TATATCAACCCTCTCGACGGCACTTATCCGCCTGGTACTGAGCAAAACCCCGCTAATCCTAATCC
TTCTCTTGAGGAGTCTCAGCCTCTTAATACTTTCATGTTTCAGAATAATAGGTTCCGAAATAGGCA
GGGGGCATTAACTGTTTATACGGGCACTGTTACTCAAGGCACTGACCCCGTTAAAACTTATTACC
AGTACACTCCTGTATCATCAAAAGCCATGTATGACGCTTACTGGAACGGTAAATTCAGAGACTGC
GCTTTCCATTCTGGCTTTAATGAGGATTTATTTGTTTGTGAATATCAAGGCCAATCGTCTGACCTG
CCTCAACCTCCTGTCAATGCTGGCGGCGGCTCTGGTGGTGGTTCTGGTGGCGGCTCTGAGGGT
GGTGGCTCTGAGGGTGGCGGTTCTGAGGGTGGCGGCTCTGAGGGAGGCGGTTCCGGTGGTGG
CTCTGGTTCCGGTGATTTTGATTATGAAAAGATGGCAAACGCTAATAAGGGGGCTATGACCGAAA

ATGCCGATGAAAACGCGCTACAGTCTGACGCTAAAGGCAAACTTGATTCTGTCGCTACTGATTAC
GGTGCTGCTATCGATGGTTTCATTGGTGACGTTTCCGGCCTTGCTAATGGTAATGGTGCTACTGG
TGATTTTGCTGGCTCTAATTCCCAAATGGCTCAAGTCGGTGACGGTGATAATTCACCTTTAATGAA
TAATTTCCGTCAATATTTACCTTCCCTCCCTCAATCGGTTGAATGTCGCCCTTTTGTCTTTGGCGC
TGGTAAACCATATGAATTTTCTATTGATTGTGACAAAATAAACTTATTCCGTGGTGTCTTTGCGTTT
CTTTTATATGTTGCCACCTTTATGTATGTATTTTCTACGTTTGCTAACATACTGCGTAATAAGGAGT
CTTAATCATGCCAGTTCTTTTGGGTATTCCGTTATTATTGCGTTTCCTCGGTTTCCTTCTGGTAACT
TTGTTCGGCTATCTGCTTACTTTTCTTAAAAAGGGCTTCGGTAAGATAGCTATTGCTATTTCATTGT
TTCTTGCTCTTATTATTGGGCTTAACTCAATTCTTGTGGGTTATCTCTCTGATATTAGCGCTCAATT
ACCCTCTGACTTTGTTCAGGGTGTTCAGTTAATTCTCCCGTCTAATGCGCTTCCCTGTTTTTATGT
TATTCTCTCTGTAAAGGCTGCTATTTTCATTTTTGACGTTAAACAAAAAATCGTTTCTTATTTGGAT
TGGGATAAATAATATGGCTGTTTATTTTGTAACTGGCAAATTAGGCTCTGGAAAGACGCTCGTTAG
CGTTGGTAAGATTCAGGATAAAATTGTAGCTGGGTGCAAAATAGCAACTAATCTTGATTTAAGGCT
TCAAAACCTCCCGCAAGTCGGGAGGTTCGCTAAAACGCCTCGCGTTCTTAGAATACCGGATAAG
CCTTCTATATCTGATTTGCTTGCTATTGGGCGCGGTAATGATTCCTACGATGAAAATAAAAACGGC
TTGCTTGTTCTCGATGAGTGCGGTACTTGGTTTAATACCCGTTCTTGGAATGATAAGGAAAGACA
GCCGATTATTGATTGGTTTCTACATGCTCGTAAATTAGGATGGGATATTATTTTTCTTGTTCAGGA
CTTATCTATTGTTGATAAACAGGCGCGTTCTGCATTAGCTGAACATGTTGTTTATTGTCGTCGTCT
GGACAGAATTACTTTACCTTTTGTCGGTACTTTATATTCTCTTATTACTGGCTCGAAAATGCCTCT
GCCTAAATTACATGTTGGCGTTGTTAAATATGGCGATTCTCAATTAAGCCCTACTGTTGAGCGTTG
GCTTTATACTGGTAAGAATTTGTATAACGCATATGATACTAAACAGGCTTTTTCTAGTAATTATGAT
TCCGGTGTTTATTCTTATTTAACGCCTTATTTATCACACGGTCGGTATTTCAAACCATTAAATTTAG
GTCAGAAGATGAAATTAACTAAAATATATTTGAAAAAGTTTTCTCGCGTTCTTTGTCTTGCGATTG
GATTT

5. **Digestion Oligos:**

5.1. **<u>For 1496:</u>**

- **1496-Dig-1**: GTACAACGGAGATTTGTATCATCGCCTGAT
- **1496_Dig-2**: TGACGGAAATTATTCATTAA

5.2. **<u>For 2824:</u>**

- **1496-Dig-1:** GTACAACGGAGATTTGTATCATCGCCTGAT
- **2824_Dig-1**: TGT AAA TGC TGA TGC AAA TCC AAT CGC AAG

5.3. **<u>For 792:</u>**

- **1496-Dig-1**: GTACAACGGAGATTTGTATCATCGCCTGAT
- **792-Dig-1**: AGCCCGAAAGACTTCAAATA

5.4. **<u>For 2288:</u>**

- **792-Dig-1:** AGCCCGAAAGACTTCAAATA
- **1496_Dig-2**: TGACGGAAATTATTCATTAA

### 6. Origami staples sequences:

#### - 1496-based origami:

| Name | Sequence |
|---|---|
| Core Oligo1 | TTTTTAATCAGTATTATTAGCGTTTGTTTTT |
| Core Oligo2 | TTTTTAGTACCGCCACCCGAACCCATGTACCGTAACATTTTT |
| Core Oligo3 | TTTTTCTGAGTTTCGTCACCAGTATTCC |
| Core Oligo4 | CCGCCCGTATAAATTAAAGCCCGTCGAGA |
| Core Oligo5 | TTTTTATTTGGGAATTAGAGCACCGTTTTTT |
| Core Oligo6 | TTTTTCCAGCATTACATGGCTTTTGATTTTT |
| Core Oligo7 | GCCGCCACTTAACGGGTTTACCGTGGGGTTTT |
| Core Oligo8 | AAGCGCAGCCTTGATAAGCCACCAGAACCGCC |
| Core Oligo9 | AGGCCGTCACCCCGAAGGCACCAACCTAAAATTTTT |
| Core Oligo10 | AGACCTAAACAATTGAAAATCTCCAAAAAAATTTTT |
| Core Oligo11 | TCCCTCAGGGCTTTGAAGTTGCGCACAAAGTACAA |
| Core Oligo12 | TTTTTGCCGCTTTTGCGGGATTTGCAGGGAGTTAAAGTTTTT |
| Core Oligo13 | AGCCCCCGCGACAGCGATAGCAGCCAGCA |
| Core Oligo14 | GGGTAAAATACGTAATACACTCATCCGATATA |
| Core Oligo15 | GCTACAGAAGCCGCCACCCACAACAAGGATCAGAA |
| Core Oligo16 | TTTTTTCTGTATGGGATTTTGGTTAGTAAATGAATTTTTTTT |
| Core Oligo17 | GGGTTGATATAAGTATCCACCCTCCCCTCATT |
| Core Oligo18 | TTCGGTCTAATTGTTTTCACGCTTTCAA |
| Core Oligo19 | TTTTTAGGCTCCAAAAGGAGCCTTGCTG |
| Core Oligo20 | GTCGTCTCAAACTACCAATAGTCAGAACC |
| Core Oligo21 | TTTTTTGATACAGGAGTGTGAGACTCTTTTT |
| Core Oligo22 | TTTTTCCATCTTTTCATAAGCCGCCGTTTTT |
| Core Oligo23 | GCTCAGTACCAGGCGGTCGGAACCCTTGAGTA |
| Core Oligo24 | CAGTTTCATAGCAAGCCAACGCCTATAGGTGT |
| Core Oligo25 | TGAAACATGAAAGTATGGATTAGCTCCAGTAA |
| Core Oligo26 | TTCAGGGAGCGGAGTGAACGATCTATCAGCTT |
| Core Oligo27 | TCACCGGAGAAACCATAATCAAGTGTGAATTA |
| Core Oligo28 | ACAGTGCCACCCTCAGTTCACAAAAGCTTGAT |
| Core Oligo29 | GCCACCCTCAGAACCGAGCCCGGAGTAGCATT |
| Core Oligo30 | TTCATGAGAGCGCGAACGACAATG |
| Core Oligo31 | AAATCACCAGTAGCACCATTAAAGTTGCCTTT |
| Core Oligo32 | TTGCGAATAGCATCGGACGCATAACTTTGACC |
| SLB Oligo1 | AAAGAAAAAGGGGAAAAGCGTCAGGTTTTCATCACCACCGCCCTCAGA |
| SLB Oligo2 | AAAGAAAAAGGGGAAAGCGTCATGACAGGACCACCAGATCAAAA |
| SLB Oligo3 | AAAGAAAAAGGGGAAAACGATTGGTCTCTGAAGTCAGTGCTATTATTC |
| SLB Oligo4 | AAAGAAAAAGGGGAAAACAACAACGGTGAATT |
| SLB Oligo5 | AAAGAAAAAGGGGAAAACCAGCGCACTGTGATGGAAAGGCTTAGCCTAAAGACTTT |
| SLB Oligo6 | AAAGAAAAAGGGGAAATCACCGTCACCGACTTGAGCCTTTTT |
| SLB Oligo7 | AAAGAAAAAGGGGAAACCACAGACGTTAGCGTAGAATAGATAAAGGAA |
| SLB Oligo8 | AAAGAAAAAGGGGAAAGCTTTCGACATCGCCCAACGAGGGCATTAAAC |
| SLB Oligo9 | AAAGAAAAAGGGGAAAATCACCGTACTCAGGAGGTTTTTTTT |
| SLB Oligo10 | AAAGAAAAAGGGGAAACCCAGCGATTATACCAGAAGTTTCTAGCAACG |

| | |
|---|---|
| Capture 1_5' | ACACATAATAAGTTCAGAACCAGGTTGAGGTCGGTCAT |
| Capture 1_3' | TTTTTCTCAAGAGAAGGATTATAAGAGGCTACTGGTATGTGT GTGGAGTAGTGTCAT |
| Capture 2_3' | TTTTTCGAAAGAGGCAAAAGAATACACTAAAGCCACTATCAGCAGATGTGT |
| Capture 2_5' | ACACATCGAAAGACAATAATTTATCGGTTTAAAGTTTT |
| Capture 3_5' | ACACATTCACCAATACCAGAGCCGGCATTTCAGGTCAG |
| Capture 3_3' | CGGAGATTTTATTCATTACCACGGAAACGATGTGT |
| Capture 4_5' | ACACATATGCCCCCTGCCTATTATAAGTGCAGAATGGA |
| Capture 4_3' | TCTTAAACCAAATAAATCCTCATAAGCCCTCACAGCACCAAGAGCTTAATGTGT |

- **792-based origami:**

| **Name** | **Sequence** |
|---|---|
| Core Oligo1 | TTTTTACTTTGAAAGAGGTCATCGCCTTTTT |
| Core Oligo2 | TTTTTAGATTTAGGAAATTCATTGAATTTTTT |
| Core Oligo3 | CCCTCGTCCAGGACAGATTAAATG |
| Core Oligo4 | TTTTTAGGCATAGTAGCGAGAGGCTTTTTTT |
| Core Oligo5 | TTTTTACGAGGCGCAGATAAG |
| Core Oligo6 | AAATCTACTTGTGAATATAA |
| Core Oligo7 | ACGATCAAAAAGATTAAGAG |
| Core Oligo8 | TTTTTCAAGAGTAACCGAACTGACCATTTTT |
| Core Oligo9 | GAAGCCCGAAATCCGCGACAAT |
| Core Oligo10 | TTTTTATTGGGCTTGAGAACCTTCATTTTTT |
| Core Oligo11 | TTTTTTAGTCAGAAGCAAAGCTTTACCCTTTTG |
| Core Oligo12 | TCAGGTCGGATTGCACGATAAAA |
| Core Oligo13 | AAAGGGACGTTTAGAAAGATTCATCAGTTGTTTTT |
| Core Oligo14 | TTTAATCAGTTAATAAGGAT |
| Core Oligo15 | TTTTTTTGCAAAAGAAGTGACTATTATTTTT |
| Core Oligo16 | GGAATCTTGACTGGCTGTGGTTTAATTTCAAC |
| Core Oligo17 | TTTTTAATAAGGCTTGCAACACCAGAACGAGTAGTAATTTTT |
| Core Oligo18 | ACCAAAATAAGAGCAGCCA |
| Core Oligo19 | TTTTTATTATACCAGTCAGAATTACGTTTTT |
| Core Oligo20 | CACGGTCTGGACAAAGCT |
| Core Oligo21 | ATTCTCAACGTAATAGCGTC |
| Core Oligo22 | TTTTTCCCCCTCAAATGCTTTAAA |
| Core Oligo23 | TTTAGACCCATAAATGACTGCTCCATGT |
| Core Oligo24 | ACCAACCGAGACCTGAGAACTAACGGA |
| Core Oligo25 | CCATCATAAATTACCACAT |
| SLB Oligo1 | AAAGAAAAAGGGGAAACAGTTCAGCGGAATCGGAGG |
| SLB Oligo2 | AAAGAAAAAGGGGAAACAATACTGAAAACGAGAATCAAAAA |
| SLB Oligo3 | AAAGAAAAAGGGGAAATCAACTAAATTACAGGGGGAAGAA |
| SLB Oligo4 | AAAGAAAAAGGGGAAATACTTAGCCGGATTTTT |
| SLB Oligo5 | AAAGAAAAAGGGGAAAACAACATTTGCCCAAAATTACAGAAATAGTAACCAG |
| SLB Oligo6 | AAAGAAAAAGGGGAAAGCTCATTCAGTGTTTTT |
| Capture 1_5' | ACACATAACGGTGTCGCATAGGCAAGA |
| Capture 1_3' | TTTTTTGATAAATTGTGTCGAAAGGTAACAGATGATGTGT GTGGAGTAGTGTCAT |

| | |
|---|---|
| Capture 2_5' | GGGTTACATAACACACTATCTACCTTATATGTGT |
| Capture 2_3' | ACACATGCGATTTTAAGAACTGGCTCTTTTT |

- **2288-based origami:**

| **Name** | **Sequence** |
|---|---|
| Core Oligo1 | ATCACCGTAAACAACTTTCAACA |
| Core Oligo2 | AGCAAAGCAATAATTTTCGAGGTGAATTTC |
| Core Oligo3 | GAAGCCCGAATAGAAAAGTTTTGTCGTCTTTC |
| Core Oligo4 | ATCAGGTCAAAAAGGCTCCAATTTTT |
| Core Oligo5 | TTTTTCAGTTCAGAAAACGAGAATTCGGTCGCTGAGGTTTTT |
| Core Oligo6 | ACGTCATAAAAAATAGCCATACAGAGGATTATACC |
| Core Oligo7 | TTTTTGCCATTTGGGATAGCGTCAGACTTTTT |
| Core Oligo8 | TAGCACCATAATCAGTCGCCACCCAGTACCAG |
| Core Oligo9 | ACTGCGGACTTTTGCATGAGGAAGACACTAAA |
| Core Oligo10 | CCCCCTCAAGACGACGGTAATGCCACTACTTTTT |
| Core Oligo11 | AGGGGGTTGTTTAGATCGGAACGACCGATAG |
| Core Oligo12 | TTTGCCTTATTAGAGCCCCAATAGCCAGTACA |
| Core Oligo13 | CAGCACCGTTACCATTAGCCACCAACAGACAG |
| Core Oligo14 | GCGAGAGGATCGTCATGGGATCGTTCGCCCAC |
| Core Oligo15 | TTTTTCCTCGTTTACCAATGCTTTAAATTTTT |
| Core Oligo16 | ATAACGCCTTATTACAGCCTGATAAACCGAAC |
| Core Oligo17 | AGAGCAACTACGTTAAGCTCCATGTTACTTTTTT |
| Core Oligo18 | TTTTTTGTAGCGCGTTAACCGCCACCCTTTTT |
| Core Oligo19 | CCCCCTTAGAACCGCCAGAGGCTGCAGTGCCT |
| Core Oligo20 | AGTTGAGACCACATTCCCAGCGCTTTGAGG |
| Core Oligo21 | ACCCTCAGTTCATCGGCGAGAGGGACCGTACT |
| Core Oligo22 | TTTTTGAAGAAAAAATCACTATCATAACTTTTT |
| Core Oligo23 | GAACAACAAAAAGGAAAAAAGAATTTTCCATT |
| Core Oligo24 | CACCACCGTTAGCGTTTTTTGCTCTCAGAACC |
| Core Oligo25 | TTTTTTCAGAGCCACCCAGAATGGAAAGCTTTTT |
| Core Oligo26 | GGAGGTGTTTAAGATGTGAACCAGGCGCATAGGCT |
| Core Oligo27 | CACCACCATATTCACACATGGCTTTTGATGAT |
| Core Oligo28 | ATACCAGTCTCATTCAACCCAAATCAACGTAATTTTT |
| Core Oligo29 | TACCTTATGAAACACCGAGTAATCTTGACAAG |
| Core Oligo30 | TTTTTCAAAGCTGCAGGACGTTGGTTTTT |
| Core Oligo31 | GGCTTGCCCTGACGAGCGATTTTCATAAGGGAATTGTGT |
| Core Oligo32 | GGCCTTGAGAGCCGCCAACGGGGTAGACTCCT |
| Core Oligo33 | TGGGCTTGATTTCAACAGGACAGAGTACAAC |
| Core Oligo34 | AATCCTCATTAAAGCACCCTCAGGTATAAACTATTCTGA |
| Core Oligo35 | TTTTTGCAGTCTCTGAATTTACCGTT |
| Core Oligo36 | TTTTTTAGCCGGAACGAG |
| Core Oligo37 | TTTTTCTATTTCGGAACCTATAGTTAATGCCCCCTGCTTTTT |
| Core Oligo38 | TTTTTGAAGGCACCAACC |
| Core Oligo39 | TTTTTCCGGAATAGGTGTATCTTGATATAAGTATAGCTTTTT |
| Core Oligo40 | TTTTTCTTGCAGGGAGTT |
| Core Oligo41 | GCAGCGAACTTTTTCAAAAGAAGTGCAGATAC |
| Core Oligo42 | TTTTTACACTGAGTTTCGTCAGAACCCATGTACCGTATTTTT |
| Core Oligo43 | TTTTTAAGGAGCCTTTAATTGTATCG |

| | |
|---|---|
| Core Oligo44 | CAGACGTTACGCCTGTCATTAAAGATCACCAG |
| Core Oligo45 | TTTTTATTTTGCTCACCGACTTGATTTTT |
| Core Oligo46 | TTGCGAATGGATTGCAAGCTTGATAGGGTAGC |
| Core Oligo47 | TTGAAAATCTCCAAATTTACCCTAACAACCACACCCTCA |
| Core Oligo48 | GTTTCAGCGGAGTGAGAAAGTATTAGCATTCCCCCTCATT |

| | |
|---|---|
| SLB Oligo1 | AAAGAAAAAGGGGAAAGTTTATCACAATGACGACTATTACGTCCAAT |
| SLB Oligo2 | AAAGAAAAAGGGGAAAGCATAACCGATATATGACCATAACATTGAAT |
| SLB Oligo3 | AAAGAAAAAGGGGAAACCCTCATACGATCTAAGGAACAACTAAAGGAA |
| SLB Oligo4 | AAAGAAAAAGGGGAAATTCAGGGATTTAGTACAGCGACAGGGTCATAG |
| SLB Oligo5 | AAAGAAAAAGGGGAAACAGGAGGTAGCAAGCAGCAAAGTGAATT |
| SLB Oligo6 | AAAGAAAAAGGGGAAAACTAAAGAAGACAGCACTGGATAGTAGTCAGA |
| SLB Oligo7 | AAAGAAAAAGGGGAAAAAGAGGATGGATTCGCTCACCGGATGACA |
| SLB Oligo8 | AAAGAAAAAGGGGAAATAAAACGACGCGACCTTAAAACGAGGCTCATT |
| SLB Oligo9 | AAAGAAAAAGGGGAAAAACATGAGTGCCGTCATTTTCAATCAAG |
| SLB Oligo10 | AAAGAAAAAGGGGAAAGGAGATTTCTTTGACCCAACTAATTTTGCCAG |
| SLB Oligo11 | AAAGAAAAAGGGGAAAGTAGGTAAGTACTCAGGGCAGGTCAGACGATT |
| SLB Oligo12 | AAAGAAAAAGGGGAAAGCGCAGACTATTCATTGTGAATAA |
| SLB Oligo13 | AAAGAAAAAGGGGAAAGGCTGACCTTTGAAAGTTTAATCAGATTCATC |
| SLB Oligo14 | AAAGAAAAAGGGGAAACCAGTAAAGTGCCCAGCCGCCAGCCGCC |
| SLB Oligo15 | AAAGAAAAAGGGGAAATTACGTAAGTTAGAACTCAAAAAGCGGAA |
| SLB Oligo16 | AAAGAAAAAGGGGAAAAACTACAAGTAAATGAATTTTCTGTATGGGTTTTT |
| SLB Oligo17 | AAAGAAAAAGGGGAAATTGCGCCGAGCTTGCTTTTTCACG |
| SLB Oligo18 | AAAGAAAAAGGGGAAAAAAGGCCGTAAAATACATAAAAACGCATAGTA |
| SLB Oligo19 | AAAGAAAAAGGGGAAAAAACGGGCTTTTGCAAATATTATCAAAA |
| SLB Oligo20 | AAAGAAAAAGGGGAAAGCCACCCTACCCTCAGAGCAAGGCATTAAGAG |
| SLB Oligo21 | AAAGAAAAAGGGGAAAGCGGATAAAAGTATTATCCCTCAGACCAGAAC |
| SLB Oligo22 | AAAGAAAAAGGGGAAAACACTCATGTATCATCGGTAGAAATTGTGAAT |
| SLB Oligo23 | AAAGAAAAAGGGGAAACAAGAGAATAGCGGGGTGCCATCTATCGATAG |
| SLB Oligo24 | AAAGAAAAAGGGGAAACGAAATCAAGAGGCTTACGAGCAAAATA |
| SLB Oligo25 | AAAGAAAAAGGGGAAATGAGTAACGCGTCATAAACAAATA |
| SLB Oligo26 | AAAGAAAAAGGGGAAATGACCAACTTCATCAAAGAACGAGTAGTAAAT |
| SLB Oligo27 | AAAGAAAAAGGGGAAAACAGGAGTTAAGTTTTGCCAGCATACCAGAGC |
| SLB Oligo28 | AAAGAAAAAGGGGAAAAACCGGAGGTCAATAAGAACTACTAACG |

| | |
|---|---|
| Capture 1_3' | AACCCGCCCAGAAGGCATGAAACCTTTCATAA ATGTGT GTGGAGTAGTGTCAT |
| Capture 1_5' | ACACAT TCAGAATATTTAGAAAGAAACAAATGAACGGT |

- **2824-based origami:**

| Name | Sequence |
|---|---|
| Core Oligo1 | TTAGAGGAACAACCAGCATTCCAG |
| Core Oligo2 | AATTCGAAATTATAAGC |
| Core Oligo3 | GAATAGAATACCGCCACTATTTCG |
| Core Oligo4 | CCCATCCTTTAGGCAATCAAGCTAATTTGAGATAATTAGACGATACCCA |
| Core Oligo5 | CAGCGAGTTTAACCATACCTCAAAGAACAAATAAGATAGCAACG |
| Core Oligo6 | TTACCAGCGCCAAAGACGTAGAAAGAAACCAT |

| | |
|---|---|
| Core Oligo7 | AAAAGCTTTCGAAGTACAACGGACTAAAGACTTTTT |
| Core Oligo8 | TCAATTTTAGTTGAAATACCGACCGTG |
| Core Oligo9 | TTTCTTACCAAATAATAAGCAGAGAGACTTATTAC |
| Core Oligo10 | TTTAAGAAATAACGGAGGAGAATT |
| Core Oligo11 | CGGAATCAAAGGCGTTGCGAGAAA |
| Core Oligo12 | CACATCAGCTTTCTCCAAAAACGA |
| Core Oligo13 | TTTCCTTGAATTATCCCAGAGC |
| Core Oligo14 | AAGCTTACCGAAGCCCTT |
| Core Oligo15 | CGAAAGACGGCTTTGAGGAGATTTAAATCCAAGTTA |
| Core Oligo16 | AATCAATATATAAAAGTACCATTA |
| Core Oligo17 | TCTGTATGCATCTTTGGTTGCGCCTTGCAGGGTTTTCATC |
| Core Oligo18 | CCAGACGAAATATACAGTAGGAGAAAAAAAACGAT |
| Core Oligo19 | AGGATACCGATAACCCCCAGATTAAACGGGTAAAAT |
| Core Oligo20 | AAATAGCATAAGACTCATAACATA |
| Core Oligo21 | CCAGGCGGCAACGCCTACAGACAG |
| Core Oligo22 | CTATAATTGAGTAATCAGATCTAT |
| Core Oligo23 | CCCTCATATCGCAAGATTAATTGTAACGAGGGATAAACAC |
| Core Oligo24 | GTCGTCTTCGAAACAAGGTGAATTTCGTCACCAAGTTTGC |
| Core Oligo25 | ACGTAATGGCTGAGGCGACAATGACACC |
| Core Oligo26 | CCCTCAGAACAGAATCCTCAGCAG |
| Core Oligo27 | AAACACTGGATTTTGCTAAACAACAGAGCCA |
| Core Oligo28 | ATTAGCGGTCGTAGGACGCCCAAT |
| Core Oligo29 | TATTTTCAGGTTTTGCCACC |
| Core Oligo30 | AAGCGAGCGCTAAAGAACGCAGGT |
| Core Oligo31 | CATGTACCAGCCCGGATTTTTCACGCCGCCACAGCCGCCA |
| Core Oligo32 | GGTATTCTATATCAGAGCCAGTTACAAATAAGGCCTGTTT |
| Core Oligo33 | TGTCAAGCCTTAAGAGGCATACGC |
| Core Oligo34 | CGATAGCACAGTAGCGACCGC |
| Core Oligo36 | GCTAATGCAGAACGCAATTCTGT |
| Core Oligo37 | CCACCCTCATTTTCAGAGGT |
| Core Oligo38 | GCCGCTTTAAGTTTCCCGATTATACCAAGCGTCC |
| Core Oligo39 | ATAAGTATGTAACACTAAAGTTTT |
| Core Oligo40 | AACTGAACTCCCAATCCAAAATAATAAAGCCATTTCGAGCGGTAA |
| Core Oligo41 | TTTGTTTCCTTAACAAGAAAAATAATAT |
| Core Oligo42 | GGCATTTTTCACCAGTATCACCGT |
| Core Oligo43 | AGCAAGCATAAGCCCACGCTAACGAAAATGAAAATAGC |
| Core Oligo44 | CATGGCTTAGACTCCTCCCTCAGAGTTG |
| Core Oligo45 | CAATAATATCGAGACTTATCC |
| Core Oligo46 | CCTCATTATCATTAAAACCACCGGGTAGCGCGAGTTAAAG |
| Core Oligo47 | AGCCTGTAATGCACCTTAAGCTTCAACGACAAAGAAACAATG |
| Core Oligo48 | GGAAGACAGGAGTCACAAACGGTCAGTG |
| Core Oligo49 | ACAAACTAATAAGTGCTCCAAAAGTACAA |
| Core Oligo50 | TACTCAGGGGATAGCAATGAATTT |
| Core Oligo51 | AAAACAGGTAACGTCAAGCG |
| Core Oligo52 | GCAAGGCCGTCAGACTAACCGCCTCAGC |
| Core Oligo53 | TAATAAGTTATTCTGAAGCCGCCGCTAA |
| Core Oligo54 | TCTGAATTGGGAGGGAAGGTAAATTCACCAATATACATAC |
| Core Oligo55 | AATTTCCAGAGCATTA |
| Core Oligo56 | AAATCAACAACCCGGAGTGA |
| Core Oligo57 | GAACCTATTTTAACGGAAATAAAT |
| Core Oligo58 | CACCACAAGAGAAGGGTTGAT |
| Core Oligo59 | TTTGTCATGTAGGAGCAGTA |
| Core Oligo60 | TCTTGAGAATCGCAACGCCAAAAATTCCGTAGCCAACATGTTCA |

| | |
|---|---|
| Core Oligo61 | GCGTAAAAAGGCCGTCGAGAGGATTAGG |
| Core Oligo62 | TTGATACCGTTCCAGTACCGATCATTAAGCGTCATA |
| Core Oligo63 | TTGAATTGCGAAGTTAGTAAAGCCCAATAGGAACC |
| Core Oligo64 | TCTGAGTTTCGTCACCAGT |
| Core Oligo65 | AAACCCTCAGCAGAACCATGGAAAACAGGAGTGTACTGG |
| Core Oligo66 | GTTGATAGAAGGACAAGCAAGCCGTTTT |
| Core Oligo67 | CAGAACCACCAGAACATGAATATCACCG |
| Core Oligo68 | ACTTTTTCACGACAATACATGTAATAATTTACAAACCAAT |
| Core Oligo69 | TACACTCCTCCCGACTTGCGGGGAGGCGTTTTAGCGAATACACT |
| Core Oligo70 | TACACTTCAGAGGGTAATTAGATAGCCGAACAAAGTTATACACT |
| Core Oligo71 | TACACTATTCTTACCAGTAACAGCCATTACACT |
| Core Oligo72 | TACACTATAAGTCCTGATCATTCCAAGATACACT |
| Core Oligo73 | TACACTCTAAATTTATGCGTTATACAATACACT |
| Core Oligo74 | TACACTACGGGTATTAAACCAAGTACCGCACTCCGGC |
| Core Oligo75 | TACACTTAAAGTACCGACAAAACAGTAATAAGAGAATATACACT |
| Core Oligo76 | AGTGCCTGTTTATCAACAATAGTACACT |
| Core Oligo77 | TACACTATTATTTAACCCTGAACAAAGTACACT |
| Core Oligo78 | AGTATCATAATGGTTTAATTTCATCTTCTGACTACACT |
| Core Oligo79 | TACACTCCAGAAGGAAACCGAGGAAACGCAATAAAGT |
| Core Oligo80 | TACACTGGAATTAGAGCCAGCAAAACGGTCATAATCA |
| Core Oligo81 | CACCGACTTGAGCCATTTGTACACT |
| Core Oligo82 | TACACTAGTTTCAGATCGCCCACGCTACACT |
| Core Oligo83 | AGCCGGTGAATTAGCACCATAAACGCAAAGACACCACGGTACACT |
| Core Oligo84 | GGCCCCCCTGCCCCTCAGAACCGCCACCCTCATACACT |
| Core Oligo85 | TACACTAATAAGTTTATTTTGTCAC |
| Core Oligo86 | TACACTTTGCCATCTTTTCATAGCCCCCTTATTAGCGTTACACT |
| Core Oligo87 | TACACTACGAAAGAGGCAAAAGAAGGCACCAACCTAAATACACT |
| Core Oligo88 | TACACTTAAACAGTTAATGAGGTCAGATACACT |
| Core Oligo89 | TACACTGAACCGCCACCCTCTTTCAACTACACT |
| Core Oligo90 | CCTTGAGTAACAGTGCCCGTATACACT |
| Core Oligo91 | TACACTATAACCGATATATTCGGTCCCACTACGAATACACTA |
| Core Oligo92 | TACACTCGATTGGCCTTGATATGTTGA |
| Core Oligo93 | TACACTCGATTGGCCTTGATATGTTGA |

| | |
|---|---|
| SLB Oligo1 | AAAGAAAAAGGGGAAACATGAGGTGCGGGATCTT |
| SLB Oligo2 | AAAGAAAAAGGGGAAATTTTTGTTGAAGCGCACCCAC |
| SLB Oligo3 | AAAGAAAAAGGGGAAACTTTAGCGGAAACGATTGACGATATGGT |
| SLB Oligo4 | AAAGAAAAAGGGGAAAATAAAGGTGGCAACAGAA |
| SLB Oligo5 | AAAGAAAAAGGGGAAAAAAGAACTGGCATGATATAG |
| SLB Oligo6 | AAAGAAAAAGGGGAAATGATAAATTAATTACTGCTT |
| SLB Oligo7 | AAAGAAAAAGGGGAAAGCAGTATGTTAGCAAACAAAAGGGCCGA |
| SLB Oligo8 | AAAGAAAAAGGGGAAAGCTACAGAAGCATCGGATCGG |

| | |
|---|---|
| Capture 1_3' | AGACTAATAATATAGGTGAGTATTA ATGTGT CCACAAGTCAGCGAG |
| Capture 1_5' | ACACAT AGAGGCTGTTGATGATGCGCAGTC |

7. **5’ DBCO Sequences for mAb functionalization:**

**Seq1:** 5-DBCO -ACATGACACTACTCCAC

**Seq1-complement:** GTGGAGTAGTGTCAT

**Seq2:** 5-DBCO - GTTCTCGCTGACTTGTGG

**Seq2-Complement:** CCACAAGTCAGCGAGA

**Seq3:** 5-DBCO - AAGTCAGCGAGAACCAG

**Seq3-Complement:** CTGGTTCTCGCTGAC

**Seq4:** 5-DBCO - CAACAGAGATAGAACAG

**Seq4-Complement:** CTGTTCTATCTCTGT

**Seq5:** 5-DBCO - TTAAAAGGGACATTCTG

**Seq5-Complement:** CAGAATGTCCCTTTT

**Note:** the sequence complement is the capture-strand sequence introduced to the capture_3′ oligo (as in the listed origami sequence examples), allowing hybridisation to the complementary DBCO oligo conjugated to the mAb. The capture strands are designed such that, upon hybridisation, two nucleotides at the 5′ end of the DBCO oligo remain unpaired.

8. **Cholesterol Oligo for SLB anchoring:**

5'- TTTCCCCTTTTTCTTTCC - 3’ cholesterol

9. **Strand Displacement sequences:**

- **Capture sequence:**

**1496-structure, capture 3_3’SD:**
CGGAGATTTTATTCATTACCACGGAAACG ATGTGT GTGGAGTAGTGTCAT GTA CTG TTC TAT CTC TGT

- **5’ DBCO oligo used:**

**Seq1:** 5-DBCO -ACATGACACTACTCCAC

- **SD sequence:**

ACAGAGATAGAACAGTACATGACACTACTCCAC

### V. Movies:

**Movie S1:**
5 s of a representative median-subtracted movie of a dynamic MP measurement of Trf-DNs tethered to a SLB composed of DOPE-PEG2k and DOPC (1.4:98.6%). The field of view is 10.8 × 10.8 μm², and the contrast scale is from −0.03 to +0.03. The raw frames were saved at 360 Hz, with a median window size of 600 frames. The movie shows 1800 frames between frames 1200 and 3000, with playback at 30 Hz. Scale bar is 1 μm.

**Movie S2:**
5 s of a representative median-subtracted movie of a dynamic MP measurement of Trf-DNs tethered to a SLB composed of DOPE-PEG2k and DOPC (1.4:98.6%), following the addition of 400 pM recombinant Trf protein and incubation for 5 min. The field of view is 10.8 × 10.8 μm², and the contrast scale is from −0.03 to +0.03. The raw frames were saved at 360 Hz, with a median window size of 600 frames. The movie shows 1800 frames between frames 12000 and 13800, with playback at 30 Hz. Scale bar is 1 μm.

**Movie S3:**
5 s of a representative median-subtracted movie of a dynamic MP measurement of p53-DNs tethered to a SLB composed of DOPE-PEG2k and DOPC (1.4:98.6%), following the addition of 40 nM recombinant p53 protein and incubation for 5 min. The field of view is 10.8 × 10.8 μm², and the contrast scale is from −0.03 to +0.03. The raw frames were saved at 360 Hz, with a median window size of 600 frames. The movie shows 1800 frames between frames 12000 and 13800, with playback at 30 Hz. Scale bar is 1 μm.

**Movie S4:**
5 s of a representative median-subtracted movie of a dynamic MP measurement of Dual EGF-DNs tethered to a SLB composed of DOPE-PEG2k and DOPC (1.4:98.6%), before the addition of EGF. The field of view is 10.8 × 10.8 μm², and the contrast scale is from −0.03 to +0.03. The raw frames were saved at 360 Hz, with a median window size of 600 frames. The movie shows 1800 frames between frames 12000 and 13800, with playback at 30 Hz. Scale bar is 1 μm.

**Movie S5:**
5 s of a representative median-subtracted movie of a dynamic MP measurement of Dual EGF-DNs tethered to a SLB composed of DOPE-PEG2k and DOPC (1.4:98.6%), following the addition of 2 nM recombinant EGF protein and incubation for 5 min. The field of view is 10.8 × 10.8 μm², and the contrast scale is from −0.03 to +0.03. The raw frames were saved at 360 Hz, with a median window size of 600 frames. The movie shows 1800 frames between frames 12000 and 13800, with playback at 30 Hz. Scale bar is 1 μm.

**Movie S6:**
5 s of a representative median-subtracted movie of a dynamic MP measurement of a four-structure, mass–diffusion-encoded DN panel tethered to a SLB composed of DOPE-PEG2k and DOPC (1.4:98.6%). The panel consists of four distinct DNA origami nanostructures with defined mass and diffusivity signatures, enabling simultaneous multiplexed detection within a single field of view. The field of view is 10.8 × 10.8 μm², and the contrast scale is from −0.03 to +0.03. Raw frames were acquired at 360 Hz, with a median background subtraction window of 600 frames. The movie shows 1800 frames between frames 12000 and 13800, played back at 30 Hz. Scale bar is 1 μm.

**Movie S7:**
5 s of a representative median-subtracted movie of a dynamic MP measurement of a four-structure, mass–diffusion-encoded DN panel tethered to a SLB composed of DOPE-PEG2k and DOPC (1.4:98.6%), recorded in 1% human plasma containing native Trf and spiked recombinant ErbB2 (0.5 nM), following a 5 min incubation. The field of view is 10.8 × 10.8 $\mu m^2$, and the contrast scale is from −0.03 to +0.03. Raw frames were acquired at 360 Hz, with a median background subtraction window of 600 frames. The movie shows 1800 frames between frames 12000 and 13800, played back at 30 Hz. Scale bar is 1 µm.
